# A Theoretical Framework for the Hemodynamic Role of Sarcomere Length Dynamics During the Isovolumic Phase of the Left Ventricle

**DOI:** 10.64898/2026.03.16.712012

**Authors:** Shiro Kato, Kota Kishida, Akira Amano, Yukiko Himeno, the Lorem Ipsum Consortium

## Abstract

The left ventricle (LV) exhibits torsional deformation during systole, and mechanical relaxation begins during the isovolumic phase. Recent advances in imaging techniques, such as MRI, have revealed that myocardial tissue deformation and sarcomere length changes occur during the isovolumic relaxation phase, when chamber volume remains constant. Although such ventricular deformation during the isovolumic phase is considered important for blood ejection and filling efficiency, its mechanistic contribution to contraction and relaxation remains unresolved.

In this study, we hypothesized that the sarcomere length dynamics during isovolumic phase affects the isovolumic contraction and relaxation time (IVCT and IVRT) by regulating the contraction force. To investigate this hypothesis, we focused on experimentally reported differences in the relationship between sarcomere length and LV volume across the endocardial and epicardial layers, as described by Rodriguez et al. We constructed and compared two types of hemodynamic models within the same integrated framework consisting of a circulation model, a LV model, and a myocardial cell contraction model by Negroni-Lascano et al., which differ only in how sarcomere length is determined: a volume-based length model (VL model), in which sarcomere length is uniquely determined by LV volume, and a volume–force–coupled length model (VFL model), in which sarcomere length is determined by the balance between LV volume and contraction force.

Simulation results showed that in the VFL model, compared to the VL model, sarcomere length changed during the isovolumic phase, promoting a decrease in contraction force and shortening of IVRT. These results indicate that sarcomere length dynamics can mechanically regulate force decay during isovolumic relaxation, even under constant left ventricular volume. This study provides a theoretical framework for understanding the contributions of different layers within the LV wall to diastolic function during the isovolumic relaxation phase.

**Author summary:** Same as Abstract

## Introduction

Diastolic relaxation is as crucial as systolic contraction for maintaining heart pump function. Diastolic dysfunction is more common than systolic dysfunction in congestive heart failure patients [1], and particularly common in the elderly [2]. Isovolumic Relaxation Time (IVRT) is considered an important index for assessing diastolic dysfunction, suggesting its importance for physiological blood circulation. Historically, the key parameters determining isovolumic relaxation indices, including force decline rate, pressure decline rate, and the relaxation time constant, were examined. From the report by Wiegner et al. [3], maximum isovolumic relaxation rate (max dF/dt) increases with increasing end-systolic muscle fiber length. From the report by Gaasch et al. [4], the isovolumic relaxation time constant (*τ* ) increases with increase in systolic pressure, and linearly relates to systolic fiber length. Similarly, from the report by Raff et al. [5], the time constant of exponential isovolumic pressure fall T increases with increase in the end diastolic pressure. However, from the report by Gaasch et al. [6], in the experiments on the single cardiac muscle fiber, the preload had no relation with the isovolumic relaxation time constant.

A different type of analysis was performed by Wiegner et al. [7], where the simple mathematical model of cardiac relaxation was constructed and used to analyze the parameters which determine the relaxation characteristics. However, in this model, force was represented by a simple exponential function whose amplitude and time constant were uniquely determined by the stimulation conditions. As a consequence, fiber length during the isovolumic phase was assumed to remain constant, and the effect of length change was not considered.

In subsequent studies, improvements in image processing techniques allowed the analysis of changes of fiber length during the isovolumic relaxation phase, and the effect of the left ventricular (LV) untwisting is considered to be an important contributor to the pressure drop during isovolumic relaxation phase. In this context, the release of the elastic potential energy accumulated in the passive elastic element during systolic deformation is considered to be an important contributor to the isovolumic relaxation. Recent imaging techniques have revealed that the ventricular tissue have deformation during isovolumic relaxation phase and also the fiber length is simultaneously changing.

Many reports have been published on untwisting in the isovolumic phase using image measurements. According to MRI measurements of dogs by Rademakers et al., [8], it was confirmed that untwisting occurred even in the isovolumic phase, and that the circumferential segmental length when viewed in cross section also changed. Furthermore, with dobutamine infusion, torsion at end systole and reversal relaxation were greater than in controls. In more detailed reports, left ventricular untwisting rate (UTR) is measured using speckle-tracking echocardiography [9] and tagging MRI [10]. In the report by Notomi et al. [11], UTR is reported to correlate with the isovolumic relaxation rate. Moreover, approximately 40% of the untwisting occurs during the isovolumic relaxation phase.

Rodriguez et al. reported the results of measuring myocardial tissue strain on the epi-, mid-, and endocardium throughout the cardiac cycle [12]. In this experiment, beads were implanted in multiple locations in a dog’s heart, and tissue deformation was estimated by taking X-ray videos from two directions. The LV volume was also measured simultaneously, allowing the evaluation of correlation between the changes in tissue length relative to LV volume change for the epi-, mid-, and endocardium. Interestingly, while tissue length and LV volume were roughly proportional in the endocardium, the relationship between LV volume and tissue length on the epicardium changed between systole and diastole, and furthermore, tissue length changed rapidly during the isovolumic relaxation phase.

Apart from these organ- and tissue-level studies, the properties of ventricular myocyte contractility, such as the force-velocity relationship [13, 14] and the force drop due to instantaneous shortening [15], have long been known. If untwisting during the isovolumic relaxation phase involves the elongation of myocardial cells, it is possible that the force decrease due to the force-velocity relationship has a greater impact on isovolumic relaxation than the potential energy release of the elastic element.

From these considerations, the length change during isovolumic relaxation may contribute to improved hemodynamic efficiency. However, because sarcomere length and LV volume are not uniquely correlated in the epicardium but exhibit hysteresis, their impact on hemodynamics may not be uniformly beneficial. In this study, we aimed to elucidate the effect of sarcomere length change during isovolumic phase by constructing a hypothetical relation between sarcomere length and LV volume affected by cellular force. Since the geometrical detail of the LV volume, sarcomere length and cellular force in epi- and endocardium are not understood, we hypothesized that sarcomere length changes in the epicardium during the isovolumic phase are caused by shape changes due to changes in contractile force, and constructed a left ventricular volume, contractile force, sarcomere length relation model that reproduces the measurement data of Rodriguez et al. We therefore constructed integrated hemodynamic models in which sarcomere length is either uniquely determined by LV volume or dynamically determined by the balance between LV volume and contraction force. By comparing these models, we aimed to clarify whether sarcomere length dynamics during the isovolumic phase can mechanically regulate the time course of force and IVRT under constant ventricular volume.

## Simulation Model and Computational Method

### Model structure

To analyze the relation between sarcomere length changes and hemodynamics during isovolumic phase, two hemodynamic models were used in this research: the volume-based length model (VL) and the volume-force-coupled length model (VFL).

In many previous researches which aims at analyzing the relation between the hemodynamics and the characteristics of the ventricular cells, the relation between the LV volume and the ventricular cell length was typically based on the hemispheric geometry relationship. Here, We refer to this volume-length relationship as the VL model. The VL model is constructed from the equation where the half sarcomere length (*L*) is uniquely determined by the LV volume (*V*_*lv*_). In this model, *L* remains constant during the isovolumic phase.

In contrast, in the VFL model, *L* of ventricular cell is not determined by only *V*_*lv*_ but determined by the balance between *V*_*lv*_ together with the ventricular cell contraction force (*F* ). As a result, *L* can dynamically change even during the isovolumic phase.

Both models are constructed by integrating a circulation model, a Left Ventricular model, and a ventricular cell contraction model, based on the hemodynamic model which was proposed by our group [16–18].

### Circulation model

The simplified windkessel based circulation model was used in this study (Fig. 1). This model is identical to that used in our previous research [18].

**Fig 1.**
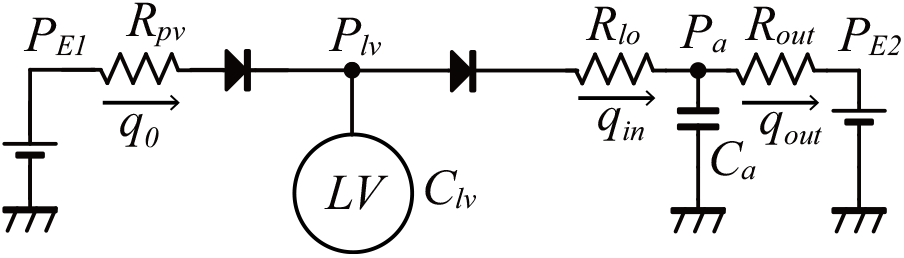
Circulation model. Reproduced from [18], under the terms of the Creative Common Attribution License.

The parameters applied in this research are explained in Table 1, and are primarily derived from the circulation models proposed by Heldt et al. [19] and Liang et al. [20]. We used constant values for pulmonary venous pressure (*P*_*E*1_) and peripheral pressure (*P*_*E*2_), and moreover the baroreflex was excluded from the model as the aim was to reproduce the baseline hemodynamics. For the aortic compliance(*C*_*a*_), it has a relation with the aortic pressure and volume as follows.

**Table 1.**
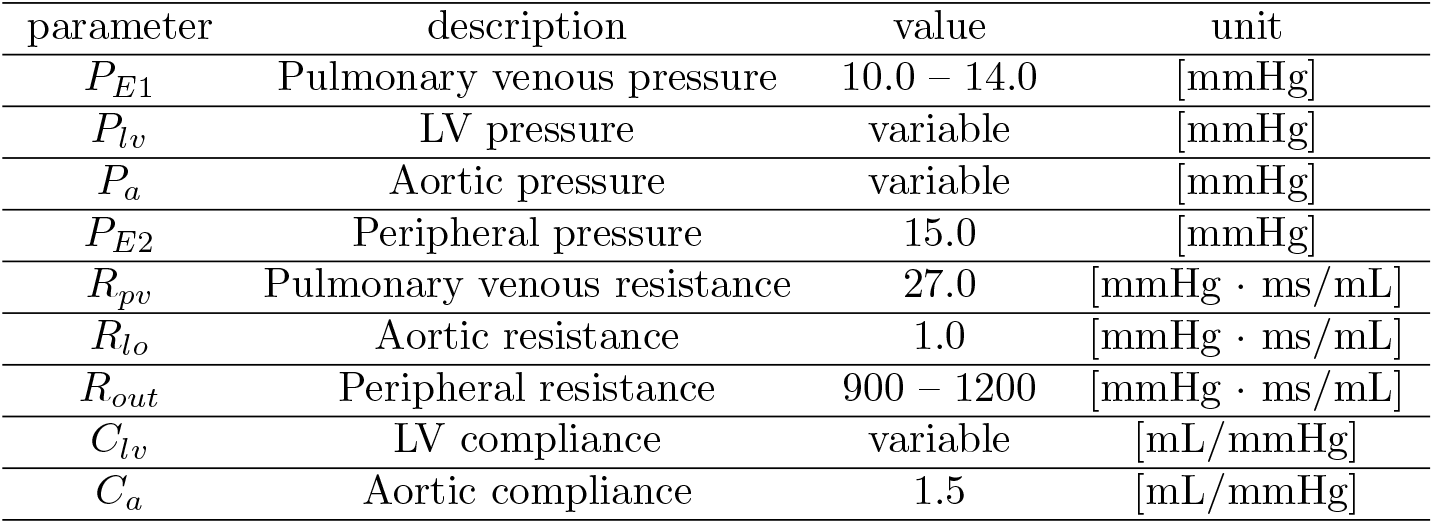
Descriptions of the circulation model variable.

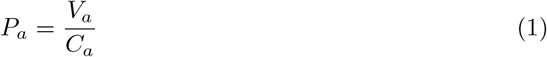

Additionally, the cardiac cycle length was fixed at 1000 [ms], and the several values for peripheral resistance i.e. afterload (*R*_*out*_) and pulmonary venous pressure i.e. preload (*P*_*E*1_) were used to analyze the effect of afterload and preload variation.

### Ventricular cell contraction force model

The muscle cell contraction force model proposed by Negroni and Lascano (NL08) is a well-established model capable of reproducing key features of myocardial mechanics, including isometric contraction, isotonic shortening, and force-velocity relation (FVR). In this study, we applied the model in the hemodynamic model for reproducing contraction force. A complete description of NL08, including the full set of state-transition equations and parameter definitions, is available in the original publication [21]. Here we summarize only the components directly related to this study.

The active force generation element of NL08 model consists of two components: (i) a chemical component that reproduces the calcium transition state within troponin system (See Supplementary Material S2 ModelEquation Fig.1), and (ii) a mechanical component that represents the crossbridge dynamics (Fig. 2). There are two types of the states in the forming crossbridge: the weakly bound state (∼) and the strongly bound state (∗ ). The contraction force (*F*_*b*_) shown in Eq.(2) represents the sum of the forces generated by the crossbridges in these two states, multiplied by the crossbridge lengths and the concentrations of troponin system binding the crossbridges in each state. *A*_*w*_ and *A*_*p*_ are constants that are the stiffness of the crossbridges in the weakly and the strongly bound state, respectively.

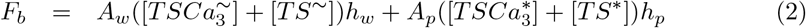

**Fig 2.**
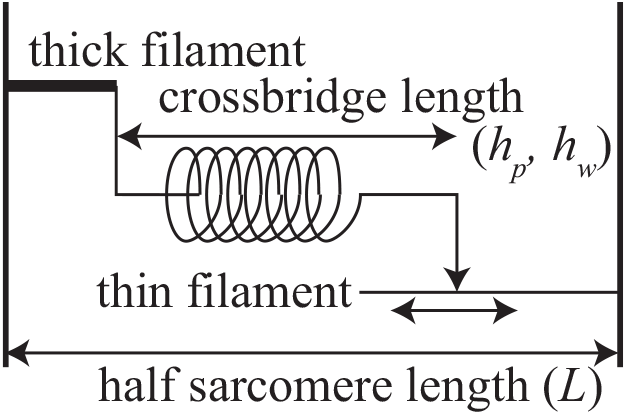
NL08 mechanical model. Reproduced from [18], under the terms of the Creative Common Attribution License. The crossbridge lengths in the weakly bound state and the strongly bound state are represented by *h*_*w*_ and *h*_*p*_, respectively.

The passive force *F*_*p*_ of NL08 model is shown in Eq.(3) and the formulation represents the force generated by the passive elastic elements within the sarcomere. *K*_*e*_ and *L*_*e*_ denote the stiffness of the nonlinear and linear elements, and *L*_0_ is the reference length at which passive force becomes zero.

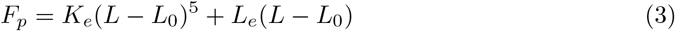

Experimental measurements of passive and active force are obtained from different preparations: passive force is typically measured in tissue samples [22, 23], whereas active force is often measured either in single isolated cells or in tissue preparations. Therefore, the absolute magnitudes of *F*_*p*_ and *F*_*b*_ in the NL08 model may not correspond to the cellular force of the LV wall. The scaling coefficients *K*_*ps*_ and *K*_*bs*_ were introduced for a single cellular force (*F* ) to adjust the magnitudes of *F*_*p*_ and *F*_*b*_, respectively. The values of *K*_*ps*_ and *K*_*bs*_ were manually adjusted to reproduce the physiological human hemodynamics in the section of simulation conditions, which results in *K*_*bs*_ = 2.0 and *K*_*ps*_ = 0.5. Finally, cellular force *F* [mN/mm^2^] was calculated as follows.

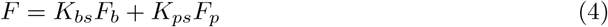

The crossbridge length in the weakly bound state (*h*_*w*_) is defined as the difference between the sarcomere length (*L*) and the non-elastic portion (*X*_*w*_) as shown in Eq.(5). The time derivative of *X*_*w*_ is given in Eq.(6), where *B* controls the rate of change of *X*_*w*_, and *h*_*wr*_ is the reference length of *h*_*w*_.

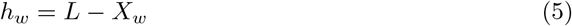

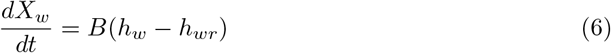

In the NL08 model, the troponin transition state is influenced by force-length relation (FLR) and force-velocity relation (FVR). FLR is represented by the actin-myosin overlap function in the original formulation. On the other hand, the FVR, which describes the decrease in force with sarcomere shortening or lengthening velocity, is controled by *g* and *g*_*d*_, which determines the crossbridge detachment velocity. Both *g* and *g*_*d*_ depend on the crossbridge extension, and these values increase with (*h*_*w*_ − *h*_*wr*_)^2^. In this study, we adopted the modified equations for *g* (Eq.(7)) and *g*_*d*_ (Eq.(8)) proposed by Utaki et al. [17] to improve the hemodynamic reproducibility when the NL08 model is incorporated into the circulation model. They introduced a parameter *γ*_*m*_ (Eq.(9)) that reduces the detachment rate during sarcomere lengthening 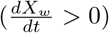.

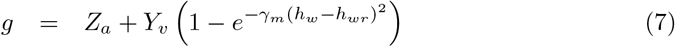

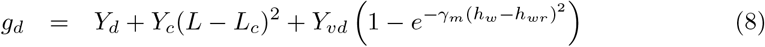

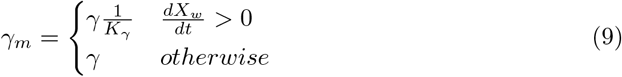

### Left Ventricular pressure and the cell contraction force relation model

Our aim was to analyze how sarcomere length dynamics during isovolumic phase influences the hemodynamics. To have the simulation model that achieves the aim, Left Ventricular model is necessary to link the molecular contraction model to the circulation model. Thus, the model relates LV pressure and volume (calculated from the circulation model) to sarcomere length and contraction force. Based on Laplace’s law, we used the equations to derive the LV internal radius (*R*_*lv*_), the wall thickness (*h*_*lv*_) and sarcomere length (*L*) as explained below. The constants used in this section are summarized in Table 2.

**Table 2.**
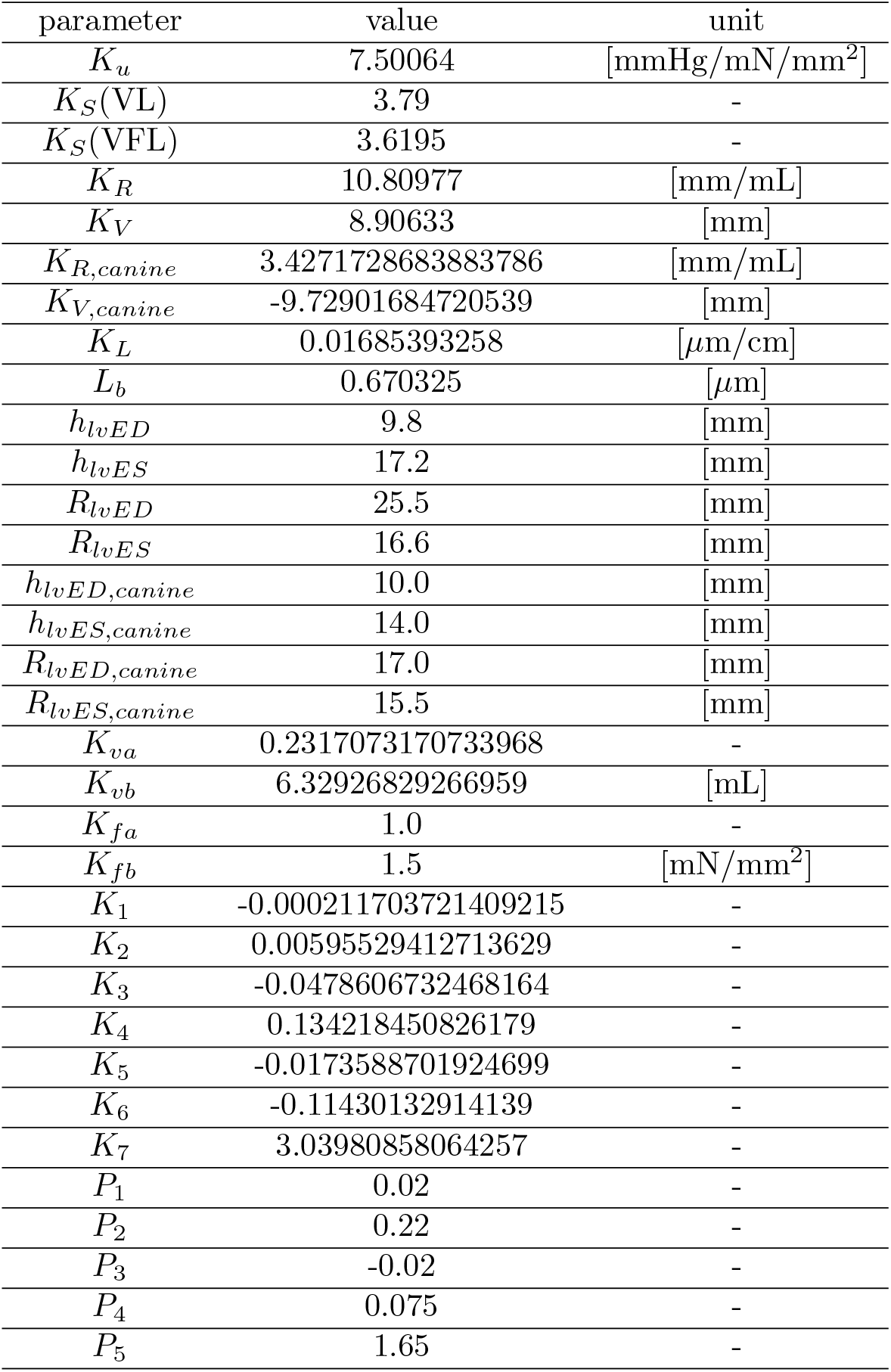
Parameters used for Left Ventricular model.

Laplace’s law is well-known to represent the relation between LV pressure (*P*_*lv*_), wall thickness (*h*_*lv*_), wall tension (*F*_*ext*_) and internal radius (*R*_*lv*_) [24]. To calculate *P*_*lv*_ from *F*_*ext*_, *h*_*lv*_ and *R*_*lv*_ in our model, we used the following equation.

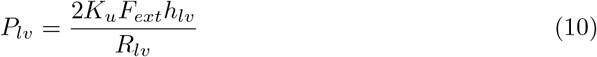

Here, *K*_*u*_ is constant that converts units from [mN/mm^2^] to [mmHg]. The derivations of *F*_*ext*_, *R*_*lv*_, *h*_*lv*_, and *L* are explained in the following.

As explained in the previous section, the contraction force of the cell (*F* ) is derived from the ventricular cell contraction model. Since this value is at the cellular level, we need to convert it into the LV wall tension in order to apply Laplace’s law. Thus, we assumed LV wall tension to be proportional to the cellular contraction force and introduced a scaling parameter (*K*_*S*_) to convert the cellular contraction force (*F* ) into the LV wall tension (*F*_*ext*_). The values of *K*_*S*_ were manually adjusted so that the ejection fraction (EF) remained at 60.0% at the periodic limit cycle under each simulation condition that are explained in the section of Simulation conditions.

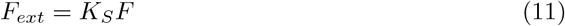

Since Laplace’s law assumes that the physical shape of LV is spherical, we took this assumption into account and used the following equation that incorporates the relation between the volume of a sphere and its internal radius.

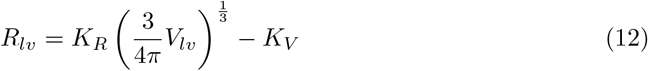

Here, *K*_*R*_ and *K*_*V*_ are constant and determined by solving a system of simultaneous equations, in which the values of *V*_*lv*_ and *R*_*lv*_ at end-diastole and end-systole were substituted into Eq. (12). The values of *V*_*lv*_ were extracted from the data used by Utaki et al. [17], and the values of *R*_*lv*_ were taken from data reported by Sutton et al. [25].

LV wall thickness is maximal at end-systole and minimal at end-diastole, and its time course during the cardiac cycle has been extensively studied. Yun et al. [26] reported that the wall thickness is related to LV twist angle and volume. Additionally, Sutton et al. [25] showed that wall thickness varies depending on the position of the LV and is not always proportional to LV volume. Althogh the mechanism underlying these changes has been partially revealed, the details remain unresolved. In this study, we assumed that wall thickness(*h*_*lv*_) is proportional to the internal radius(*R*_*lv*_).

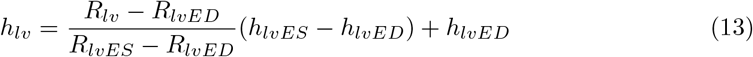

Note that *h*_*lvED*_ and *h*_*lvES*_ represent the LV wall thickness at end-diastole and end-systole, respectively. Gerstenblith et al. [27] measured the LV wall thickness in human adults using echocardiographic methods and reported the average values for each age group. We extracted the median values of end-diastole and end-systole from their data and used for *h*_*lvED*_ and *h*_*lvES*_. Here, *R*_*lvED*_ and *R*_*lvES*_ represent LV internal radius at end-diastole and end-systole, respectively. We used those values from the data reported by Sutton et al. [25].

Next, in this study, we focused on the differences of sarcomere length change across the LV wall, which is reported by Rodriguez et al. [12]. We constructed two types representing layer-dependent sarcomere length behavior : a volume-dependent length (VL) model corresponding to the endocardium, and a volume-force-coupled length (VFL) model corresponding to the epicardium. These models were used to analyze how the sarcomere length change during isovolumic phase affects the hemodynamics.

- Sarcomere length of volume-dependent length model (VL) The VL model represents the endocardial behavior, where the relation between sarcomere length and LV volume is approximately linear. Utaki et al. [17], Taniguchi et al. [16] and Kato et al. [18] employed the following formulation, and their papers explained the details of the derivation. This equation assumes that the relation between *L* and LV internal radius(*R*_*lv*_) is linear, with *R*_*lv*_ uniquely determined by *V*_*lv*_ as shown in Eq. (12). Thus, *L* becomes constant during isovolumic phase.

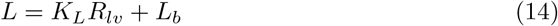
- Sarcomere length of volume-force-coupled length model (VFL) The VFL model represents epicardial behavior, where sarcomere length exhibits hysteresis with respect to LV volume. We introduced a new equation describing changes in sarcomere length through the following process. Rodriguez et al. [12] measured the time course of LV pressure, volume and sarcomere length during cardiac cycles in the hearts of canines. Subbah et al. [28] reported the time course of LV internal radius and wall thickness. We combined their data and calculated the contraction force(*F*_*ext*_) by using the Eqs. (10), (12) and (13). In addition, we obtained the relation between LV volume, contraction force and sarcomere length in the canines’ hearts as shown in Fig.3. By applying polynomial approximation to this relation, we derived the following equation which determines *L* as a function of *V*_*lv*_ and *F*_*ext*_ using fitted parameters *P*_1_ to *P*_5_ and polynomial terms *h*_1_ to *h*_7_.

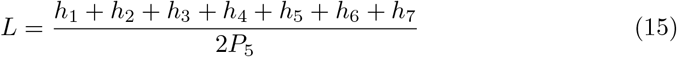

**Fig 3.**
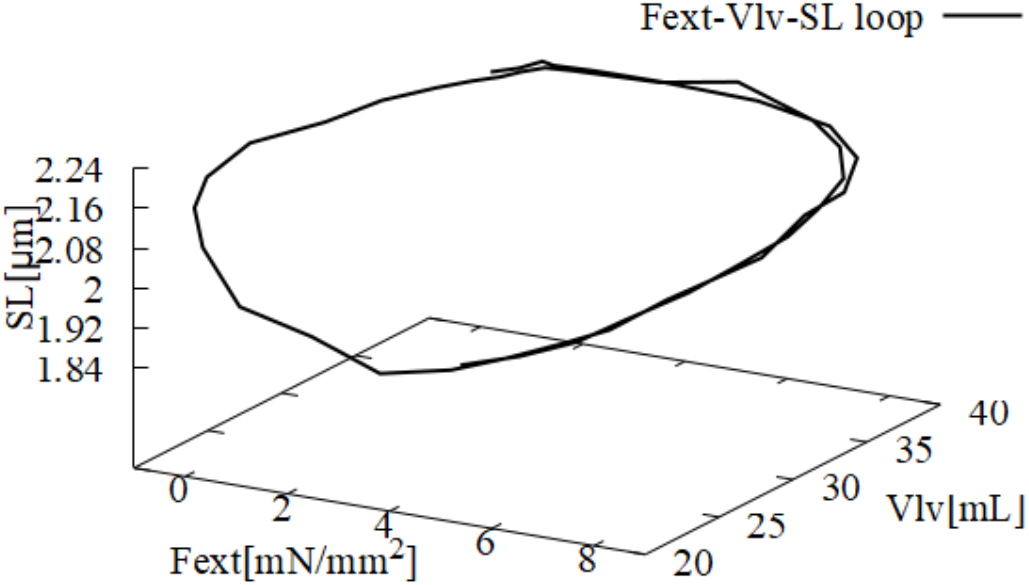
*F*_*ext*_-*V*_*lv*_-*SL* loop during cardiac cycle on 3D space. The sarcomere length (*SL*) and the LV volume (*V*_*lv*_) are ploted from data Rodriguez et al. reported in [12]. *F*_*ext*_ is calculated by combining wall thickness and internal radius Subbah et al [28] reported.

Here, the terms for *h*_1_ to *h*_7_ are shown in Supplementary Materials ( S2 ModelEquation Eqs.(16)–(22) ). The following are the steps of the derivation of this equation.

a. We converted Eqs. (12) and (13) into Eqs. (16) and (17), scaling them for the canine’s heart using the LV volume from Rodriguez et al. [12], and the LV internal radius and wall thickness from Sabbah et al. [28]. We then obtained the time course of the canine’s LV internal radius (*R*_*lv,canine*_) and wall thickness (*h*_*lv,canine*_) by substituting LV volume(*V*_*lv,canine*_) from Rodriguez et al. [12] into these equations.

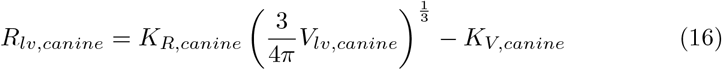

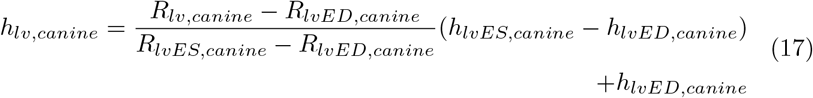
b. We derived contraction force (*F*_*ext,canine*_) by substituting the LV pressure in Rodriguez et al. [12] and *R*_*lv,canine*_, *h*_*lv,canine*_ derived in above a) into Laplace’s law (Eq.(10)).
c. Here, we deived x-axis (Eq.(18)) and y-axis (Eq.(19)) by manually adjusting *P*_1_–*P*_5_ so that the sarcomere length-LV volume-LV wall tension(*SL*-*V*_*lv*_-*F*_*ext*_) loop in Fig.3 forms a curved line as shown in Fig.4. The values of *P*_1_–*P*_5_ are shown in Table 2. Note that muscle cell contraction model use half of sarcomere length (*L*). Thus, we divide *SL* by 2 to convert *SL* to *L*.

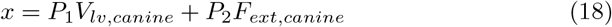

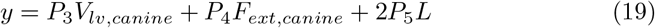
d. By applying a sixth-order polynomial approximation to the plotted curve by Eqs. (18) and (19), we obtained the following approxmation equation.

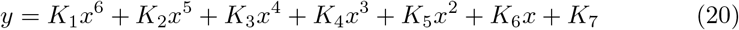
e. We solved for *L* by combining Eqs.(18), (19) and (20), and obtained Eq.(15). Eq.(15) is derived from measured canine’s LV data. Thus, the convertion from human-scaled to canine-scaled needs to incorporate with the equation. Assumed that the half sarcomere length of LV is identical between human and canine, we derived the convertion equations of LV volume and contraction force from human to canine (Eqs.(21) and (22)).

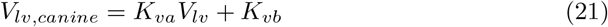

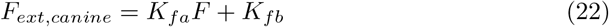

The parameters *K*_*va*_ and *K*_*vb*_ were determined by solving the simultaneous equation for Eq.(21), with the values of reported data for human LV volume (*V*_*lv*_) and canine LV volume (*V*_*lv,canine*_) at end-diastole and end-systole.

Incorporation of Eq.(15) into hemodynamic model with NL08 model revealed the computational instability. We observed that trajectories passing through certain regions of the *F*_*ext*_-*V*_*lv*_-*SL* space defined by Eq. (15) led to oscillatory or numerically unstable solutions, reflecting strong nonlinear feedback between force generation and sarcomere dynamics. To ensure that trajectories remain within physiologically relevant, non-oscillatory regions of this space, the scaling parameters *K*_*fa*_ and *K*_*fb*_ were constrained to ranges that prevent traversal of these unstable regions. As a consequence of this restriction, simulated trajectories do not necessarily coincide with the experimentally measured data from canine LV by Rodriguez et al. Moreover, these constraints affect the conversion from human-scaled LV wall tension *F*_*ext*_ to canine-scaled’s tension *F*_*ext,canine*_ so that the resulting trajectory in the *F*_*ext*_–*V*_*lv*_–*L* space avoids unstable regions.

The model parameter values we used in Left Ventricular model are summarized in Table 2.

**Fig 4.**
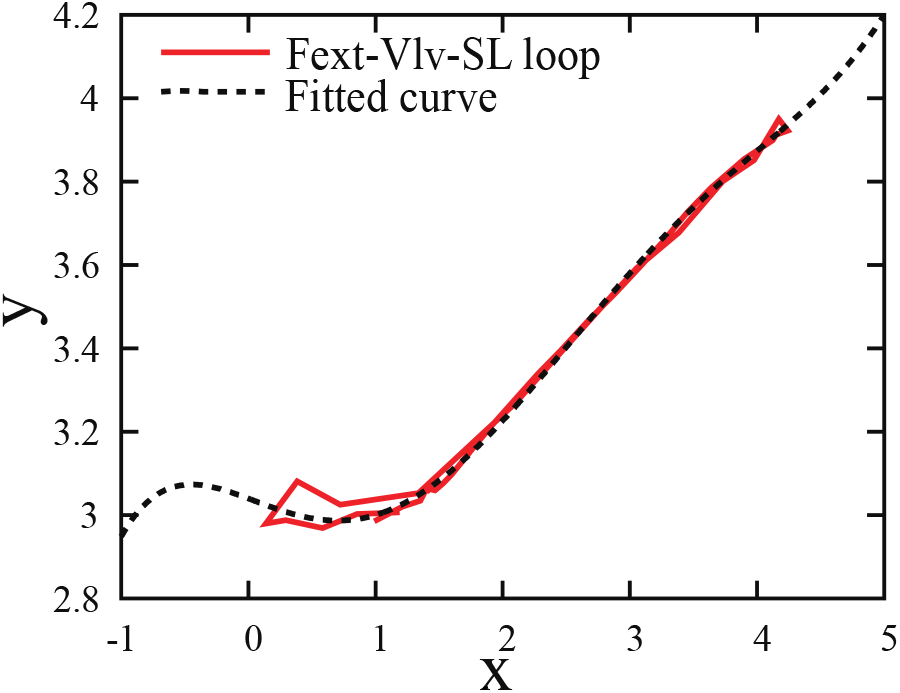
*F* -*V* -*SL* loop and fitted curvexon 2D(x-y) plain. The x-y plain is determined from the specific angle on *F*_*ext*_-*V*_*lv*_-*SL*.

### Computational method

In terms of the computational method, the forward Euler method was applied to simulate the state of muscle cell contraction model’s troponin transition, and the nested bisection method was used to solve the coupled relation between LV volume (*V*_*lv*_), the contraction force (*F*_*ext*_) and the half sarcomere length (*L*). This method enables to search the values of *V*_*lv*_, *L*, and *F*_*ext*_ that balance with the toroponin transition. First, the initial bisection method is used to find the solution that converges to the given *V*_*lv*_ at an arbitrary time step. Then, the nested bisection method is used to search *L* that balances with *F*_*ext*_ and *V*_*lv*_ which is given by the initial bisection method. Through these approach, it is possible to simultaneously find the state where *V*_*lv*_, *F*_*ext*_ and *L* are balanced based on the troponin state. Additionally, this approach contributes to ensuring the stability and accuracy of the simulation. The detailed computational steps are as follows, with the calculation process illustrated in Fig 5.

**Fig 5.**
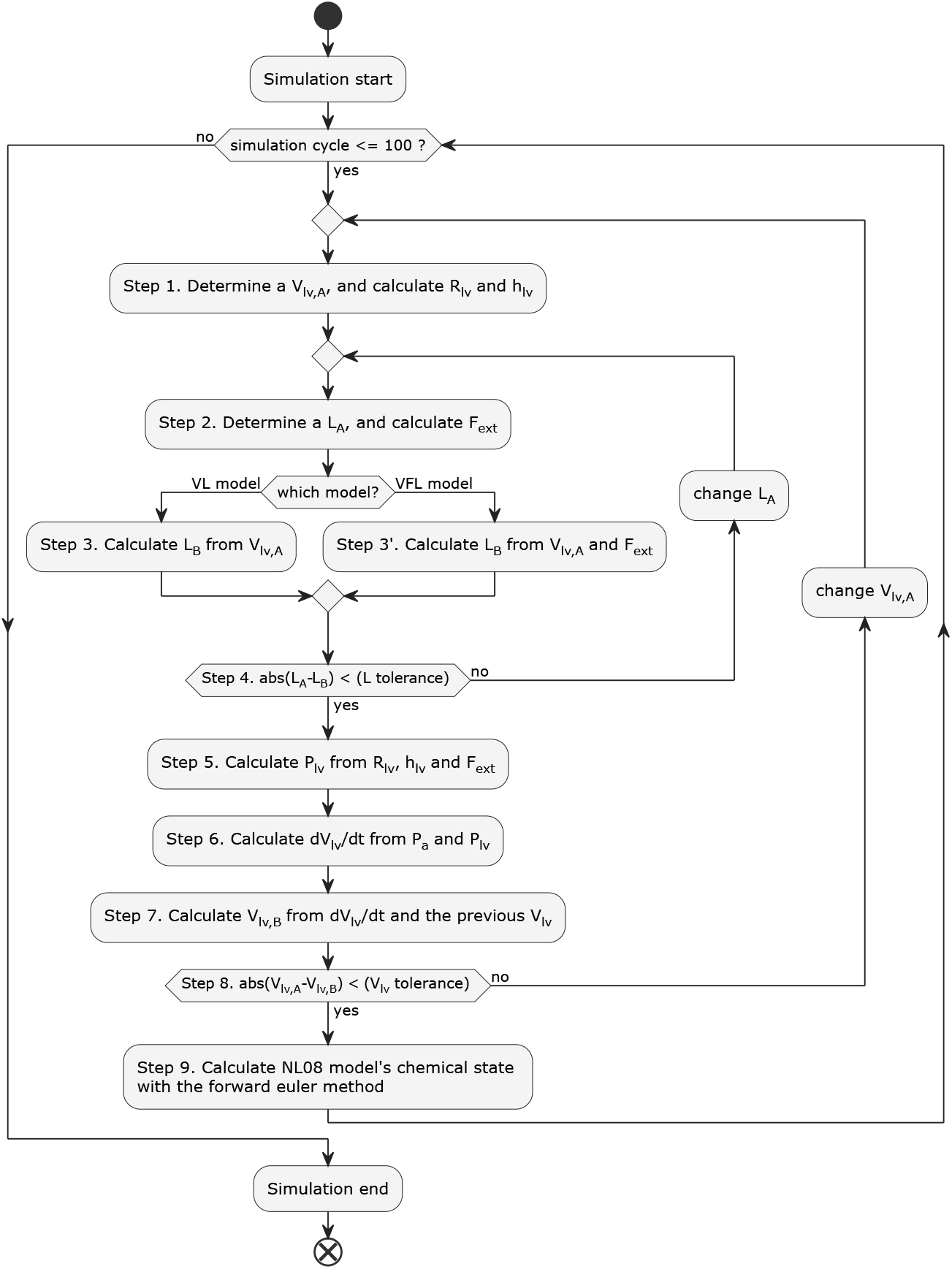
Computational process.

Step 1. Determine an arbitrary *V*_*lv,A*_ and calculate *R*_*lv*_ and *h*_*lv*_.

Step 2. Determine an arbitrary *L*_*A*_ and calculate *F*_*ext,A*_ using the troponin state.

Step 3. In the case of the VL model, calculate *L*_*B*_ from *V*_*lv,A*_. In the case of the VFL model, calculate *L*_*B*_ from both *V*_*lv,A*_ and *F*_*ext,A*_.

Step 4. Calculate the difference between *L*_*A*_ and *L*_*B*_, and return to Step 2 if the absolute difference does not meet the tolerance, with adjustment in *L*_*A*_.

Step 5. Calculate *P*_*lv*_ from *R*_*lv*_, *h*_*lv*_, and *F*_*ext*_ using Laplace’s law.

Step 6. Calculate *dV*_*lv*_*/dt* from *P*_*a*_ and *P*_*lv*_ using the circulation model.

Step 7. Calculate *V*_*lv,B*_ using *dV*_*lv*_*/dt* and the previous step’s *V*_*lv*_ with the Euler method.

Step 8. Calculate the difference between *V*_*lv,A*_ and *V*_*lv,B*_, and return to Step 1 if the absolute difference does not meet the tolerance, with adjustment in *V*_*lv,A*_.

Step 9. Calculate the troponin transition state for the next step using the forward Euler method.

### Simulation conditions

The simulation program was implemented in C language, with a fixed cardiac cycle duration of 1000 [ms]. Calculations were performed 100 cycles to achieve a periodic limit cycle, using a time step of 0.01 [ms]. We conducted following three sets of simulations to compare the VL model and the VFL model.

1. **fixed preload and afterload:** To investigate differences in the *L*-*V*_*lv*_ relation between the two models under fixed preload and afterload.
2. **afterload variations:** To assess changes in systolic and diastolic function (e.g., *τ* and Tei-index) in response to different afterload conditions.
3. **preload variations:** To evaluate how changes in end-diastolic volume caused by different preload conditions affect the end-systolic pressure-volume realtion (ESPVR) and contractility.

To ensure comparability of the two models, the scale parameter(*K*_*S*_) was adjusted for the two models as shown in Table 1, and the ejection fraction (*EF* ) was fixed at 60.0 % with fixed preload (*P*_*E*1_ = 12) and afterload (*R*_*out*_=1050) for both models. In addition, to reproduce load conditions, *P*_*E*1_ was varied as 10, 11, 12, 13, 14 [mmHg] for preload variation, *R*_*out*_ was varied as 900, 950, 1000, 1050, 1100, 1150, 1200 **[mmHg**·**ms/mL]** for afterload variation.

### Evaluation metrics

For evaluation of the preload variations, we used ESPVR and its slope as a measure of contractility. In contrast, for the afterload variations, additionlly the time-based indicators were derived from isovolumic contraction and relaxation phases. This is because the afterload primarily affects the time courses of contraction and relaxation rather than pressure-volume relation. These indices were obtained from the simulation results and compared between twomodels (Table 3).

**Table 3.**
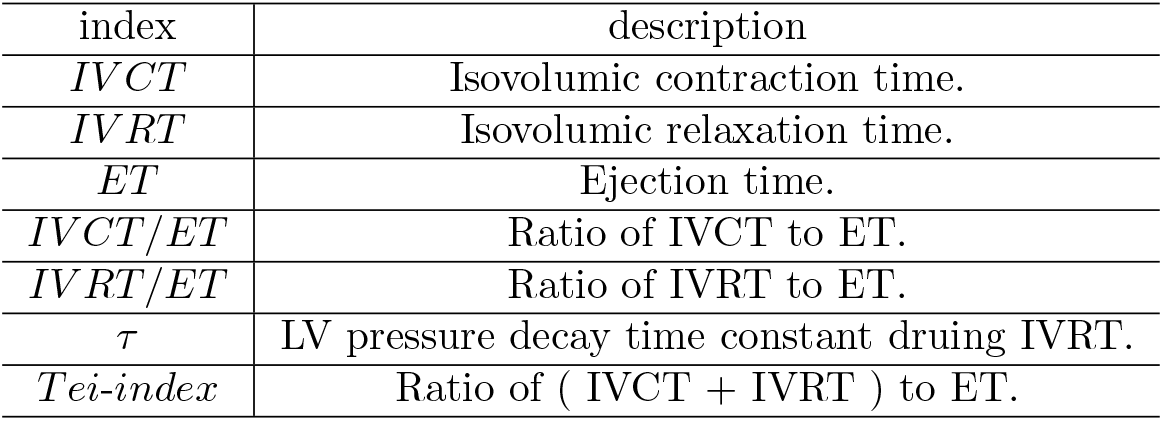
Summary of evaluation indices for afterload variation.

| index | description |
| --- | --- |
| $IVCT$ | Isovolumic contraction time. |
| $IVRT$ | Isovolumic relaxation time. |
| $ET$ | Ejection time. |
| $IVCT/ET$ | Ratio of IVCT to ET. |
| $IVRT/ET$ | Ratio of IVRT to ET. |
| $\tau$ | LV pressure decay time constant during IVRT. |
| $Tei-index$ | Ratio of ( $IVCT + IVRT$ ) to ET. |

For analysis of the isovolumic contraction and relaxation, we used IVCT/ET and IVRT/ET. IVCT and IVRT are considered to be different among each model, therefore these were standardized by each Ejection Time(ET), which makes each models independent of the heart rate. In addition, the time constant of LV relaxation (*τ* ) was calculated as follows ( Eq.(23) ).

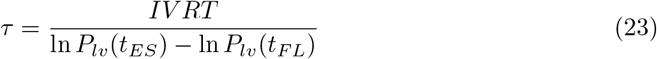

*P*_*lv*_(*t*_*ES*_) and *P*_*lv*_(*t*_*F L*_) represent the LV pressure at the end-systole and at the beginning of blood filling, respectively. These values are obtained from the final cycle of simulation results. In animal experiments, *τ* indicates the LV relaxation performance, and its value increases in case that LV relaxation capacity is reduced (Wang et al. [9]).

Tei-index was also employed to evaluate comprehensive systole and diastole function( Eq.(24)). Tei et al. originally proposed the left ventricular myocardial performance index (Tei-index) using echocardiographic measurements and reported a physiological value of approximately 0.35 – 0.45 in healthy subjects, with values exceeding 0.45 considered abnormal [29].

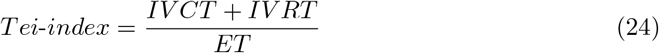

## Results

### Validation of the VFL formulation

In this section, we examine the properties of the introduced VFL model from two perspectives: (1) the time-course of sarcomere length (*SL*), and (2) the interaction among the three variables—LV volume (*V*_*lv*_), contraction force (*F* ), and sarcomere length (*SL*).

Firstly, we compare the time-course of *SL* generated by the VFL model with that reported by Rodriguez et al [12]. Fig.6 shows the time course of *SL* from the study, as well as the *SL* obtained by substituting the time course of LV volume and pressure into Eq.(15). We observe that the Eq.(15) can reproduce the overall temporal behavior of sarcomere length during a cardiac cycle.

**Fig 6.**
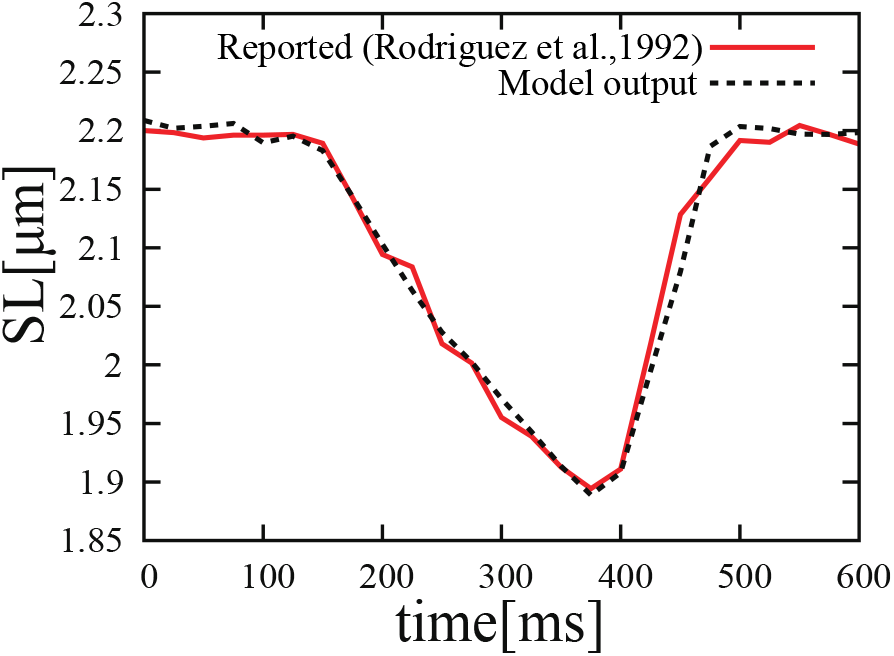
The time-course of *SL*. The red and black line represents the reported data (Rodriguez.et al. [12]) and Eq.(15) output using parameters from Rodriguez et al., respectively.

Next, we examine the relation between *V*_*lv*_ and *SL* when *F* is fixed at values of -0.5, 0.0, 1.0, 2.0, 3.0 and 4.0, as shown in Fig.7. The loop in Fig.3 shows the property that *SL* decreases with the reduction in *V*_*lv*_ and increases with the reduction in *F*_*ext*_. The *V*_*lv*_-*L* relation in Fig.7 can reproduce the property. At relatively higher contraction forces (*F* = 3.0, 4.0), the slope of *V*_*lv*_-*L* relation slightly increases, indicating that the LV volume becomes more sensitive to the sarcomere length variation.

### Model behavior under fixed preload and afterload

To investigate the relation between sarcomere length (*L*) and LV volume (*V*_*lv*_) under fixed ejection fraction (*EF* ), preload (*P*_*E*1_) and afterload (*R*_*out*_), we performed simulations using the both models under *P*_*E*1_ = 12 and *R*_*out*_=1050. The time courses of *L* and *V*_*lv*_ are shown in Fig.8 (A) and (B), respectively. From these figures, in the VL model, the time-course curves of *L* and *V*_*lv*_ are almost the same shape and *L* remains constant during isovolumic phase. On the other hand, in the VFL model, changes in *L* can be observed during the two isovolumic phases. Additionally, Fig.9 illustrates the relation between *L* and *V*_*lv*_ over a cardiac cycle for both models. It can be seen that the *L*-*V*_*lv*_ relation differ between two models due to the sarcomere contraction and elongation during the isovolumic phase.

**Fig 7.**
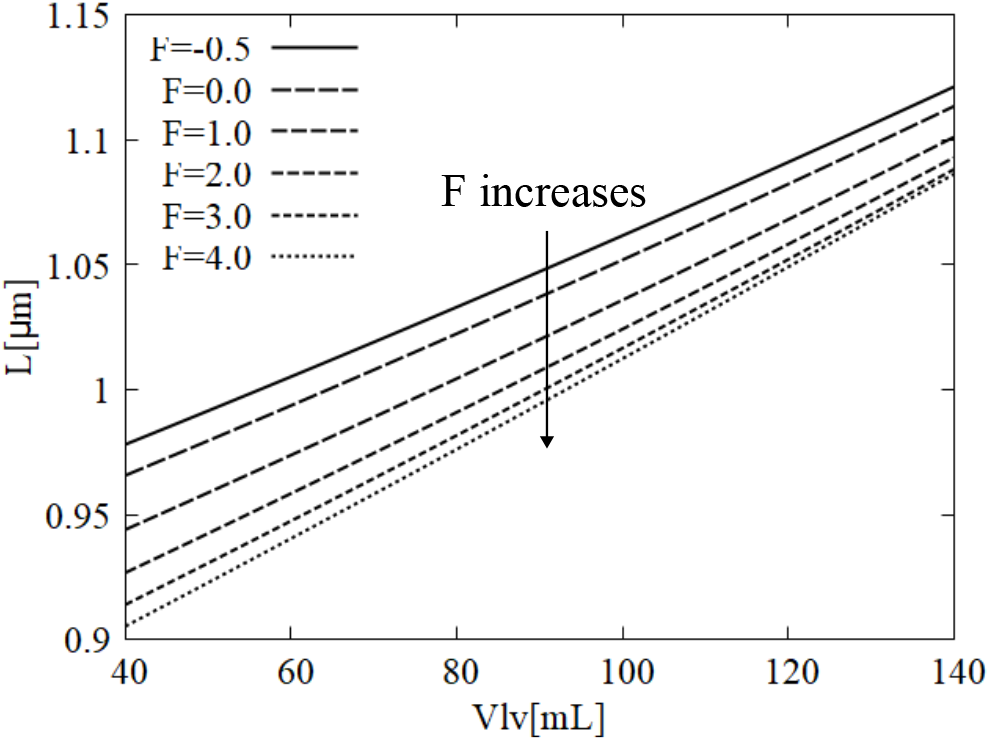
The relation between *V*_*lv*_ and *L* with fixed *F* values. The lines represent *L* − *V*_*lv*_ when several fixed F values are introduced.

**Fig 8.**
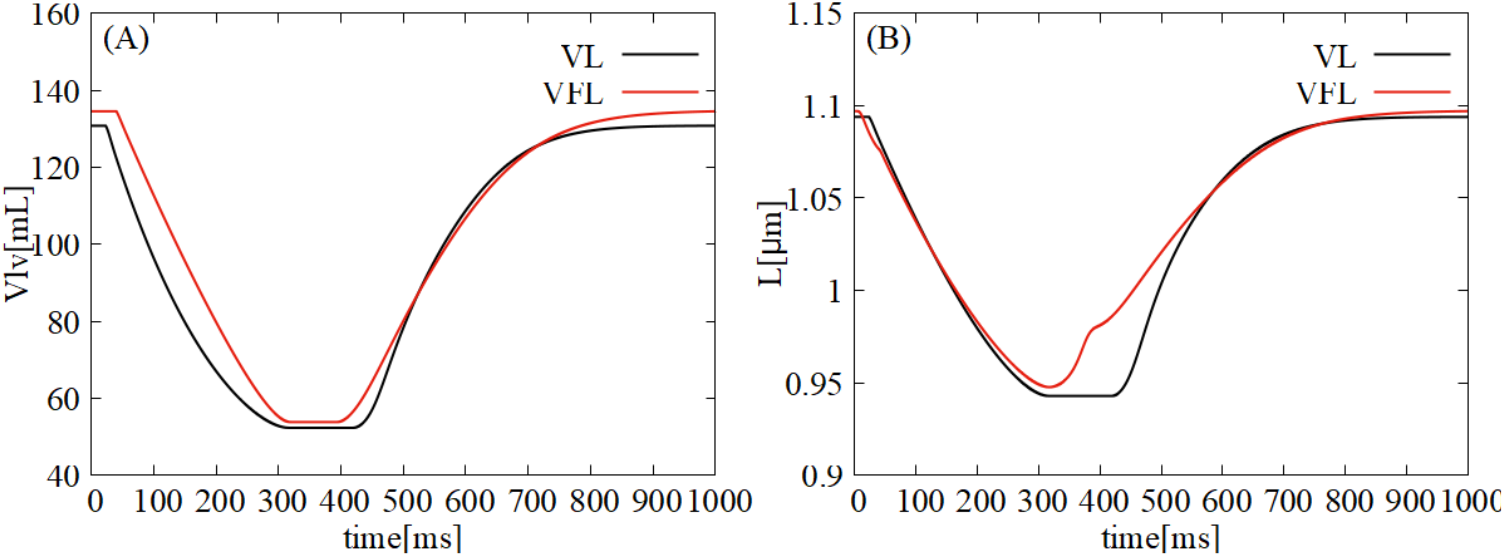
The time-course of *V*_*lv*_ and *L*.

### Hemodynamic changes induced by afterload variation

To assess the effect of afterload on the systolic and diastolic fuction such as *τ* and Tei-index, we performed simulations with the two models varying *R*_*out*_ values ( 900, 950, 1000, 1100, 1150, 1200 ). The PV loops of the VL and the VFL models with *R*_*out*_ = 900, 1200 are shown in Fig.10 (A) and (B), respectively. In both models, the slope of the ESPVR becomes positive, but the slope of the VFL model is smaller than that of the VL model.

The time courses of *V*_*lv*_, *L, F*, and *P*_*lv*_ under different afterload conditions (*R*_*out*_ = 900 and 1200) are shown in Fig. 11. Panels (A), (C), (E), and (G) present the results of the VL model, while (B), (D), (F), and (H) show those of the VFL model.

**Fig 9.**
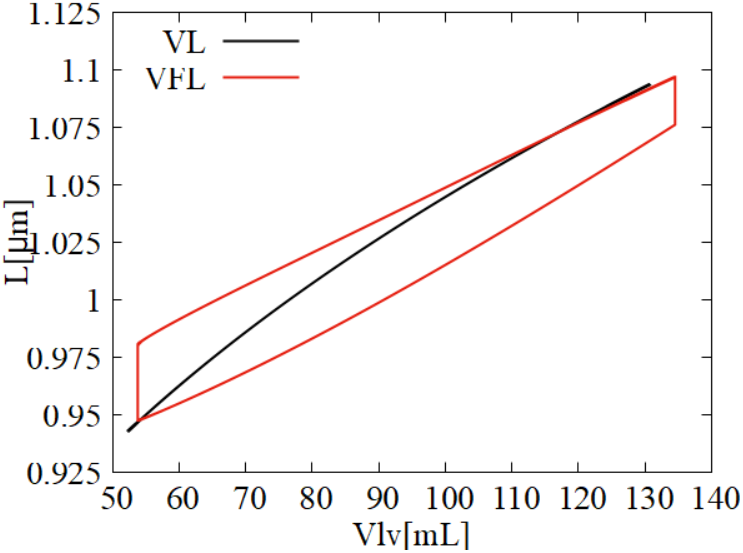
The relation between *L* and *V*_*lv*_ during a cardiac cycle.

**Fig 10.**
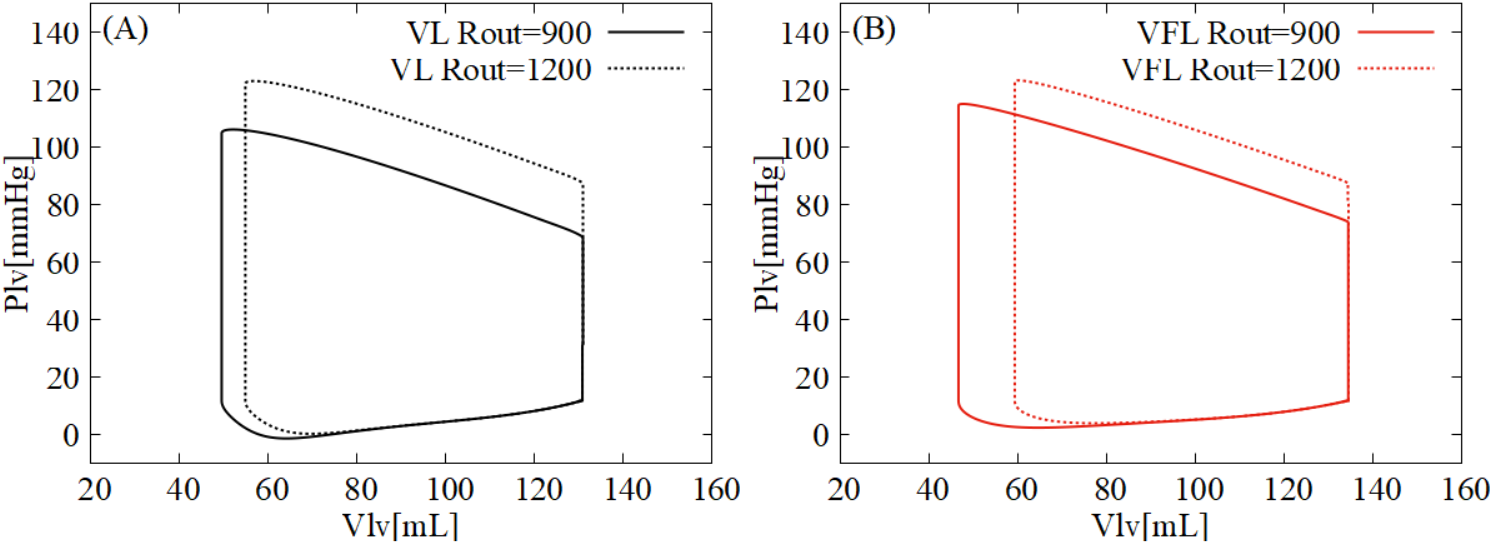
PV loops with afterload variation.

**Fig 11.**
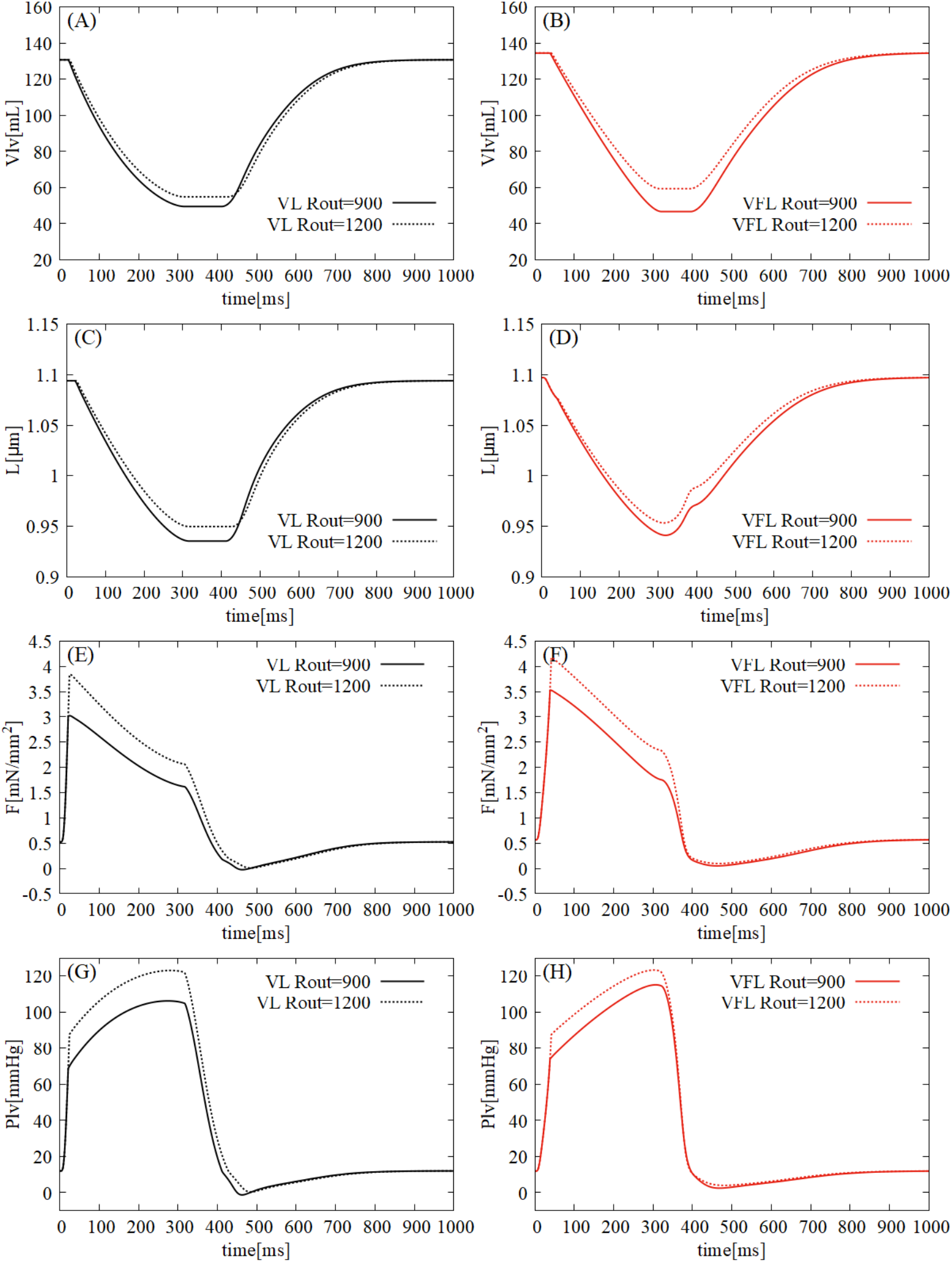
The time-course of *V*_*lv*_, *L, F* and *P*_*lv*_ with afterload variation.

Comparing panels (A) and (B), the difference in *V*_*lv*_ between the two afterload conditions is greater in the VFL model during 200–300 ms. In panels (C) and (D), the VFL model exhibits a more pronounced change in *L* during the isovolumic phases compared with the VL model. Panels (E) and (F) show that during isovolumic relaxation, *F* decreases more rapidly in the VFL model than in the VL model. Simirarly, in panels (G) and (H), the VFL model shows a steeper decline in *P*_*lv*_ during isovolumic relaxation compared to the VL model.

Then, we calculated the index values ( *EF, IV CT/ET, IV RT/ET, τ*, Tei-index ) by using the afterload variation results. Each result is illustrated in Fig.12(A-E), respectively. The calculated indices for *R*_*out*_ = 900, 1050, and 1200 along with reference values, are listed in Table 4.

**Table 4.** Index values with *R*_*out*_ = 900, 1050 and 1200.

| index | VL model |  |  | VFL model |  |  | physiological values |
| --- | --- | --- | --- | --- | --- | --- | --- |
| | $R_{out}=900$ | 1050 | 1200 | $R_{out}=900$ | 1050 | 1200 | |
| $EF$ % | 62.13 | 60.0 | 58.06 | 65.27 | 60.0 | 55.87 | 55–70 [30] |
| $IVCT$ [ms] | 18.29 | 19.78 | 21.18 | 33.21 | 35.13 | 36.58 | 30–50 [29,31] |
| $IVRT$ [ms] | 94.89 | 103.93 | 112.01 | 74.75 | 75.88 | 77.84 | 60–90 [32–34] |
| $IVCT/ET$ | 0.0619 | 0.0675 | 0.0727 | 0.1173 | 0.1260 | 0.1328 | - |
| $IVRT/ET$ | 0.3214 | 0.3544 | 0.3846 | 0.2642 | 0.2722 | 0.2826 | - |
| $\tau$ [ms] | 43.75 | 46.19 | 48.27 | 33.13 | 33.11 | 33.47 | 30–45 [35,36] |
| Tei-index | 0.3834 | 0.4219 | 0.4573 | 0.3816 | 0.3982 | 0.4154 | 0.35–0.45 [29] |

Physiological values represent typical ranges observed in healthy human LV.

**Fig 12.**
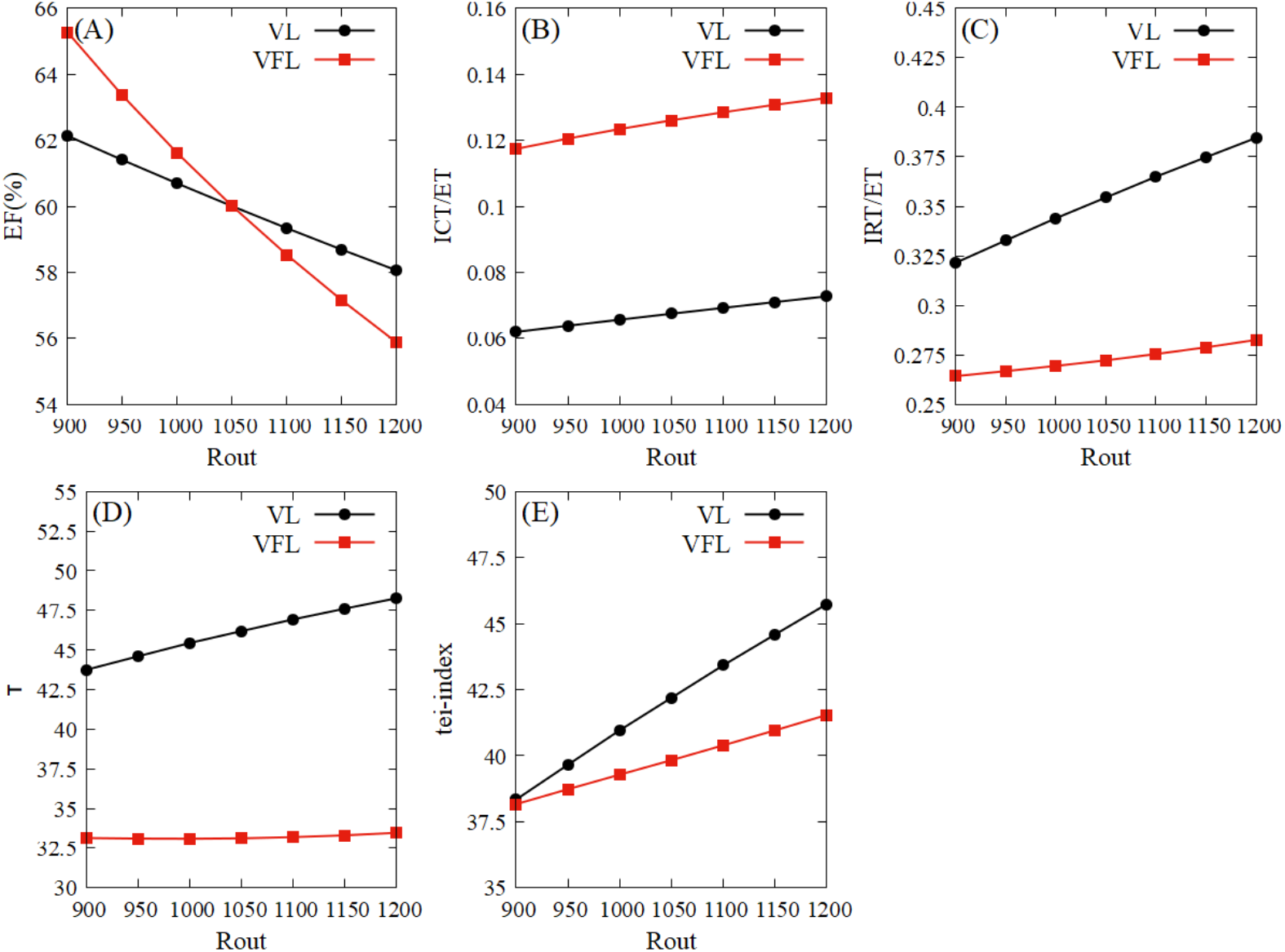
Index values with *R*_*out*_ varying.

Focusing on the condition of *R*_*out*_ = 1050, the following observations can be made: Compared to the VL model, the VFL model exhibits an approximately 75% longer isovolumic contraction time (IVCT) and a 25% shorter isovolumic relaxation time (IVRT), suggesting that the VFL model more closely reproduces physiological behavior. In addition, the Tei-index of VFL model is also smaller and closer to the physiological range from 34 to 42 than that of the VL model. These findings suggest that favorable hemodynamics are achieved when changes in *L* occur during the isovolumic phases.

As *R*_*out*_ increases, all indices except *τ* showed consistent trends in both models. However, the magnitude of changes in IVRT/ET, *τ*, and Tei-index was smaller in the VFL model compared to the VL model. This relative stability of the VFL model provides more robust diastolic function under varying afterload conditions.

### Hemodynamic changes induced by preload variation

To evaluate the effect of preload variation on the LV pressure-volume relation, we performed simulations with the two models varying *P*_*E*1_ ( 10, 11, 12, 13, 14 ). The PV loops of the VL model and the VFL model with *P*_*E*1_=10, 14 are shown in (A) and (B) of Fig.13, respectively. As shown in panel (A), increasing *P*_*E*1_ leads to increases in both the end-systolic pressure and volume, yielding a positive ESPVR slope. Conversely, panel (B) shows that increasing *P*_*E*1_ results in a decrease in the end-systolic pressure and volume, producing a nagative ESPVR slope.

Fig.14 shows the simulation results of the time-course of *V*_*lv*_, *L, F*, and *P*_*lv*_ under different preload conditions (*P*_*E*1_ = 10 and 14). The panel (A), (C), (E), and (G) represent the results of VL model, while (B), (D), (F), and (H) show those from the VFL model. In both models, *L* and *F* exhibit the parallel shift for the preload variation (*P*_*E*1_ = 10 and 14)during ejection (approximately 50-300 [ms] ). Notably, panel (B) shows that the difference in *V*_*lv*_ between two preload conditions progressively increases from 200 to 300 [ms], indicating a more pronounced volume resopnse in the VFL model during the ejection phase, compared to panel (A).

**Fig 13.**
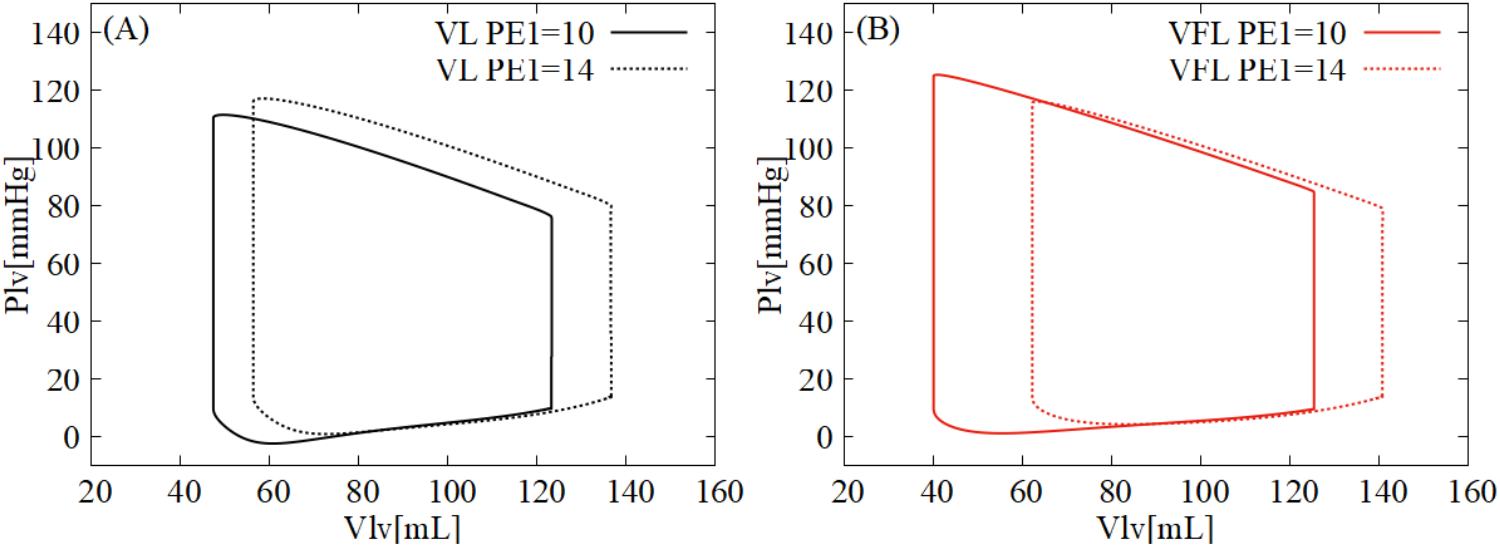
PV loops with preload variation.

**Fig 14.**
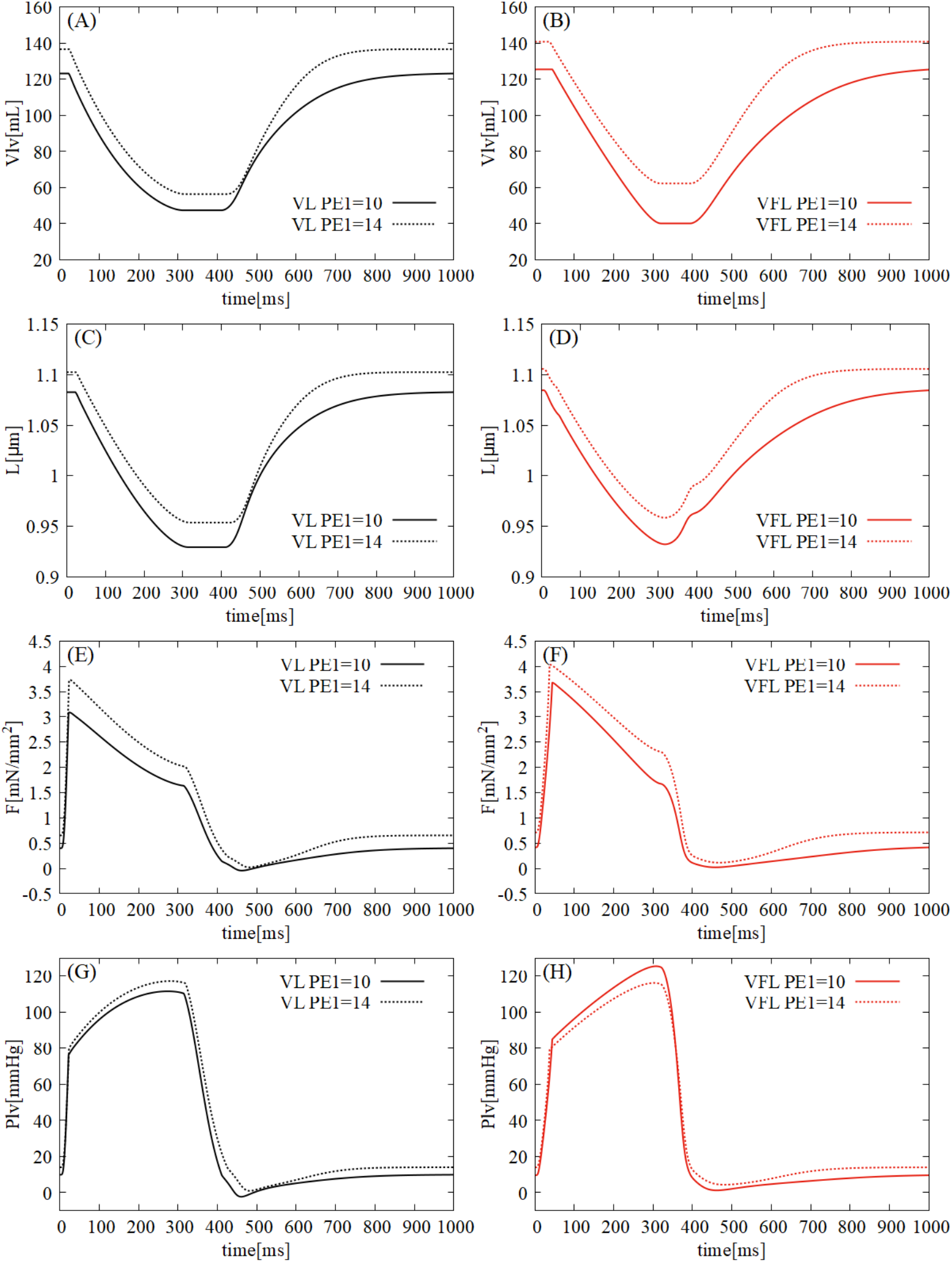
The time-course of *V*_*lv*_, *L, F* and *P*_*lv*_ with preload variation.

## Discussion

### Evaluation of the two models

From the simulation results, the VFL model exhibits a smaller increase in IVRT with afterload increase compared to the VL model. Additionaly, the VFL model shows a better maintenance of *τ* with afterload increase. These findings suggest that the sarcomere length dynamics governed by balance between LV volume and contraction force may mechanically contribute to stabilizing the relaxation performance.

Under preload variation, the VL model shows a positive ESPVR slope, reflecting normal contractile behavior, while the VFL model shows a negative slope. These differences suggest that the VL and VFL model have different mechanical roles in terms of contractile and relaxation properties. According to Rodriguez et al. [12], endocardial cells exhibit a linear sarcomere length-volume relationship similar to the VL model, while epicardial cells show a hysteresis-like relationship similar to the VFL model. Thus, endocardial properties may be important for pressure maintenance during systole, and epicardial properties for pressure reduction during diastole. To achieve a more physiological LV performance where both contraction and relaxation are properly considered, a multilayer model that incorporates both VL-like and VFL-like properties across the ventricular wall would be required.

### Analysis of the time course of contraction force

#### Comparison approach for the two models

To analyze the differences in *IV CT* and *IV RT* between the two models, we conducted two additional simulations which focus on the increase and decrease velocities of the contraction force during isovolumic phases. To ensure comparability, we used the VFL model for 99 cycles, then transitioned to the VL model at specific points in the final cycle. The first simulation transitioned to the VL model at end-diastole (“ED-transitioned model”) to analyze the increase velocity of the contraction force from end-diastole to the onset of ejection. The second simulation transitioned to the VL model at end-systole (“ES-transitioned model”) to analyze the decrease velocity of the contraction force from end-systole to the filling onset. As the control model, to compare the two results, we used the VFL model throughout all cycles without any transitions. The conditions for these analyses are summarized in Table 5.

**Table 5.** Simuation conditions for analysis.

| condition | description |
| --- | --- |
| VFL model ( control ) | VFL model throughout all the cycles without transition |
| ED-transitioned model | Transitioned from the VFL model to the VL model at end-diastole in the final cycle |
| ES-transitioned model | Transitioned from the VFL model to the VL model at end-systolic in the final cycle |

Abbreviations: ED and ES stand for end diastole and end systole, respectively.

#### The time course of active and passive force

To assess how each force in wall tension changes during the isovolumic phases, we examined the time courses of wall tension (*F* ), active force (*F*_*b*_), and passive force (*F*_*p*_) in the VFL, the ED- and ES-transitioned models.

Fig.15(A)-(C) shows results during isovolumic contraction phase (IVCP). The VFL model exhibited a faster increase in *F* and a delayed ejection onset compared to the ED-transitioned model (panel (A)). Since *F*_*b*_ closely follows *F* (panel (B)), and *K*_*ps*_*F*_*p*_ varies less than *K*_*bs*_*F*_*b*_ (panel (C)), the active component dominates *F* . Additionally, on the right-hand side of Eq.(2), *h*_*w*_ which appears in the first term becomes approximately 2% of *h*_*p*_ which appears in the second term. Therefore the wall tension (*F*_*ext*_) during isovolumic contraction phase can be approximated as Eq.(25).

**Fig 15.**
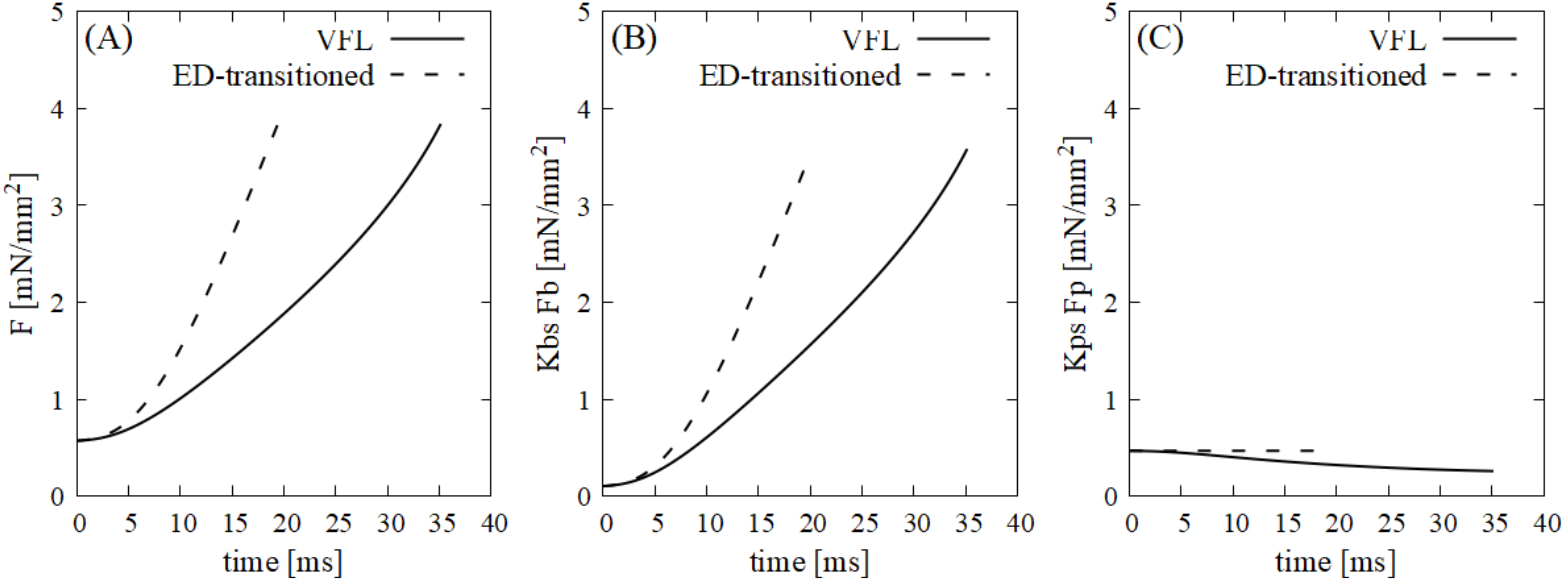
The time course of *F, F*_*b*_, and *F*_*p*_ during IVCP.

Similarly, Fig.16(A)-(C) shows results during isovolumic relaxation phase (IVRP). The VFL model initially had a higher *F* than the ES-transitioned model, but it decreased over time, falling below the ES-transitioned model at the onset of blood filling (panel (A)). Also, the time course of *F*_*b*_ was nearly the same as that of *F*, and *K*_*ps*_*F*_*p*_ changes only slightly (panels (B), (C)), indicating *K*_*bs*_*F*_*b*_ is the main contributor to *F* . These findings support the validity of the approximation shown in Eq.(25) for both IVCP and IVRP phases.

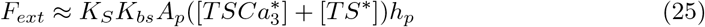

**Fig 16.**
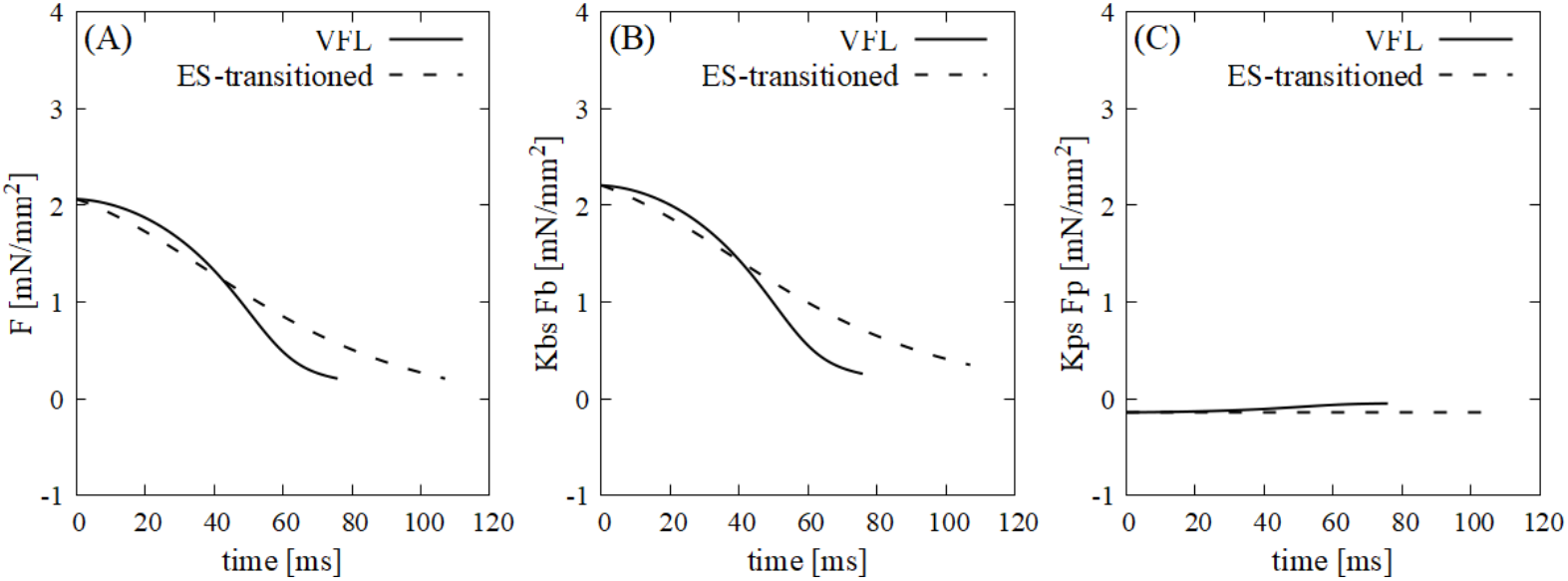
The time course of *F, F*_*b*_, and *F*_*p*_ during IVRP.

#### Recoil effect of the two models

Recoil effect is often explained as the force to restore the original length of the muscle tissue after a rapid decrease in the contraction force, and considered to be important for LV pressure decrease [8]. In this study, the effect can be understood as the parallel elastic component (*F*_*p*_) in the two models, because its force increases in the relaxation direction when shortening. As shown in Fig.16, *F*_*p*_ remains nearly constant during the IVRP in both the VFL and ES-transitioned models, compared to the change in *F*_*b*_. Therefore, within our model formulations, the recoil-related passive elastic component does not play a dominant role in LV pressure decline during relaxation; instead, the rapid force decrease is primarily governed by the force-velocity relationship of the active force (*F*_*b*_).

On the other hand, experimental LV pressure measurements, such as those reported by Rodriguez et al. [12], do not provide sufficient information to uniquely separate the contributions of active force and passive elastic force. Our results suggest that the dominant factor for LV pressure decay during the IVRP cannot be uniquely attributed to the recoil effect alone.

#### Underlying crossbridge and troponin mechanisms for active force generation

To discuss how *h*_*p*_ and 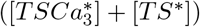 affect *F*_*ext*_, we examined the changes in contraction model variables (Fig.17). Panel (A) shows *h*_*p*_ remains nearly constant in the ED-transitioned model, but decreases by up to 25% in the VFL model. Panel (B) shows 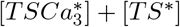 is similar in both models for the initial 10 ms, but the slope becomes smaller in the VFL model later. Thus, during early IVCP, *F*_*ext*_ increases due to changes in *h*_*p*_, while later it is influenced by both *h*_*p*_ and 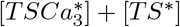.

**Fig 17.**
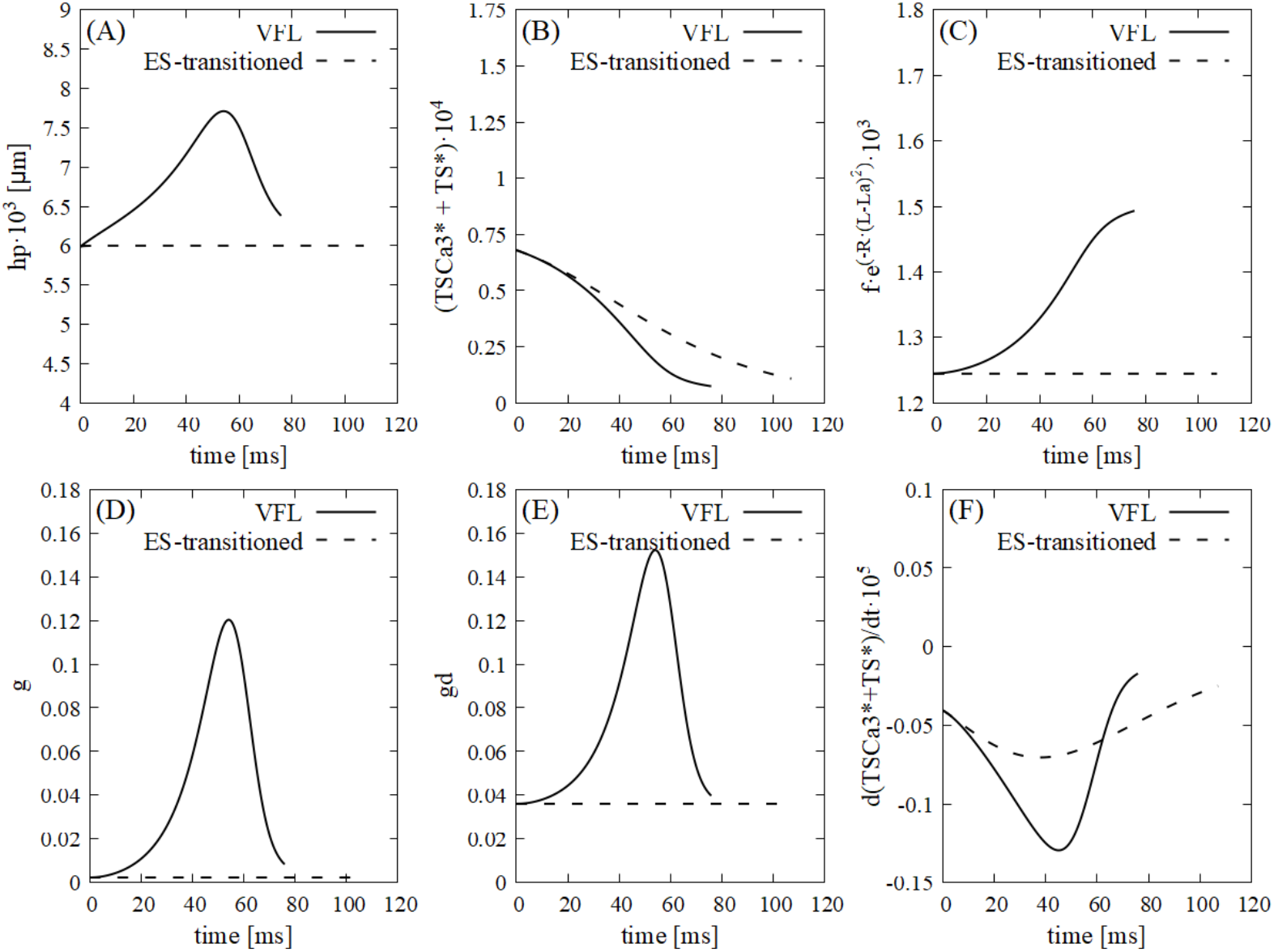
Changes in variables of contraction model during IVCP.

Panel (C) shows the overlap function 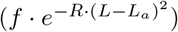 decreases with time in the VFL model, but remains constant in the ED-transitioned model. Panels (D) and (E) show crossbridge detachment velocities (*g, g*_*d*_) increasing and decreasing with sarcomere shortening in the VFL model, but remaining constant in the ED-transitioned model. Panel (F) shows the rate of increase in 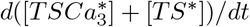 is similar in both models for the initial 10 ms, but lower in the VFL model later.

In summary, sarcomere length changes during IVCP reduce actin-myosin overlap and increase crossbridge detachment, slowing the increase in 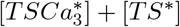. This leads to a smaller pressure increase and prolongs IVCP.

Similarly, during IVRP (Fig.18), *h*_*p*_ increases and peaks at 40-60 ms in the VFL model, but remains constant in the ES-transitioned model. 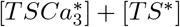 is similar initially, but the slope is smaller in the VFL model later. The overlap function and crossbridge detachment velocities show similar trends as in IVCP. The rate of decrease in 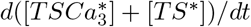 is lower in the VFL model after 60 ms.

**Fig 18.**
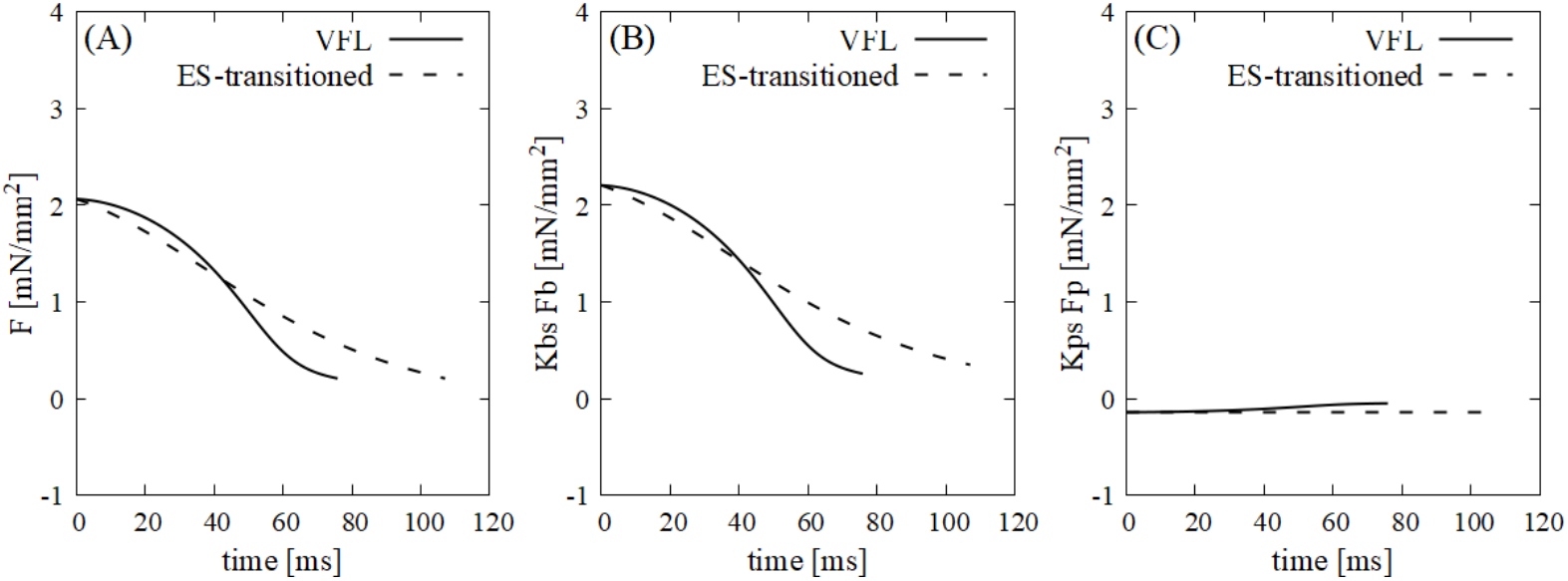
Changes in variables of contraction model during IVRP.

Thus, sarcomere lengthening during IVRP increases actin-myosin overlap and crossbridge detachment, accelerating LV pressure decrease and shortening IVRP.

### Analysis of isovolumic time for afterload variation

#### Derivation of analytical expressions for isovolumic time

To analyze the effect of afterload(*R*_*out*_) on IVCT and IVRT, we derived equations describing the impact of *R*_*out*_ increase on IVCT and IVRT. The discussion is based on the simulation results and the equations derived from the model equations.

- For IVCT analysis *IV CT* can be defined as the time required for the LV pressure to increase from the end-diastole to the onset of ejection.

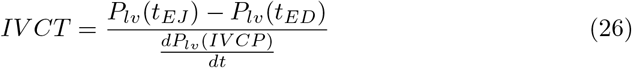

Here, 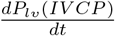 denotes the increase of *P*_*lv*_ during IVCP. As *P*_*lv*_(*t*_*ED*_) is always constant under a fixed preload(*P*_*E*1_) in our circulation model, *P*_*lv*_(*t*_*EJ*_ ) is the dominant determinant of *IV CT* . Additionally, since the LV volume(*V*_*lv*_) remains constant during IVCP, the wall thickness(*h*_*lv*_) and internal radius(*R*_*lv*_) are also constant. Therefore, 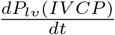 in the denominator of Eq.(26) can be obtained by differentiating Eq.(10) with time(*t*).

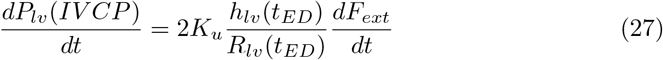

Here, 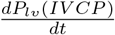 is determined by *V*_*lv*_(*t*_*ED*_) and 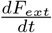, as *h*_*lv*_(*t*_*ED*_) and *R*_*lv*_(*t*_*ED*_) are derived from *V*_*lv*_(*t*_*ED*_) as shown in Eqs. (12) and (13). Moreover, changes in *IV CT* (Δ*IV CT* ) with *R*_*out*_ can be expressed as follows, when the increase of *P*_*lv*_ at the onset of ejection with afterload is represented as Δ*P*_*lv*_(*t*_*EJ*_ ).

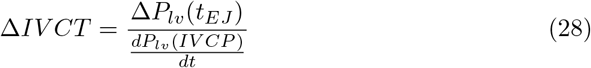
- For IVRT analysis IVRT can be defined as the time required for the LV pressure to decrease from the end-systole(*P*_*lv*_(*t*_*ES*_)) to the onset of blood filling(*P*_*lv*_(*t*_*F L*_)).

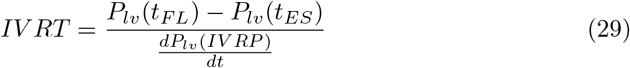

Here, 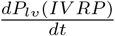 denotes the increase of *P*_*lv*_ during IVRP. As *P*_*lv*_(*t*_*F L*_) is always constant under a fixed preload(*P*_*E*1_) in our circulation model, *P*_*lv*_(*t*_*ES*_) is the dominant determinant of *IV RT* . Additionally, as with Eq. (27),since the LV volume(*V*_*lv*_) remains constant during IVRP, the wall thickness(*h*_*lv*_) and internal radius(*R*_*lv*_) are also constant. Therefore, 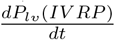 in the denominator of Eq. (29) can be obtained by differentiating Eq. (10) by time (*t*).

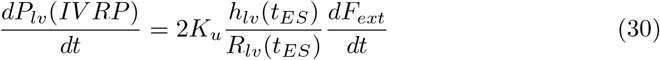

Here, 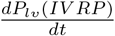 is determined by *V*_*lv*_(*t*_*ES*_) and 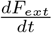, as *h*_*lv*_(*t*_*ES*_) and *R*_*lv*_(*t*_*ES*_) are derived from *V*_*lv*_(*t*_*ES*_) as shown in Eqs. (12) and (13). Also, 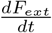 changes depending on the value of *R*_*out*_. Moreover, changes in *IV RT* (Δ*IV RT* ) with *R*_*out*_ can be expressed as follows.

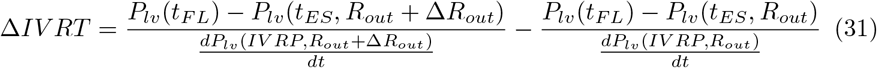

Note that *P*_*lv*_(*t*_*ES*_, *R*_*out*_) and *P*_*lv*_(*t*_*ES*_, *R*_*out*_ + Δ*R*_*out*_) are *P*_*lv*_ at end-systole in the condition of *R*_*out*_ and *R*_*out*_ + Δ*R*_*out*_, respectively. Similarly, 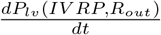 and 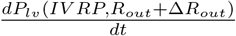 are derived from Eq.(30) in the condition of *R*_*out*_ and *R*_*out*_ + Δ*R*_*out*_, respectively.

#### Determinants and mechanistic interpretation

To clarify how afterload (*R*_*out*_) influences the isovolumic time indices (*IV CT, IV RT* ), the simulation results were substituted into Eqs.(26)-(31). The determinants and computed values are summarized in Table 6 and Table 7.

**Table 6.** IVCT determinant values.

| determinant | VL model | VFL model |
| --- | --- | --- |
| $P_{lv}(t_{EJ}, R_{out} = 900)$ | 68.45 | 73.53 |
| $P_{lv}(t_{EJ}, R_{out} = 1200)$ | 87.35 | 87.14 |
| $\Delta P_{lv}(t_{EJ})$ | 18.90 | 13.61 |
| $h_{lv}(t_{ED})/R_{lv}(t_{ED})$ | 0.40 | 0.39 |
| $dF_{ext}/dt$ | 1.09 | 0.70 |
| $dP_{lv}(IVCP)/dt$ | 6.54 | 4.10 |
| $\Delta IVCT$ | 2.89 | 3.33 |

**Table 7.** IVRT determinant values.

| determinant | VL model | VFL model |
| --- | --- | --- |
| $P_{lv}(t_{ES}, R_{out} = 900)$ | 104.901 | 114.493 |
| $P_{lv}(t_{ES}, R_{out} = 1200)$ | 122.093 | 122.780 |
| $h_{lv}/R_{lv}\vert t_{ES}, R_{out} = 900$ | 1.141 | 1.202 |
| $h_{lv}/R_{lv}\vert t_{ES}, R_{out} = 1200$ | 1.039 | 0.965 |
| $dF_{ext}/dt\vert R_{out} = 900$ | -0.057 | -0.076 |
| $dF_{ext}/dt\vert R_{out} = 1200$ | -0.063 | -0.098 |
| $dP_{lv}(IVRP)/dt\vert R_{out} = 900$ | -0.979 | -1.371 |
| $dP_{lv}(IVRP)/dt\vert R_{out} = 1200$ | -0.983 | -1.423 |
| $\Delta IVRT$ | 17.122 | 3.090 |

Both models showed an increase in IVCT as *R*_*out*_ increased. However, the magnitude of change was slightly larger in the VFL model (+3.3 ms) than in the VL model (+2.9 ms), which are consistent with the theoretical values computed from Eq. (38). The major determinant of IVCT was the rate of force development (*dF*_*ext*_*/dt*), whereas the geometric term (*h*_*lvED*_*/R*_*lvED*_) remained nearly constant across conditions. In the VFL model, sarcomere shortening during the isovolumic contraction phase reduced (*dF*_*ext*_*/dt*) by approximately 35%, leading to a slower pressure rise (*dP*_*lv*_(*IV CP* )*/dt*) and a prolonged IVCT compared to the VL model.

IVRT also increased with higher *R*_*out*_, but the extent of increase was substantially smaller in the VFL model (+3 ms) than in the VL model (+17 ms), as summarized in Table 7. According to Eq.(31), *dF*_*ext*_*/dt* during the relaxation phase dominantly determines the rate of pressure decay. This difference can be attributed to cross-bridge dynamics during the isovolumic relaxation phase, as shown in Fig. 19. Specifically, the peak cross-bridge extension (*h*_*p*_) and the detachment-related velocity components (*g, g*_*d*_) increase with *R*_*out*_ in the VFL model, promoting faster cross-bridge detachment and thereby enhancing force decay. In the VFL model, sarcomere lengthening enhanced cross-bridge detachment and maintained a faster decline in *F*_*ext*_, even under high afterload. Consequently, *dP*_*lv*_(*IV RP* )*/dt* remained greater in magnitude, resulting in a more stable IVRT against afterload elevation.

**Fig 19.**
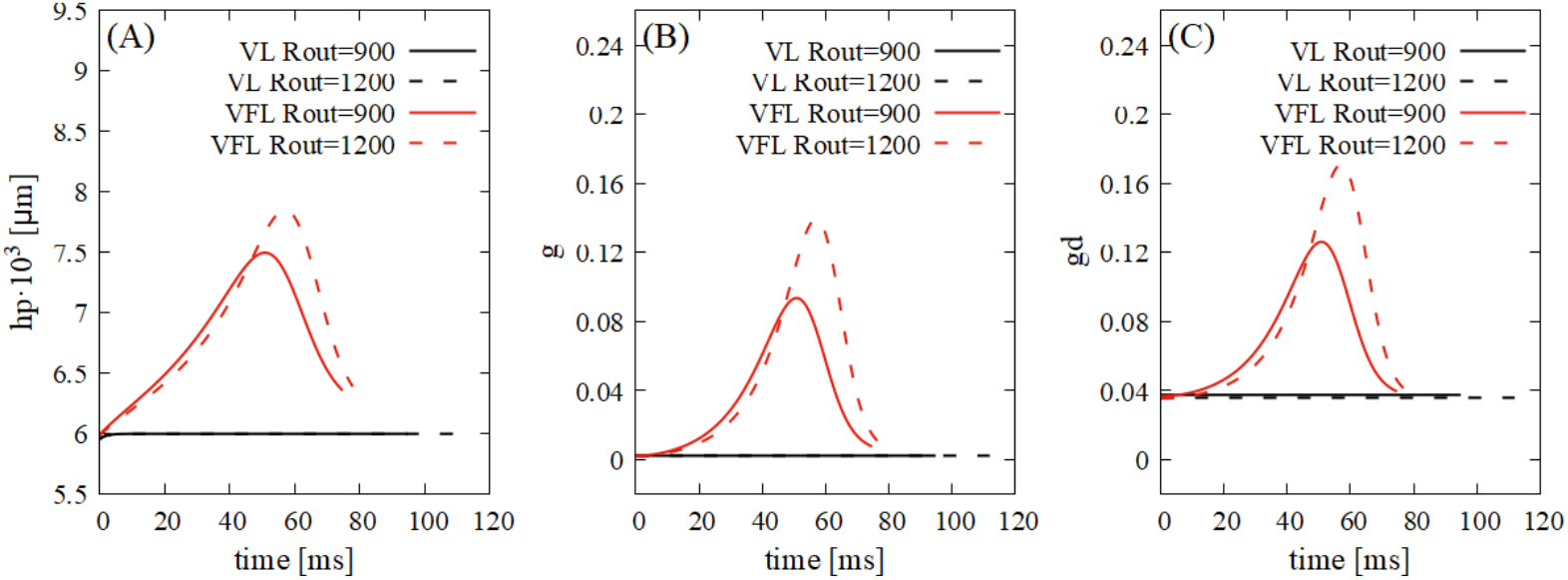
The time course of *h*_*p*_, *g*, and *g*_*d*_ during IVRP.

Overall, these analyses indicate that incorporating sarcomere length variation (the VFL model) slows the force development during isovolumic contraction but accelerates force decay during relaxation. These effects produce a prolonged IVCT and stabilized IVRT under increased afterload, reproducing physiological left ventricular behavior more accurately than the conventional VL model.

### Mechanism of the negative ESPVR slope in the VFL model

The VFL model showed that the slope of the ESPVR became negative as preload increased. From Fig.14 (H), the LV pressure at the onset of ejection with *P*_*E*1_ = 14 was lower than that with *P*_*E*1_ = 10. Because the Windkessel-based aortic component causes the end-systolic aortic pressure to influence the LV pressure in the subsequent cycle, we examined *V*_*lv*_, *L, F* and *P*_*lv*_ when the pressure difference first appears. Fig.20 shows these variables during the first two cycles. Around 1300 [ms], panels (G) and (H) show that in the VL model, the end-systolic pressure increases with increasing *P*_*E*1_, while in the VFL model, it decreases. During ejection (approximately 1100-1300 [ms]), both models exhibit nearly parallel shifts with increasing *P*_*E*1_ (panels (C)-(F)). However, as shown in panels (A) and (B), the VFL model shows a larger increase in *V*_*lv*_ with preload, which leads to a relative reduction in wall tension and thus lower LV pressure. This indicates that the pressure decrease arises from the larger volume change in the VFL model, a consequence of the model’s coupling between sarcomere length and contraction force.

We then examined the *L*-*V*_*lv*_ relation during ejection phase (Fig.21) using model equation curves with fixed *F* . In the figure, the VL model exhibits an upward-convex curve, while the VFL model shows a downward-convex curve. From the model-derived curves, the curve shifts downward with increasing *F*, which explains the VFL model having a downward-convex trajectory during the ejection phase due to the decreasing *F* with time. At end-systole, *L* is similar between the two models, but *V*_*lv*_ differs significantly, indicating that the VFL model is more sensitive to end-systolic volume changes, contributing to the negative ESPVR slope.

**Fig 20.**
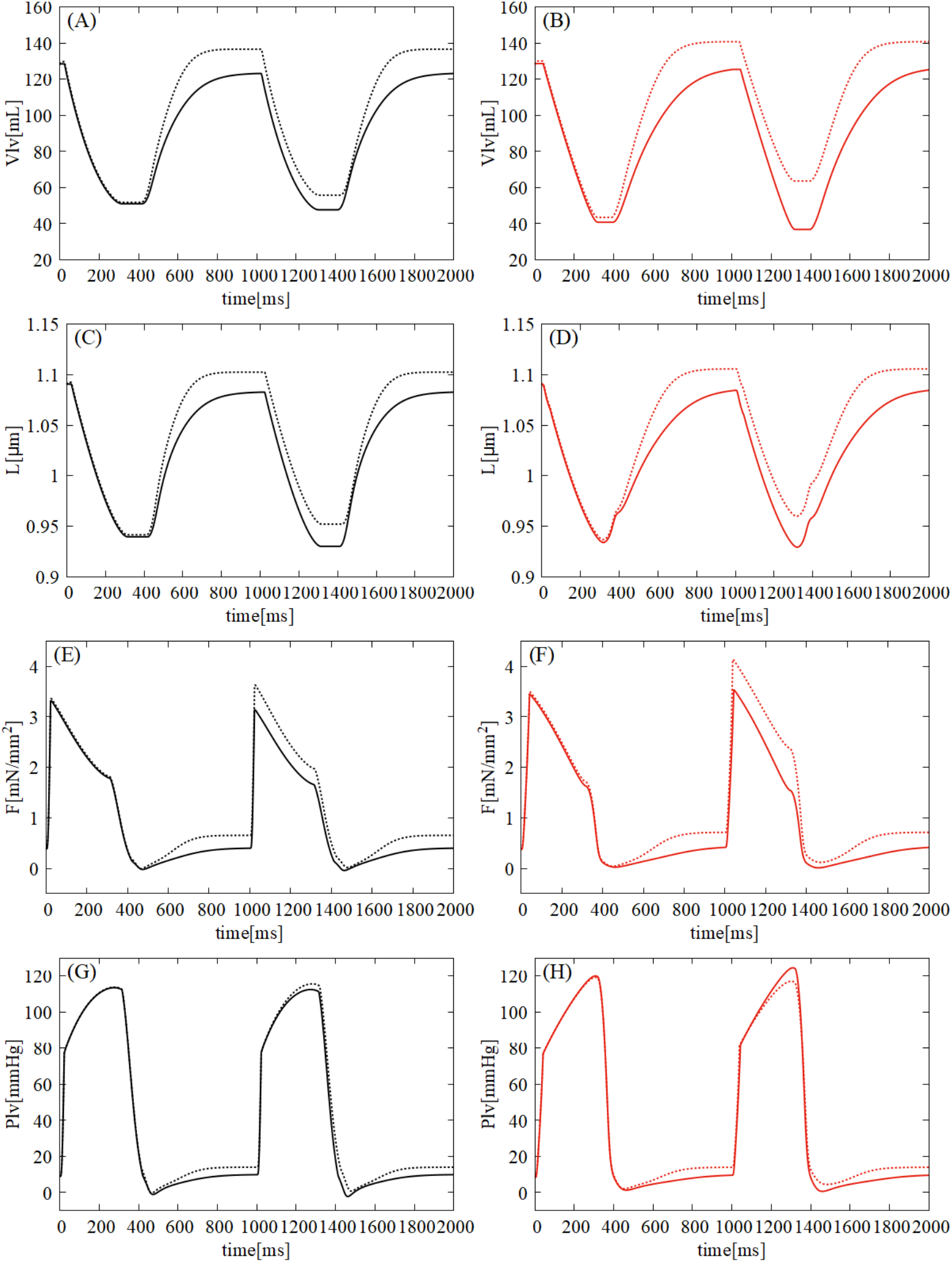
The time-course of *V*_*lv*_, *L, F* and *P*_*lv*_ with preload variation during first two cycles Solid and dashed lines represent conditions with *P*_*E*1_ = 10 and *P*_*E*1_ = 14, respectively. Black lines indicate the VL model, and red lines indicate the VFL model.

### Limitations of simulation model

In this study, we constructed VL model and VFL model as single-layer models for simplified analysis. Incorporating multi-layer structure model may allow a more physiologically realistic analysis. However, developing such a model presents challenges, including computational complexity and numerical instability, which should be addressed in future work.

Furthermore, the simulation results of the VFL model are insufficient to fully interpret the measured data by Rodriguez et al. [12] although it was constructed to reproduce the hysteresis relationship between sarcomere length and LV volume in the epicardium based on the reported data. Fig. 22 shows the comparison of the − *F*_*ext*_ − *V*_*lv*_ − *SL* loop between the reported data by Rodriguez et al., model equation and simulation result which is scaled into canine LV by using Eqs. (21), (22). Both of the reported data’s trajectory and simulation result’s are plotted on the model equation’s plane, however their trajectories do not coincide. This is because the force scaling was constrained to avoid unstable regions in the *F*_*ext*_ − *V*_*lv*_ − *SL* space, rather than to match the trajectories.

The primary aim of this study was to theoretically evaluate the impact of sarcomere length dynamics during isovolumic phases on hemodynamics, rather than to quantitatively fit experimental trajectories. Thus, we did not attempt to adjust the VFL model parameters to reproduce the experimental data. Nevertheless, comparison between the VL and VFL models provides a useful theoretical framework for understanding how sarcomere length changes during isovolumic phases influence LV mechanics and hemodynamics.

## Conclusion

Many studies have suggested that the myocyte length change during isovolumic phase contributes to the diastolic relaxation efficiency. Experimental observations have shown that the sarcomere length dynamics during isovolumic phases differ across the ventricular wall layers : while the relationship between LV volume and sarcomere length is nearly linear in the endocardium, it exhibits a hysteresis curve in the epicardium.

**Fig 21.**
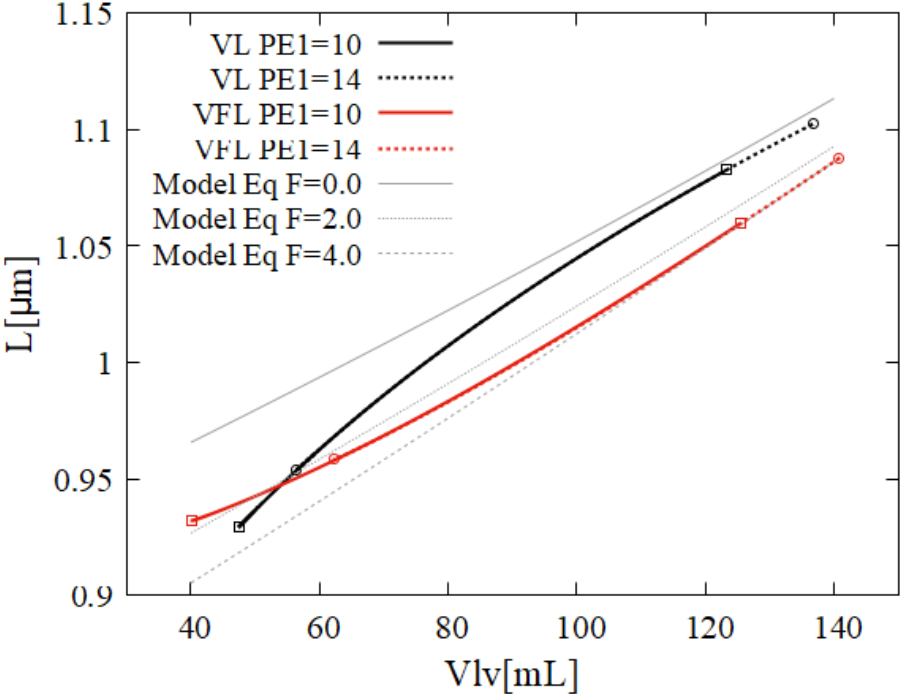
*L*-*V*_*lv*_ relation of the two models during ejection phase. Square and circle markers indicate the results of *P*_*E*1_ = 10 and *P*_*E*1_ = 14, respectively. Model Eq curves(gray) shows the results of Eq.(15) with fixed values *F* (0.0, 2.0, and 4.0).

**Fig 22.**
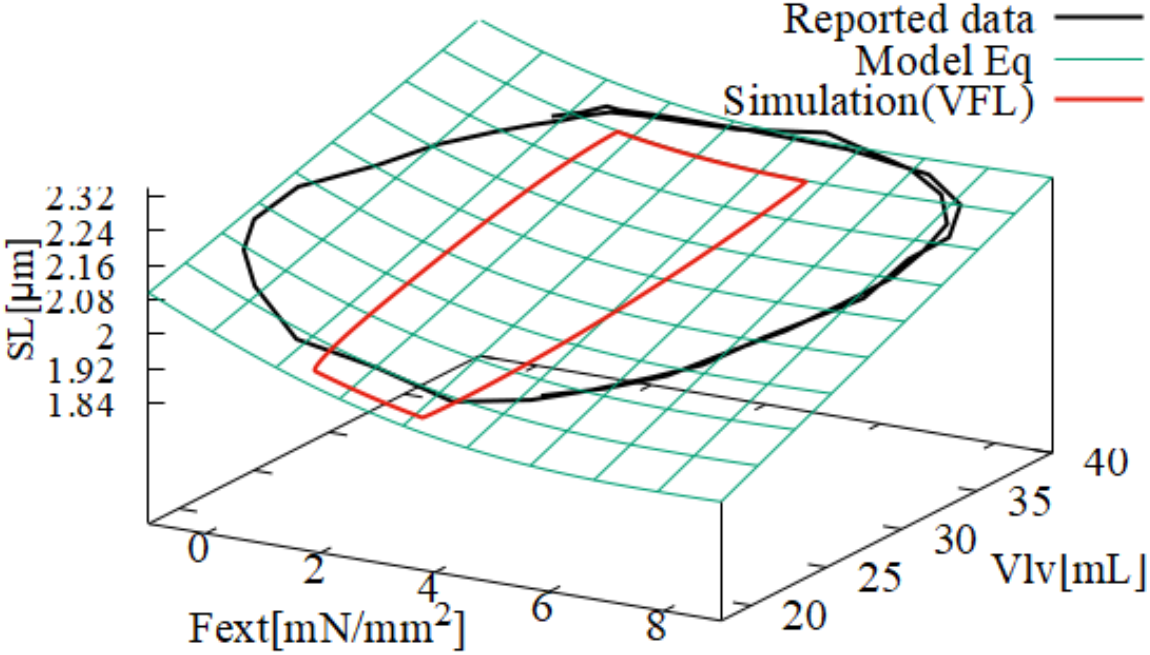
Comparison of *F*_*ext*_ − *V*_*lv*_ − *SL* loop between the reported data by Rodriguez et al., model equation and simulation result. Black and red lines represent Reported data by Rodriguez et al. [12] and simulation result of the VFL model. Green surface represents the model equation of the VFL model (Eq.(15)).

In this study, to theoretically investigate the effect of the sarcomere length change during isovolumic phases on hemodynamics, we constructed two LV contraction models: a volume-based length (VL) model representing the LV volume and sarcomere length relationship in the endocardium, and a volume-force-coupled length (VFL) model representing the epicardial behavior. The VFL model showed superior relaxation performance but relatively reduced contraction performance compared with the VL model.

These results suggest that such performance differences may reflect the structure of the actual LV wall which incorporates both epicardial and endocardial volume–sarcomere length characteristics. The findings of this study further highlight that, even under constant volume conditions, layer-dependent sarcomere length dynamics induce force-velocity relationship effects that influence LV pressure decay and isovolumic relaxation time.

## Supporting information

SourceCode

ModelDescription

## Supporting information

**S1 Fig. Bold the title sentence**. Add descriptive text after the title of the item (optional).

**S2 Fig. Lorem ipsum**. Maecenas convallis mauris sit amet sem ultrices gravida. Etiam eget sapien nibh. Sed ac ipsum eget enim egestas ullamcorper nec euismod ligula. Curabitur fringilla pulvinar lectus consectetur pellentesque.

**S1 File. Lorem ipsum**. Maecenas convallis mauris sit amet sem ultrices gravida. Etiam eget sapien nibh. Sed ac ipsum eget enim egestas ullamcorper nec euismod ligula. Curabitur fringilla pulvinar lectus consectetur pellentesque.

**S1 Video. Lorem ipsum**. Maecenas convallis mauris sit amet sem ultrices gravida. Etiam eget sapien nibh. Sed ac ipsum eget enim egestas ullamcorper nec euismod ligula. Curabitur fringilla pulvinar lectus consectetur pellentesque.

**S1 Appendix. Lorem ipsum**. Maecenas convallis mauris sit amet sem ultrices gravida. Etiam eget sapien nibh. Sed ac ipsum eget enim egestas ullamcorper nec euismod ligula. Curabitur fringilla pulvinar lectus consectetur pellentesque.

**S1 Table. Lorem ipsum**. Maecenas convallis mauris sit amet sem ultrices gravida. Etiam eget sapien nibh. Sed ac ipsum eget enim egestas ullamcorper nec euismod ligula. Curabitur fringilla pulvinar lectus consectetur pellentesque.

## Acknowledgments

Cras egestas velit mauris, eu mollis turpis pellentesque sit amet. Interdum et malesuada fames ac ante ipsum primis in faucibus. Nam id pretium nisi. Sed ac quam id nisi malesuada congue. Sed interdum aliquet augue, at pellentesque quam rhoncus vitae.

## Notes

### Competing Interest Statement

The authors have declared no competing interest.

### Summary of Updates

Supplemntary materials were uploaded to the system.

