## Supplementary material for "A Theoretical Framework for the Hemodynamic Role of Sarcomere Length Dynamics During the Isovolumic Phase of the Left Ventricle": SourceCode: summary_design.html


 

### Summary designs of simulation models

#### Models Description

##### circulation model

The following 2 models are created with strong-coupled method.  
These models can be changed with option value(setting file).

- volume-based length model : VL model
- volume-force coupled length model : VFL model

#### Model Diagrams

#### Partially strong-coupled hemodynamic model

In this study, two models(VL model and VFL model) simulates hemodynamics.  
These models are made with partial strong-couple using the following calculation flow.

##### Sequence Diagram
