## Supplementary material for "A Theoretical Framework for the Hemodynamic Role of Sarcomere Length Dynamics During the Isovolumic Phase of the Left Ventricle": SourceCode: summary_design.md

## Summary designs of simulation models
### Models Description
#### circulation model
The following 2 models are created with strong-coupled method.
These models can be changed with option value(setting file).
- volume-based length model : VL model
- volume-force coupled length model : VFL model

### Model Diagrams
### Partially strong-coupled hemodynamic model
In this study, two models(VL model and VFL model) simulates hemodynamics.
These models are made with partial strong-couple using the following calculation flow.

![](./computational_method_V2.svg)

<!-- ```puml
Start
:Simulation start;
while(simulation cycle <= 100 ?) is (yes)
    repeat
        :Step 1. Determine a V<sub>lv,A</sub>, and calculate R<sub>lv</sub> and h<sub>lv</sub>;
        repeat
            :Step 2. Determine a L<sub>A</sub>, and calculate F<sub>ext</sub>;
            if(which model?) then(VL model)
                :Step 3. Calculate L<sub>B</sub> from V<sub>lv,A</sub>;
            else(VFL model)
                :Step 3'. Calculate L<sub>B</sub> from V<sub>lv,A</sub> and F<sub>ext</sub>;
            endif
            backward:change L<sub>A</sub>;
        repeat while (Step 4. abs(L<sub>A</sub>-L<sub>B</sub>) < (L tolerance) ) is (no) not (yes)
        :Step 5. Calculate P<sub>lv</sub> from R<sub>lv</sub>, h<sub>lv</sub> and F<sub>ext</sub>;
        :Step 6. Calculate dV<sub>lv</sub>/dt from P<sub>a</sub> and P<sub>lv</sub>;
        :Step 7. Calculate V<sub>lv,B</sub> from dV<sub>lv</sub>/dt and the previous V<sub>lv</sub>;
        backward:change V<sub>lv,A</sub>;
    repeat while(Step 8. abs(V<sub>lv,A</sub>-V<sub>lv,B</sub>) < (V<sub>lv</sub> tolerance) ) is (no) not (yes)
    :Step 9. Calculate NL08 model's chemical state \nwith the forward euler method;
endwhile (no)
:Simulation end;
End
``` -->


#### Sequence Diagram

![](./sequence_diagram.svg)

<!-- ```puml
participant main as main
participant conf_reader as cr
participant csvmaker as cm
participant vol_coupling as volco
participant rad_by_sphere as rad
participant wall_thickness_by_rad as wall
participant length_coupling as lenco
participant NL08 as nl
participant length_by_volume as lenbi
participant length_by_volume_force as lentri
participant laplace as lap
participant windkessel as wk

== check and get variables ==
main -> main : get configuration ini file
main -> cr : open ini file \nand get varibles
cr -> main : success or fail
alt fail
main -> main : error exit
end 
== simulation ==
main -> cm : create csv file
cm -> main : success or fail

loop t < duration

main -> volco : t
loop abs( Vlv_A - Vlv_B ) < volume_tolerance
volco -> volco : Determine Vlv_A
volco -> rad : Vlv_A
rad ..> volco : Rlv
volco -> wall : Rlv
wall ..> volco : hlv
volco -> lenco : Vlv_A
loop abs( L_A - L_B ) < length_tolerance
lenco -> lenco : Determine L_A
lenco ..> nl : L_A
nl ..> lenco : Fext
alt VL model 
lenco -> lenbi : Vlv_A
lenbi ..> lenco : L_B
else VFL model
lenco -> lentri : Vlv_A, Fext
lentri -> lenco : L_B
end
end
lenco -> lenco : calculate L \nfrom L_A and L_B
lenco ..> volco : L
volco -> lap : Fext, hlv, Rlv
lap ..> volco : Plv
volco ..> wk : Plv
wk -> wk: derive dVlv/dt and Vlv_B \nfrom prev Vlv
wk ..> volco : Vlv_B
end
volco -> volco : calculate Vlv \nfrom Vlv_A and Vlv_B
volco ..> main : Vlv, L
main -> main : set Vlv,L,Fext,Rlv,hlv,Plv to global values
main -> nl : L,t
nl -> nl : calculate chemical \nstate in NL08
nl ..> main : 
main -> cm : fprint variables into csv file
cm ..> main : 

end
``` -->
