## Supplementary material for "A Theoretical Framework for the Hemodynamic Role of Sarcomere Length Dynamics During the Isovolumic Phase of the Left Ventricle": SourceCode: simulation_guide.html


 

### Simulation guide

This document explains how to use this package.

#### Environment Prerequisites

- OS

  - Windows11,Windows10
  - Linux(Alma-Linux, RHEL)
- Compiler

  - GCC compiler

#### Compile

- for **windows**

  1. change the directory under circulation\_IsovolumicPhaseAnalysis\simulations\build\ in file explorer
  2. double-click the file named **"build.bat"**
  3. automatically opened command prompt, start compiling
  4. after compiling, **run\_hemodynamic\_model.exe** is generated under "circulation\_IsovolumicPhaseAnalysis\simulations\bin\hemodynamic\_model"
- for **linux**

  1. change the directory under circulation\_IsovolumicPhaseAnalysis/simulations/build/hemodynamic\_model/
  2. execute the following command

  ```
  ./bash build.sh
  ```

  3. after compiling, **run\_hemodynamic\_model.exe** is generated under "circulation\_IsovolumicPhaseAnalysis/simulations/bin/hemodynamic\_model/"

#### Model Usages

##### Hemodynamic Model

###### Configuring Simulation Conditions

- Hemodynamic model has 2 simulation executions (volume-based length model: VL model, volume-force-coupled length model: VFL model) in this package.
- These execution are changed by selecting flags in configuration file (.ini).
- Edit the variables in "circulation\_IsovolumicLchanges/simulations/configuration/hemodynamic\_model\_variables.ini" and save it.  
  The main simulation conditions are explained as followings.

| Variable | Type | Unit | Description |
| --- | --- | --- | --- |
| duration | double | ms | Total simulation time. |
| dt | double | ms | Time step. |
| cycle | double | ms | Duration of one cardiac cycle. |
| model\_flag | int (0–1) | — | 0: VL model :Volume-dependent length model.   1: VFL model :Volume-force coupled length model. |
| analysis\_flag | int (0–2) | — | 0: Full-cycle simulation.   1: ED-transition : Model transitions from VFL model to VL model at end-diastole of final cycle.   2: ES-transition : Model transitions from VFL model to VL model at end-systole of final cycle. |
| bisection\_volume\_scope | double | — | Initial search range for volume solver. |
| bisection\_length\_scope | double | — | Initial search range for length solver. |
| volume\_tolerance | double | — | Convergence tolerance for volume solver. |
| length\_tolerance | double | — | Convergence tolerance for length solver. |
| Rout | double | mmHg·s/ml | Outflow resistance (afterload). |
| Rpv | double | mmHg·s/ml | Pulmonary venous resistance. |
| Rlo | double | mmHg·s/ml | Left ventricular outflow resistance. |
| PE1 | double | mmHg | pulmonary venous pressure. |
| PE2 | double | mmHg | peripheral pressure. |
| CA | double | ml/mmHg | Aortic compliance. |
| save\_interval | double | ms | Time interval for saving output data. |
| save\_start\_time | double | ms | Simulation time at which output saving begins. |
| output\_dir | string | — | Directory for output CSV files. |
| configuration\_print | int (0 or 1) | — | Print (1) or suppress (0) calculated parameters at final cycle. |

###### Simulation Execution

1. change the directory under circulation\_IsovolumicPhaseAnalysis/simulations/bin/.
2. double-click the .exe file "run\_hemodynamic\_model.exe" (in case of linuxOS, execute command "./run\_hemodynamic\_model.exe")
3. the simulation result is generated the directory(circulation\_IsovolumicPhaseAnalysis/simulations/results/hemodynamic\_model/) and it is named "{*YYYYDDDDHHMMss*}.csv".
