## Supplementary material for "A Theoretical Framework for the Hemodynamic Role of Sarcomere Length Dynamics During the Isovolumic Phase of the Left Ventricle": SourceCode: README.md

# Package description
* This package's program is to simulate hemodynamics.
  * To analyze the sarcomere changes during isovolumic phases, two simulation model are contained in this package.
    * volume-based length model : VL model
    * volume-force-coupled length model VFL model 
  
# Folders and Contents

```
📂 RootPackage/
  - README.md : this document
  - 📂 designs/ 
    - "summary_design.md" : Design document for this simulation program
    - "summary_design.html" : html format of design document
  - 📂 documentation/
    - "simulation_guide.md" : Document to explain how to run simulation models
    - "simulation_guide.html" : html format of above document("HowToRun.md")
  - 📂 simulations/
    - 📂 bin/
      - 📂 hemodynamic_model/
        - "run_hemodynamic_model.exe" : binary file for hemodynamic model simulation
    - 📂 build/
      - 📂 hemodynamic_model/
        - "build.bat" :  compile script for windows
        - "build.sh" : compile script for linuxsimulation
    - 📂 configuration/
      - "hemodynamic_model_variables.ini" : configuration file for hemodynamic model
    - 📂 results/
      - 📂 hemodynamic_model/ : the directory to place simulation result
    - 📂 src/ : all source codes are here
      - 📂 calculation/ : utility package for calculation
      - 📂 configuration_reader/ : utility package for reading configuration
      - 📂 csvmaker/ : csv utlity package
      - 📂 logging/ : logging package
      - 📂 main/ : 
        - run_hemodynamic_model.c : main .c file for hemodynamic simulation
      - 📂 models/ 
        - 📂 circulation/ : circulation model based on windkessel 
        - 📂 contraction/ : contraction model based on Negroni Lascano 2008
        - 📂 left_ventricle/ : left ventricular model assumed as sphere shape 
```
