## Supplementary material for "A Theoretical Framework for the Hemodynamic Role of Sarcomere Length Dynamics During the Isovolumic Phase of the Left Ventricle": SourceCode: build.sh

```
#!/bin/bash

SOURCE_DIR="../../src" 
MODELS_DIR="$SOURCE_DIR/models" 
CALC_DIR="$SOURCE_DIR/calculation" 

SOURCE_FILES="$SOURCE_DIR/main/run_hemodynamic_model.c \
$SOURCE_DIR/logging/logging.c \
$CALC_DIR/euler_calculation.c \
$CALC_DIR/numeric_conversion/conversion_to_canine_param.c \
$CALC_DIR/numeric_conversion/scaling_force.c \
$SOURCE_DIR/csvmaker/csvmaker.c \
$SOURCE_DIR/configuration_reader/configuration_reader.c \
$SOURCE_DIR/plot_utility/plot_utility.c \
$CALC_DIR/multi_domain_coupling/length_coupling.c \
$CALC_DIR/multi_domain_coupling/volume_coupling.c \
$MODELS_DIR/contraction/NL08_in_circulation.c \
$MODELS_DIR/left_ventricle/length_by_volume.c \
$MODELS_DIR/left_ventricle/length_by_volume_force.c \
$MODELS_DIR/left_ventricle/laplace.c \
$MODELS_DIR/left_ventricle/wall_thickness_by_rad.c \
$MODELS_DIR/left_ventricle/rad_by_sphere_vol.c \
$MODELS_DIR/left_ventricle/length_by_radius.c \
$MODELS_DIR/left_ventricle/canine_lvf_relation.c \
$MODELS_DIR/circulation/windkessel.c \
$SOURCE_DIR/configuration_printer/configuration_printer.c"


OUTPUT_DIR="../../bin/hemodynamic_model"

OUTPUT_FILE="run_hemodynamic_model"

OUTPUT_PATH="$OUTPUT_DIR/$OUTPUT_FILE"

mkdir -p $OUTPUT_DIR

gcc -O3 -Wall -Wextra -Wpedantic -std=c11 \
-D_POSIX_C_SOURCE=200809L \
-march=native \
-o "$OUTPUT_PATH" $SOURCE_FILES -lm
```
