## Supplementary material for "A Theoretical Framework for the Hemodynamic Role of Sarcomere Length Dynamics During the Isovolumic Phase of the Left Ventricle": SourceCode: build.sh

```
#!/bin/bash

SOURCE_DIR="../../src"
MODELS_DIR="$SOURCE_DIR/models"
CALC_DIR="$SOURCE_DIR/calculation"
SOURCE_FILES="$SOURCE_DIR/main/run_LVF_relation_analysis.c \
$SOURCE_DIR/logging/logging.c \
$SOURCE_DIR/csvmaker/csvmaker.c \
$SOURCE_DIR/configuration_reader/configuration_reader.c \
$SOURCE_DIR/plot_utility/plot_utility.c \
$MODELS_DIR/left_ventricle/length_by_volume_force.c \
$MODELS_DIR/left_ventricle/canine_lvf_relation.c \
$CALC_DIR/numeric_conversion/conversion_to_canine_param.c \
$CALC_DIR/numeric_conversion/scaling_force.c

OUTPUT_DIR="../../bin/LVF_relation_analysis"

OUTPUT_FILE="run_LVF_relation_analysis.exe"

OUTPUT_PATH="$OUTPUT_DIR/$OUTPUT_FILE"

mkdir -p "$OUTPUT_DIR"

gcc -o "$OUTPUT_PATH" -g $SOURCE_FILES -lm
```
