## Supplementary material for "A Theoretical Framework for the Hemodynamic Role of Sarcomere Length Dynamics During the Isovolumic Phase of the Left Ventricle": SourceCode: build.bat

```
rem @echo off

set SOURCE_DIR=..\..\src
set MODELS_DIR=%SOURCE_DIR%\models
set CALC_DIR=%SOURCE_DIR%\calculation
set SOURCE_FILES=%SOURCE_DIR%\main\run_single_cycle_analysis.c ^
%SOURCE_DIR%\logging\logging.c ^
%SOURCE_DIR%\csvmaker\csvmaker.c ^
%SOURCE_DIR%\configuration_reader\configuration_reader.c ^
%SOURCE_DIR%\csvreader\csvreader.c

set OUTPUT_DIR=..\..\bin\single_cycle_analysis

set OUTPUT_FILE=run_single_cycle_analysis.exe

set OUTPUT_PATH=%OUTPUT_DIR%\%OUTPUT_FILE%

mkdir %OUTPUT_DIR%

gcc -o %OUTPUT_PATH% %SOURCE_FILES% -lm

pause
```
