## Supplementary material for "A Theoretical Framework for the Hemodynamic Role of Sarcomere Length Dynamics During the Isovolumic Phase of the Left Ventricle": SourceCode: hemodynamic_model_variables.ini

```
[time section]
duration=100000.0 # [ms]
dt=0.01 # [ms]
cycle=1000.0 # [ms]
[simulation conditions]
model_flag=0
# this flag is enable when analysis_flag = 0 
# 0 : volume-based Length model
# 1 : volume-force-coupled Length model
analysis_flag=0
# 0 : run simulation intended for no model transition during all cycle
# 1 : run simulation intended for analysis of Isovolumic Contraction : ED-transition(model transition at End-Diastole at final cycle)
# 2 : run simulation intended for analysis of Isovolumic Relaxation : ES-transition(model transition at End-Systole at final cycle)
bisection_volume_scope=0.01
bisection_length_scope=0.001
volume_tolerance=1.0E-8
length_tolerance=1.0E-8
max_count_volume_bisection=50
max_count_length_bisection=50
Rout = 1050.0
[scale variables - original variables]
Kva = 0.2317073170733968
Kvb = 6.32926829266959
Kfa = 1.0
Kfb = 1.5
Fini=0.35
Kbs=2.0
Kps=0.25
[simulation initial variables]
# circulation variables
Plv = 10.0 # [mmHg]
Vlv = 128.333333 # [ml]
Pa = 77.933333 # [mmHg*s/ml]
Va = 116.9 # [mmHg*s/ml]
Rpv = 27.0 # [mmHg*s/ml]
Rlo = 2.0 # [mmHg*s/ml]
PE1 = 12.0 # [mmHg*s/ml]
PE2 = 15.0 # [mmHg*s/ml]
CA = 1.5 # [mmHg*s/ml]
# contraction variables
L=1.10 # [um]
Xp=1.094 # [um]
Xw=1.0999 # [um]
hp=0.006 # [um]
hw=0.0001 # [um]
Ca=4.0E-5 # [mM]
TSCa=0.0 # [mM]
TSCap=0.0 # [mM]
TSCaw=0.0 # [mM]
TSCaeff=0.0 # [mM]
TS=0.0 # [mM]
TSp=0.0 # [mM]
TSw=0.0 # [mM]
g=0.0 # [/ms]
gd=0.0 # [/ms]
[csvmaker]
save_interval = 1000.0 # [ms]
save_start_time = 99000.0 # [ms]
output_dir = ../../results/hemodynamic_model/ #[directory]
[plot utility]
realtime_plot = 0 # 0 : no plot with simulation, 1 : plot with simulation
every_fixed_times = 10 # /[ms] plot every time which this variable configures during simulation
[configuration print]
configuration_print = 0
# 0 : not print calculated parameters 
# 1 : print calculated parameters in the last cycle which are used as the next simulation's ini file
```
