## Supplementary material for "A Theoretical Framework for the Hemodynamic Role of Sarcomere Length Dynamics During the Isovolumic Phase of the Left Ventricle": SourceCode: LVF_relation_analysis_variables.ini

```
[parameters]
Kva = 0.2317073170733968
Kvb = 6.32926829266959
Kfa = 1.0
Kfb = 1.5
[csvmaker]
output_dir = ../../results/LVF_relation_analysis/ #[directory]
[plot utility]
realtime_plot = 0 # 0 : no plot with simulation, 1 : plot with simulation
```
