## Supplementary material for "A Theoretical Framework for the Hemodynamic Role of Sarcomere Length Dynamics During the Isovolumic Phase of the Left Ventricle": SourceCode: length_coupling.c

```
#include <stdio.h>
#include <math.h>
#include "../../logging/logging.h"
#include "../../models/contraction/NL08_in_circulation.h"
#include "../../models/left_ventricle/length_by_volume.h"
#include "../../models/left_ventricle/length_by_volume_force.h"

extern char *INFO;
extern char *ERROR;
extern char *WARN;

extern double L;
extern double bisection_length_scope;
extern int max_count_length_bisection;
extern double length_tolerance;

extern int model_flag;

inline double calc_L(double input_L, double lv_volume){
    double output_L;
    
    if(model_flag == 0){
        output_L = calculate_length_from_volume(lv_volume);
    }else{
        double Fb_local = calculate_active_force(input_L);
        double Fp_local = calculate_passive_force(input_L);
        double Fext_local = calculate_total_force(Fb_local, Fp_local);
        output_L = calculate_length_from_volume_force(lv_volume, Fext_local);
    }

    return output_L;
}

int calculate_length_coupling(double t, double lv_volume, double *result_length){

    double left_L = L - bisection_length_scope;
    double right_L = L + bisection_length_scope;
    double L_A = L;
    double L_B = calc_L(L_A, lv_volume);

    int count = 0;
    //while( right_L - left_L >= length_tolerance){
    while( fabs( L_B - L_A ) >= length_tolerance ){

        double sol = (L_B - L_A) * ( calc_L( left_L, lv_volume ) - left_L );

        if( fabs(sol) < 1.0E-18 ){
            break;
        }else if( sol < 0.0 ){
            // shift for left
            right_L = L_A;
        }else{
            // shift for right
            left_L = L_A;
        }
        L_A = ( left_L + right_L ) / 2.0;
        L_B = calc_L(L_A, lv_volume);

        count++;
        if(count > max_count_length_bisection){
            printf("Left L : %.15f\n", left_L);
            printf("L_A : %.15f\n", L_A);
            printf("L_B : %.15f\n", L_B);
            printf("right L : %.15f\n", right_L);
            log_message_with_line(ERROR, "not found length", __FILE__, __LINE__);

            // debug
            printf("L_A, L_B, L_B-L_A\n");
            for(double length_input = L - bisection_length_scope; length_input < L + bisection_length_scope ; length_input = length_input + 0.00001){
                double length_output = calc_L(length_input, lv_volume);
                printf("%.15f,%.15f,%.15f\n", length_input, length_output, length_output-length_input);
            }
            return 1;
        }
    }        

    // finally, L is a solution of this bisection calculation
    *result_length = L_A;

    return 0;
}
```
