## Supplementary material for "A Theoretical Framework for the Hemodynamic Role of Sarcomere Length Dynamics During the Isovolumic Phase of the Left Ventricle": SourceCode: volume_coupling.c

```
#include <stdio.h>
#include <math.h>
#include "../../logging/logging.h"
#include "../../models/contraction/NL08_in_circulation.h"
#include "../../models/circulation/windkessel.h"
#include "../../models/left_ventricle/rad_by_sphere_vol.h"
#include "../../models/left_ventricle/wall_thickness_by_rad.h"
#include "../multi_domain_coupling/length_coupling.h"
#include "../../models/left_ventricle/laplace.h"

extern char *INFO;
extern char *ERROR;
extern char *WARN;

extern double Vlv;
extern double bisection_volume_scope;
extern int max_count_volume_bisection;
extern double volume_tolerance;

extern double L;
extern double Fb;
extern double Fp;
extern double Rlv;
extern double hlv;
extern double F;
extern double Plv;

double cached_input_Vlv = -1.0;
double cached_output_Vlv = -1.0;

int calc_Vlv(double t, double input_Vlv, double *output_Vlv, double *output_L){

    double Rlv_local = calculate_radius_from_volume(input_Vlv);
    double hlv_local = calculate_wall_thickness(Rlv_local);

    if(calculate_length_coupling(t, input_Vlv, output_L) == 1){
        char line[512];
        snprintf(line, sizeof(line) ,"not found length which mathes to volume : time ( %.5f )", t);
        log_message_with_line(ERROR, line, __FILE__, __LINE__);
        return 1;
    }
    double L_local = *output_L;
    double Fb_local = calculate_active_force(L_local);
    double Fp_local = calculate_passive_force(L_local);
    double F_local = calculate_total_force(Fb_local, Fp_local);
    double Plv_local = calculate_pressure(F_local, Rlv_local, hlv_local);

    // set Vlv which is calculated with windkessel
    *output_Vlv = calculate_lv_volume(Plv_local);

    return 0;
}

/**
 * determine values at the specified step
*/
int determine_values(double t, double determined_Vlv, double determined_L){
    
    Rlv = calculate_radius_from_volume(determined_Vlv);
    hlv = calculate_wall_thickness(Rlv);
    L = determined_L;
    Fb = calculate_active_force(L);
    Fp = calculate_passive_force(L);
    F = calculate_total_force(Fb, Fp);
    Plv = calculate_pressure(F, Rlv, hlv);

    calculate_circulation(t);
    
    return 0;

}

int calculate_volume_coupling(double t){

    double left_Vlv = Vlv - bisection_volume_scope;
    double right_Vlv = Vlv + bisection_volume_scope;
    double Vlv_A = Vlv;

    double output_left_Vlv;
    double output_left_Vlv_L;

    // modi
    double output_right_Vlv;
    double output_right_Vlv_L;
    // if(calc_Vlv(t, left_Vlv, &output_left_Vlv, &output_left_Vlv_L)) return 1;
    // if(calc_Vlv(t, right_Vlv, &output_right_Vlv, &output_right_Vlv_L)) return 1;

    double Vlv_B;
    double Vlv_B_length;
    if(calc_Vlv(t, Vlv_A, &Vlv_B, &Vlv_B_length) == 1) return 1;

    int count = 0;
    while( fabs( Vlv_B - Vlv_A ) > volume_tolerance && fabs(right_Vlv - left_Vlv) > volume_tolerance) {
    // while( right_Vlv - left_Vlv >= volume_tolerance ){
    // while( fabs( Vlv_B - Vlv_A ) > volume_tolerance ) {
    // while( fabs(right_Vlv - left_Vlv) > volume_tolerance) {
       
        if( calc_Vlv(t, left_Vlv, &output_left_Vlv, &output_left_Vlv_L) == 1 ) return 1; // error handling
        
        double sol = ( Vlv_B - Vlv_A ) * ( output_left_Vlv - left_Vlv );
        if( fabs(sol) < 1.0E-18 ){
            break;
        }
        else if( sol < 0.0 ){
            // shift for left
            right_Vlv = Vlv_A;
        }else{
            // shift for right
            left_Vlv = Vlv_A;
        }
        Vlv_A = ( left_Vlv + right_Vlv ) / 2.0;
        if(calc_Vlv(t, Vlv_A, &Vlv_B, &Vlv_B_length) == 1) return 1;
        
        count++;
        if(count > max_count_volume_bisection){
            log_message_with_line(ERROR, "not found resolved value by bisection method limit", __FILE__, __LINE__);
            printf("left Vlv : %.15f\n", left_Vlv);
            printf("Vlv_A : %.15f\n", Vlv_A);
            printf("Vlv_B : %.15f\n", Vlv_B);
            printf("Vlv_B_L : %.15f\n", Vlv_B_length);
            printf("right Vlv : %.15f\n", right_Vlv);
            printf("t : %f\n", t);
            return 1;
        }
    }    
    
    return determine_values(t, Vlv_B, Vlv_B_length);
}
```
