## Supplementary material for "A Theoretical Framework for the Hemodynamic Role of Sarcomere Length Dynamics During the Isovolumic Phase of the Left Ventricle": SourceCode: volume_coupling.h

```
#ifndef VOLUME_COUPLING_H
#define VOLUME_COUPLING_H

double calc_Vlv(double input_Vlv);

int determine_values(double t, double determined_Vlv, double determined_L);

int calculate_volume_coupling(double t);

#endif
```
