## Supplementary material for "A Theoretical Framework for the Hemodynamic Role of Sarcomere Length Dynamics During the Isovolumic Phase of the Left Ventricle": SourceCode: conversion_to_canine_param.c

```
#include<stdio.h>
#include<math.h>
#include "../../logging/logging.h"
#include "../euler_calculation.h"

extern double Kva;
extern double Kvb;
extern double Kfa;
extern double Kfb;

double convert_to_canine_lv_volume(double human_lv_volume){
    //return (human_lv_volume - 59.0) / 4.31578947368 + 20.0;
    return ( Kva * human_lv_volume + Kvb );
}

double convert_to_canine_lv_Fext(double Ks, double human_lv_force){
    
    return (Kfa * human_lv_force + Kfb);
}
```
