## Supplementary material for "A Theoretical Framework for the Hemodynamic Role of Sarcomere Length Dynamics During the Isovolumic Phase of the Left Ventricle": SourceCode: conversion_to_canine_param.h

```
#ifndef CONVERSION_TO_CANINE_PARAM_H
#define CONVERSION_TO_CANINE_PARAM_H

double convert_to_canine_lv_volume(double human_lv_volume);

double convert_to_canine_lv_Fext(double Ks, double human_lv_force);

#endif
```
