## Supplementary material for "A Theoretical Framework for the Hemodynamic Role of Sarcomere Length Dynamics During the Isovolumic Phase of the Left Ventricle": SourceCode: scaling_force.c

```
#include<stdio.h>
#include<math.h>
#include "../../logging/logging.h"

double get_force_scale_param(int model_flag, int mode_change_flag){

    double Ks;
    if( model_flag == 0 && mode_change_flag == 0 ){
        // ★どうやってスケールしたか謎
        // Ks = 3.78174; // original 
        Ks = 3.790;
    } else {
        // ★どうやってスケールしたか謎
        //Ks = 3.6177425; // original
        Ks = 3.6195;
        //Ks = 3.2; // trial
    }
    return Ks;
}
```
