## Supplementary material for "A Theoretical Framework for the Hemodynamic Role of Sarcomere Length Dynamics During the Isovolumic Phase of the Left Ventricle": SourceCode: configuration_printer.c

```
#include<stdio.h>
#include<string.h>
#include<stdlib.h>
#include "../logging/logging.h"
#include<time.h>
#include<stdarg.h>

// logging global variables
extern const char *ERROR;
extern const char *WARN;
extern const char *INFO;

extern double Plv;
extern double Vlv;
extern double Pa;
extern double Va;
extern double Rpv;
extern double Rlo;
extern double PE1;
extern double PE2;
extern double CA;
extern double L;
extern double Xp;
extern double Xw;
extern double hp;
extern double hw;
extern double Ca;
extern double TSCa;
extern double TSCap;
extern double TSCaw;
extern double TSCaeff;
extern double TS;
extern double TSp;
extern double TSw;
extern double g;
extern double gd;

extern int configuration_print;

int printout_config(const char *output_dir){

    if(configuration_print == 0){
        return 0;
    }

    log_message(INFO, "start printing configuration parameters");

    FILE *file;
    time_t rawtime;
    struct tm * timeinfo;
    char timestamp[20];
    char file_path[256];

    time(&rawtime);
    timeinfo = localtime(&rawtime);

    strftime(timestamp,sizeof(timestamp), "%Y%m%d%H%M%S", timeinfo);

    snprintf(file_path, sizeof(file_path), "%s%s%s", output_dir, timestamp, ".ini");

    file = fopen(file_path, "w");
    if(file == NULL){
        log_message(ERROR, "failed to open ini file");
        return 1;
    }
    log_message(INFO, "opened ini file");

    fprintf(file, "%s = %f\n", "Plv", Plv);
    fprintf(file, "%s = %f\n", "Vlv", Vlv);
    fprintf(file, "%s = %f\n", "Pa", Pa);
    fprintf(file, "%s = %f\n", "Va", Va);
    fprintf(file, "%s = %f\n", "Rpv", Rpv);
    fprintf(file, "%s = %f\n", "Rlo", Rlo);
    fprintf(file, "%s = %f\n", "PE1", PE1);
    fprintf(file, "%s = %f\n", "PE2", PE2);
    fprintf(file, "%s = %f\n", "CA", CA);
    fprintf(file, "%s = %f\n", "L", L);
    fprintf(file, "%s = %f\n", "Xp", Xp);
    fprintf(file, "%s = %f\n", "Xw", Xw);
    fprintf(file, "%s = %f\n", "hp", hp);
    fprintf(file, "%s = %f\n", "hw", hw);
    fprintf(file, "%s = %f\n", "Ca", Ca);
    fprintf(file, "%s = %f\n", "TSCa", TSCa);
    fprintf(file, "%s = %f\n", "TSCap", TSCap);
    fprintf(file, "%s = %f\n", "TSCaw", TSCaw);
    fprintf(file, "%s = %f\n", "TSCaeff", TSCaeff);
    fprintf(file, "%s = %f\n", "TS", TS);
    fprintf(file, "%s = %f\n", "TSp", TSp);
    fprintf(file, "%s = %f\n", "TSw", TSw);
    fprintf(file, "%s = %f\n", "g", g);
    fprintf(file, "%s = %f\n", "gd", gd);

    log_message(INFO, "finishied writing ini file, successfully");

    fclose(file);
    log_message(INFO, "closed ini file, successfully");

    return 0;

}
```
