## Supplementary material for "A Theoretical Framework for the Hemodynamic Role of Sarcomere Length Dynamics During the Isovolumic Phase of the Left Ventricle": SourceCode: configuration_reader.c

```
#include<stdio.h>
#include<string.h>
#include<stdlib.h>
#include "configuration_reader.h"
#include "../logging/logging.h"

// logging global variables
extern const char *ERROR;
extern const char *WARN;
extern const char *INFO;

// key and value in ini file
extern char key[128];
extern char value_str[128];

int open_ini(FILE **file, const char *conf_path){
    *file = fopen(conf_path, "r");
    if(*file == NULL){
        log_message(ERROR, "failed to open configuration file");
        return 0;
    }
    log_message(INFO, "opened configuration file");
    return 1;
}

int set_double_var(const char *expected_key, double *val){
    if(strcmp(key, expected_key) == 0){
        // set value because of correspond
        if(sscanf(value_str, "%lf", val) != 1) {
            log_message(ERROR, "unexpected variable in ini file");
            return -1;
        }
        //printf("%f\n", *val); // for test
        return 1;
    }else{
        // not set value because of different key
        return 0;
    }
}

int set_char_var(const char *expected_key, char *val){
    if(strcmp(key, expected_key) == 0){
        // set value because of correspond
        strcpy(val, value_str);
        return 1;
    }else{
        // not set value because of different key
        return 0;
    }
}

int set_int_var(const char *expected_key, int *val){
    if(strcmp(key, expected_key) == 0){
        // set value because of correspond
        if(sscanf(value_str, "%d", val) != 1) {
            log_message(ERROR, "unexpected variable in ini file");
            return -1;
        }
        return 1;
    }else{
        // not set value because of different key
        return 0;
    }
}

void close_ini(FILE **file){
    fclose(*file);
    log_message(INFO, "closed configuration file");
}
```
