## Supplementary material for "A Theoretical Framework for the Hemodynamic Role of Sarcomere Length Dynamics During the Isovolumic Phase of the Left Ventricle": SourceCode: configuration_reader.h

```
#ifndef CONFIGURATION_READER_H
#define CONFIGURATION_READER_H

int open_ini(FILE **file, const char *conf_path);

int set_double_var(const char *expected_key, double *val);

int set_char_var(const char *expected_key, char *val);

int set_int_var(const char *expected_key, int *val);

void close_ini(FILE **file);

#endif
```
