## Supplementary material for "A Theoretical Framework for the Hemodynamic Role of Sarcomere Length Dynamics During the Isovolumic Phase of the Left Ventricle": SourceCode: csvmaker.c

```
#include<stdio.h>
#include<string.h>
#include<stdlib.h>
#include "../logging/logging.h"
#include<time.h>
#include<stdarg.h>

// logging global variables
extern const char *ERROR;
extern const char *WARN;
extern const char *INFO;


int create_csv(FILE **csv_file, const char *output_dir){

    time_t rawtime;
    struct tm *timeinfo;
    char timestamp[20];
    char file_path[256];

    time(&rawtime);
    timeinfo = localtime(&rawtime);

    strftime(timestamp,sizeof(timestamp), "%Y%m%d%H%M%S", timeinfo);

    snprintf(file_path, sizeof(file_path), "%s%s%s", output_dir, timestamp, ".csv");

    *csv_file = fopen(file_path, "w");
    if(*csv_file == NULL){
        log_message(ERROR, "failed to open csv file");
        return 0;
    }
    log_message(INFO, "opened csv file");

    return 1;
}

void close_csv(FILE **csv_file){
    fclose(*csv_file);
    log_message(INFO, "closed csv file, successfully");
}

void print_var_csv(int num, FILE **csv_file, ...){
    va_list args;

    va_start(args, *csv_file);

    for(int i = 0 ; i< num ; i++){
        double value = va_arg(args, double);
        if(i == 0){
            fprintf(*csv_file, "%.15f" , value);
            //fprintf(*csv_file, "%.5f", value); // plot time
        }else{
            fprintf(*csv_file, ",%.15f" , value);
        }
        
    }
    fprintf(*csv_file, "\n");

    va_end(args);
}

void print_char_csv(int num, FILE **csv_file, ...){  
    va_list args;

    va_start(args, *csv_file);

    for(int i = 0 ; i< num ; i++){
        //char *value = va_arg(args, char*);
        char value[128] = "";
        strcpy(value, va_arg(args, char*));
        if(i == 0){
            fprintf(*csv_file, "%s" , value);
        }else{
            fprintf(*csv_file, ",%s" , value);
        }
        
    }
    fprintf(*csv_file, "\n");

    va_end(args);
}
```
