## Supplementary material for "A Theoretical Framework for the Hemodynamic Role of Sarcomere Length Dynamics During the Isovolumic Phase of the Left Ventricle": SourceCode: csvmaker.h

```
#ifndef CSVMAKER_H
#define CSVMAKER_H
#include<stdio.h>

int create_csv(FILE **csv_file, const char *output_dir);

void close_csv(FILE **csv_file);

void print_char_csv(int num, FILE **csv_file, ...);

void print_var_csv(int num, FILE **csv_file, ...);

#endif
```
