## Supplementary material for "A Theoretical Framework for the Hemodynamic Role of Sarcomere Length Dynamics During the Isovolumic Phase of the Left Ventricle": SourceCode: csvreader.c

```
#include<stdio.h>
#include<string.h>
#include<stdlib.h>
#include "../logging/logging.h"

// logging global variables
extern const char *ERROR;
extern const char *WARN;
extern const char *INFO;

FILE *get_csv_pointer(char *csv_file_path){
    FILE *fp = fopen(csv_file_path, "r");
    if(fp == NULL){
        log_message(ERROR, "failed to open csv file");
        return NULL;
    }
    return fp;
}

void release_csv(FILE *fp){
    fclose(fp);
}


int read_csv(FILE *fp, double **data, int *rows_num, int *columns_num){

    char line[1024];
    int rows = 0, cols = 0;
    while(fgets(line, 1024, fp) != NULL){
        char *token = strtok(line, ",");
        int col = 0;
        while(token != NULL){
            token = strtok(NULL, ",");
            col++;
        }
        if(col > cols) cols = col; 
        rows++;
    }
    rewind(fp);

    *data = (double *)malloc(rows * cols * sizeof(double));
    int row = 0;
    while (fgets(line, 1024, fp) != NULL) {
        char *token = strtok(line, ",");
        int col = 0;
        while (token != NULL) {
            (*data)[row * cols + col] = atof(token);
            token = strtok(NULL, ",");
            col++;
        }
        row++;
    }

    rewind(fp);

    *rows_num = rows;
    *columns_num = cols;

    return 0;
}

int get_column_num(FILE *fp, const char *param){
    int ret_num = -1;

    char line[1024];
    if (fgets(line, sizeof(line), fp) != NULL) {
        char *token = strtok(line, ",");
        int col = 0;
        while (token != NULL) {
            col++;
            if (strcmp(token, param) == 0) {
                ret_num = col-1;
                break;
            }
            // printf("%s\n",token);
            token = strtok(NULL, ",");
        }
    }

    rewind(fp);  // return file pointer to the beginning

    return ret_num;
}
```
