## Supplementary material for "A Theoretical Framework for the Hemodynamic Role of Sarcomere Length Dynamics During the Isovolumic Phase of the Left Ventricle": SourceCode: csvreader.h

```
#ifndef CSVREADER_H
#define CSVREADER_H
#include<stdio.h>

FILE *get_csv_pointer(char *csv_file_path);

void release_csv(FILE *fp);

int read_csv(FILE *fp, double **data, int *rows_num, int *columns_num);

int get_column_num(FILE *fp, const char *param);

#endif
```
