## Supplementary material for "A Theoretical Framework for the Hemodynamic Role of Sarcomere Length Dynamics During the Isovolumic Phase of the Left Ventricle": SourceCode: logging.c

```
#include<stdio.h>
#include<string.h>
#include<time.h>
#include "logging.h"

extern double cycle;

void log_message(const char *log_level, const char *message){
    printf("[%s] %s \n", log_level, message);
}

void log_message_with_line(const char *log_level, const char *message, const char *file, int line){
    printf("[%s] %s \n", log_level, message);
    printf("[%s] line : %d \n", file, line);
}

void log_progress( double t ){
    char line[256];
    time_t rawtime;
    struct tm* timeinfo;
    char buffer[20];
    time(&rawtime);
    timeinfo = localtime(&rawtime);
    strftime(buffer, sizeof(buffer), "%Y%m%d%H%M%S", timeinfo);
    snprintf(line, sizeof(line), "%s : cycle num ( %d )", buffer, ( (int)(t / cycle) + 1 ) );
    log_message("INFO", line);
}
```
