## Supplementary material for "A Theoretical Framework for the Hemodynamic Role of Sarcomere Length Dynamics During the Isovolumic Phase of the Left Ventricle": SourceCode: logging.h

```
#ifndef LOGGING_H
#define LOGGING_H

void log_message(const char *log_level, const char *message);

void log_message_with_line(const char *log_level, const char *message, const char *file, int line);

void log_progress(double t);

#endif
```
