## Supplementary material for "A Theoretical Framework for the Hemodynamic Role of Sarcomere Length Dynamics During the Isovolumic Phase of the Left Ventricle": SourceCode: run_hemodynamic_model.c

```
#include<stdio.h>
#include<string.h>
#include<stdlib.h>
#include<time.h>
#include<math.h>
#include<stdarg.h>
#ifdef _WIN32
#include <direct.h>
#define GET_CURRENT_DIR _getcwd
#else
#include <unistd.h>
#define GET_CURRENT_DIR getcwd
#endif

#include "../logging/logging.h"
#include "../calculation/euler_calculation.h"
#include "../csvmaker/csvmaker.h"
#include "../configuration_reader/configuration_reader.h"
#include "../plot_utility/plot_utility.h"
#include "../calculation/multi_domain_coupling/volume_coupling.h"
#include "../calculation/multi_domain_coupling/length_coupling.h"
#include "../models/contraction/NL08_in_circulation.h"
#include "../models/left_ventricle/rad_by_sphere_vol.h"
#include "../models/left_ventricle/laplace.h"
#include "../models/left_ventricle/length_by_radius.h"
#include "../models/circulation/windkessel.h"
#include "../models/left_ventricle/wall_thickness_by_rad.h"
#include "../configuration_printer/configuration_printer.h"
#include "../models/left_ventricle/length_by_volume_force.h"

// configuration file path (mandatory)
const char *conf_path = "../../configuration/hemodynamic_model_variables.ini";
FILE *file = NULL;

// time variables
double duration;
double dt;
double cycle;

// simulation settings
int model_flag;
int analysis_flag;
double bisection_volume_scope;
int max_count_volume_bisection;
double volume_tolerance;
double bisection_length_scope;
int max_count_length_bisection;
double length_tolerance;
double Rout;
int mode_change_flag = 0;
double Kva;
double Kvb;
double Kfa;
double Kfb;
double Fini;
double Kbs;
double Kps;

// parameters
//// circulation
double Plv;
double Vlv;
double Pa;
double Va;
double Rpv;
double Rlo;
double PE1;
double PE2;
double CA;

//// contraction
double L;
double Xp;
double Xw;
double hp;
double hw;
double Ca;
double TSCa;
double TSCap;
double TSCaw;
double TSCaeff;
double TS;
double TSp;
double TSw;
double g;
double gd;

double Fb;
double Fp;
double F;

// windkessel.c : internal use 
int q1_valve_count = 0;
int qin_valve_count = 0;
double q0 = 0.0; // [ml/ms]
double q1 = 0.0; // =dVlvdt [ml/ms]
double q2 = 0.0; // =dVadt [ml/ms]
double qin = 0.0; // [ml/ms]
double qout = 0.0; // [ml/ms]
double dPadt = 0.0; // [mmHg/ms]

//// left ventricle physics
double Rlv;
double hlv;

// NL08_chem.c : internal use
double dTSCadt = 0.0;
double dTSCawdt = 0.0;
double dTSCapdt = 0.0;
double dTSpdt = 0.0;
double dTSwdt = 0.0;

double Itropn = 0.0;
double Qpump = 0.0;
double Qrel = 0.0;
double Qsr = 0.0;
double dCadt = 0.0;
double dXpdt = 0.0;
double dXwdt = 0.0;

// make csv file
double save_interval;
double save_start_time;
char output_dir[128];

// plot utility
int realtime_plot;
int every_fixed_times;

// configuration print
int configuration_print;

// return pattern
int success_exit = 0;
int error_exit = 1;
int simulation_error_exit = -1;

// logging global variables
const char *ERROR = "ERROR";
const char *WARN = "WARN";
const char *INFO = "INFO";

// key and value in ini file
char key[128];
char value_str[128];

// scan and set initial values from ini file
int scan_initial_values(){
    ///// configuration check
    log_message(INFO, "start configuration check");
    if ( open_ini(&file, conf_path) == 0) return error_exit;
    char line[256] = "";
    char section[256] = "";

    log_message(INFO, "Here's initial values : ");
    while (fgets(line, sizeof(line), file)) {
        // cut comment-out
        if (line[0] == ';' || line[0] == '#') {
            continue;
        }
        // cut section name
        if (line[0] == '[' && line[strlen(line) - 2] == ']') {
            line[strlen(line) - 2] = '\0';
            strcpy(section, line + 1);
            continue;
        }
        // set variables which obtained from ini file to global variable
        if (sscanf(line, "%127[^= ] = %127s", key, value_str) == 2) {
            printf("%s", line);
            if (set_double_var("duration", &duration) == -1
                || set_double_var("dt", &dt) == -1
                || set_double_var("cycle", &cycle) == -1
                || set_int_var("model_flag", &model_flag) == -1
                || set_int_var("analysis_flag", &analysis_flag) == -1
                || set_double_var("bisection_volume_scope", &bisection_volume_scope) == -1
                || set_int_var("max_count_volume_bisection", &max_count_volume_bisection) == -1
                || set_double_var("volume_tolerance", &volume_tolerance) == -1
                || set_double_var("bisection_length_scope", &bisection_length_scope) == -1
                || set_int_var("max_count_length_bisection", &max_count_length_bisection) == -1
                || set_double_var("length_tolerance", &length_tolerance) == -1
                || set_double_var("Rout", &Rout) == -1
                || set_double_var("Plv", &Plv) == -1
                || set_double_var("Vlv", &Vlv) == -1
                || set_double_var("Pa", &Pa) == -1
                || set_double_var("Va", &Va) == -1
                || set_double_var("Rpv", &Rpv) == -1
                || set_double_var("Rlo", &Rlo) == -1
                || set_double_var("PE1", &PE1) == -1
                || set_double_var("PE2", &PE2) == -1
                || set_double_var("CA", &CA) == -1
                || set_double_var("L", &L) == -1
                || set_double_var("Xp", &Xp) == -1
                || set_double_var("Xw", &Xw) == -1
                || set_double_var("hp", &hp) == -1
                || set_double_var("hw", &hw) == -1
                || set_double_var("Ca", &Ca) == -1
                || set_double_var("TSCa", &TSCa) == -1
                || set_double_var("TSCap", &TSCap) == -1
                || set_double_var("TSCaw", &TSCaw) == -1
                || set_double_var("TSCaeff", &TSCaeff) == -1
                || set_double_var("TS", &TS) == -1
                || set_double_var("TSp", &TSp) == -1
                || set_double_var("TSw", &TSw) == -1
                || set_double_var("g", &g) == -1
                || set_double_var("gd", &gd) == -1
                || set_double_var("Kva", &Kva) == -1
                || set_double_var("Kvb", &Kvb) == -1
                || set_double_var("Kfa", &Kfa) == -1
                || set_double_var("Kfb", &Kfb) == -1
                || set_double_var("Fini", &Fini) == -1
                || set_double_var("Kbs", &Kbs) == -1
                || set_double_var("Kps", &Kps) == -1
                || set_double_var("save_interval", &save_interval) == -1
                || set_double_var("save_start_time", &save_start_time) == -1
                || set_char_var("output_dir", output_dir) == -1
                || set_int_var("realtime_plot", &realtime_plot) == -1
                || set_int_var("every_fixed_times", &every_fixed_times) == -1
                || set_int_var("configuration_print", &configuration_print) == -1
            ) return error_exit;
        }
    }
    if(analysis_flag != 0){
        model_flag = 1;
    }
    close_ini(&file);
    log_message(INFO, "end configuration check");
    return success_exit;
}

int exit_process(int code){
    switch(code){
        case 0 :
            log_message(INFO, "finished simulation, successfully");
            break;
        case 1:
            log_message(ERROR, "failed to finish simulation");
            break;
        case -1:
            log_message(ERROR, "stopped simulation...");
            break;
        default:
            break;
    }
    log_message(INFO, "terminating...");
    clock_t start = clock();
    while ((clock() - start) * 1000 / CLOCKS_PER_SEC < 20000);
    if(code == 0){
        return code;
    }else{
        return 1;
    }
}

int find_current_dir(){
    char cwd[512];
    if (GET_CURRENT_DIR(cwd, sizeof(cwd)) != NULL) {
        char line[256];
        snprintf(line, sizeof(line), "current directory : %s", cwd);
        log_message(INFO, line);
        return 0;
    } else {
        char line[256]="failed getcwd()";
        log_message(ERROR, line);
        return 1;
    }
}

int find_nan(int num, ...){
    int ret = 0;
    va_list args;
    va_start(args, num);
    for( int i = 0 ; i < num ; i++ ){
        double val = va_arg(args, double);
        if( isnan( val ) ){
            ret = 1;
            break;
        }
    }
    va_end(args);
    return ret;
}

int set_initial_values( double t ){
    // set initial L from initial Vlv
    Rlv = calculate_radius_from_volume(Vlv);
    if( model_flag == 0 ){
        L = calculate_length_from_radius(Rlv);
    }else if( model_flag == 1){
        L = calculate_length_from_volume_force(Vlv, Fini);
    }else{
        log_message(ERROR, "failed to set initial value : illegal model_flag");
        return error_exit;
    }
    char line[256];
    snprintf(line, sizeof(line), "initial Vlv [%f], initial Rlv [%f], initial L [%f]", Vlv, Rlv, L );
    log_message(INFO, line);
    return success_exit;
}

void change_mode_flag( double t, double prev_q1 ){
    char line[256];
    if(analysis_flag == 1 && mode_change_flag == 0 && q0 == 0.0 && qin == 0.0 && Pa > Plv){
        // change model_flag to 0 when t > tED
        model_flag = 0;
        mode_change_flag = 1;
        snprintf(line, sizeof(line), "changed from VFL model to VL model : t = %f", t);
        log_message(INFO, line);
    }
    else if(analysis_flag == 2 && mode_change_flag == 0 && q0 == 0.0 && qin == 0.0 && prev_q1 < 0 && q1 == 0.0){
        // change model_flag to 0 when t > tES
        model_flag = 0;
        mode_change_flag = 1;
        snprintf(line, sizeof(line), "changed from VFL model to VL model : t = %f", t);
        log_message(INFO, line);
    }
}

int main(void){

    if(find_current_dir() == error_exit) return exit_process(error_exit);
    if(scan_initial_values() == error_exit) return exit_process(error_exit);

    log_message(INFO, "start simulation...");
    FILE *csv_file = NULL, *gp_pressure = NULL, *gp_volume = NULL;
    if( create_csv(&csv_file, output_dir) == 0 ) return exit_process(error_exit);

    // -------- initialization plot begin --------
    //initialize_plot_process(&gp_pressure, 0.0, 1000.0, "time[ms]", -10.0, 140.0, "pressure[mmHg]");
    //initialize_plot_process(&gp_volume, 0.0, 1000.0, "time[ms]", -10.0, 140.0, "volume[mL]");
    // -------- initialization plot terminate --------

    // write out variables name in the top of sheet
    log_message(INFO, "writing simulation results in csv file");
    print_char_csv(33, &csv_file, "t", "Plv", "Vlv", "hlv", "Rlv", "L", "F", "Fb", "Fp", "Pa", "Va", "q0", "q1", "qin" , "q2", "qout", "hp", "hw", "TSCap" , "TSp", "TSCaw", "TSw", "TSCa", "TS", "dTSCapdt", "dTSCawdt", "dTSpdt", "dTSwdt", "dTSCadt", "Ca", "Itropn", "Qrel", "Qpump");

    double prev_q1 = 0.0, t = 0.0, save_end_time = save_start_time + save_interval;
    for( ; t < duration ; t = t + dt){
        double t_cycle = fmod( t, cycle);
        // -------- define plot section begin --------
        //define_pointtype_clear_prev_cycle(&gp_pressure, "7", t_cycle);
        //define_pointtype_clear_prev_cycle(&gp_volume, "8", t_cycle);
        // -------- define plot section terminate --------

        // -------- calculation section begin --------
        if(t < dt){
            if( set_initial_values(t) == error_exit )
                return exit_process(error_exit);
        }
        if(t_cycle < dt){
            log_progress(t);
        }
        prev_q1 = q1;
        if(calculate_volume_coupling(t_cycle)  == 1) return exit_process(simulation_error_exit);
        calculate_cellular_chem_state(t_cycle);
        // -------- calculation section terminate --------

        // -------- plot section begin ---------
        //plot_with_multipoints_every_fixed_time(&gp_pressure, t_cycle, t_cycle, 2, Plv, Pa);
        //plot_with_multipoints_every_fixed_time(&gp_volume, t_cycle, t_cycle, 2, Vlv, Va);
        //output_buffers_at_cycle_end(&gp_pressure, t_cycle);
        //output_buffers_at_cycle_end(&gp_volume, t_cycle);
        // -------- plot section terminate ---------

        if( (t > duration - cycle - dt) && analysis_flag != 0 ){
            change_mode_flag(t, prev_q1);
        }

        // write variables in csv file
        if(t > (save_start_time - dt) && t < (save_end_time + dt)) 
            print_var_csv(33, &csv_file, t, Plv, Vlv, hlv, Rlv, L, F, Fb, Fp, Pa, Va, q0, q1, qin, q2, qout, hp, hw, TSCap, TSp, TSCaw, TSw, TSCa, TS, dTSCapdt, dTSCawdt, dTSpdt, dTSwdt, dTSCadt,Ca, Itropn, Qrel, Qpump );
        // check NaN value during simulation
        /*if( find_nan(33, t, Plv, Vlv, hlv, Rlv, L, F, Fb, Fp, Pa, Va, q0, q1, qin, q2, qout, hp, hw, TSCap, TSp, TSCaw, TSw, TSCa, TS, dTSCapdt, dTSCawdt, dTSpdt, dTSwdt, dTSCadt,Ca, Itropn, Qrel, Qpump) == 1 ){
            char line[256];
            snprintf(line, sizeof(line), "found Nan in any variables : t = %f", t);
            log_message_with_line(ERROR, line, __FILE__, __LINE__);
            break;
        }*/
    }

    terminate_plot_process(&gp_pressure);
    terminate_plot_process(&gp_volume);
    close_csv(&csv_file);
    printout_config(&output_dir);

    return exit_process(success_exit);
}
```
