## Supplementary material for "A Theoretical Framework for the Hemodynamic Role of Sarcomere Length Dynamics During the Isovolumic Phase of the Left Ventricle": SourceCode: run_LVF_relation_analysis.c

```
#include<stdio.h>
#include<string.h>
#include<stdlib.h>
#include<time.h>
#include<math.h>
#include "../csvmaker/csvmaker.h"
#include "../logging/logging.h"
#include "../configuration_reader/configuration_reader.h"
#include "../plot_utility/plot_utility.h"
#include "../models/left_ventricle/length_by_volume_force.h"
#include "../models/left_ventricle/canine_lvf_relation.h"
#include "../calculation/numeric_conversion/conversion_to_canine_param.h"
#include "../models/left_ventricle/rad_by_sphere_vol.h"
#include "../models/left_ventricle/length_by_radius.h"

double dt = 0.01;
double cycle = 1000.0;
double every_fixed_times = 10.0;

double Kva;
double Kvb;
double Kfa;
double Kfb;

const char *conf_path = "../../configuration/LVF_relation_analysis_variables.ini";
FILE *file = NULL;
char output_dir[128];
int realtime_plot;

// key and value in ini file
char key[128];
char value_str[128];

// return pattern
int success_exit = 0;
int error_exit = 1;
int simulation_error_exit = -1;

// logging global variables
const char *ERROR = "ERROR";
const char *WARN = "WARN";
const char *INFO = "INFO";

// dummy parameters
const int model_flag = 1;
const int mode_change_flag = 0; 

int scan_initial_values(){

    ///// configuration check
    log_message(INFO, "start configuration check");

    if ( open_ini(&file, conf_path) == 0) return error_exit;
    char line[256] = "";
    char section[256] = "";

    while (fgets(line, sizeof(line), file)) {
        // cut comment-out
        if (line[0] == ';' || line[0] == '#') {
            continue;
        }
        // cut section name
        if (line[0] == '[' && line[strlen(line) - 2] == ']') {
            line[strlen(line) - 2] = '\0';
            strcpy(section, line + 1);
            continue;
        }

        // set variables which obtained from ini file to global variable
        if (sscanf(line, "%127[^= ] = %127s", key, value_str) == 2) {
            if (set_char_var("output_dir", output_dir) == -1 ) return error_exit;
            if (set_int_var("realtime_plot", &realtime_plot) == -1) return error_exit;
            if (set_double_var("Kva", &Kva) == -1) return error_exit;
            if (set_double_var("Kvb", &Kvb) == -1) return error_exit;
            if (set_double_var("Kfa", &Kfa) == -1) return error_exit;
            if (set_double_var("Kfb", &Kfb) == -1) return error_exit;
        }
    }
    close_ini(&file);
    log_message(INFO, "end configuration check");

    return success_exit;
}

int exit_process(int code){
    if(code == 0){
        log_message(INFO, "finished simulation, successfully");
    } 
    else if(code == 1){
        log_message(ERROR, "failed to finish simulation");
    }else if(code == -1){
        log_message(ERROR, "stopped simulation...");
    }
    log_message(INFO, "terminating...");
    
    clock_t start = clock();
    while ((clock() - start) * 1000 / CLOCKS_PER_SEC < 3000);
    if(code == 0){
        return code;
    }else{
        return 1;
    }
}

int main(void){

    if(scan_initial_values() == error_exit ) return exit_process(error_exit);

    FILE *csv_file = NULL;
    FILE *gp = NULL;

    if( create_csv(&csv_file, output_dir) == 0 ) return exit_process(error_exit);

    initialize_plot_process(&gp, 40.0, 140.0, "Vlv[mL]", 0.8, 1.2, "L[um]");

    print_char_csv(8, &csv_file, "Vlv", "F=-0.5", "F=0.0", "F=1.0", "F=2.0", "F=3.0", "F=4.0", "VL"); 
    double L1 = 0.0;
    double L2 = 0.0;
    double L3 = 0.0;
    double L4 = 0.0;
    double L5 = 0.0;
    double L6 = 0.0;
    double L7 = 0.0;
    // double scaled_F1 = convert_to_canine_lv_Fext(0.0, -0.5);
    // double scaled_F2 = convert_to_canine_lv_Fext(0.0, 1.75);
    // double scaled_F3 = convert_to_canine_lv_Fext(0.0, 2.0);
    // double scaled_F4 = convert_to_canine_lv_Fext(0.0, 3.25);
    // double scaled_F5 = convert_to_canine_lv_Fext(0.0, 4.5);

    for(double Vlv = 40.0; Vlv < 140.0 ; Vlv = Vlv + 0.1){
        if(Vlv == 0.0){
            define_pointtype_clear_prev_cycle(&gp, "7", 0.0);
        }
        //double scailed_volume = convert_to_canine_lv_volume(Vlv);

        L1 = calculate_length_from_volume_force(Vlv, -0.5);
        L2 = calculate_length_from_volume_force(Vlv, 0.0);
        L3 = calculate_length_from_volume_force(Vlv, 1.0);
        L4 = calculate_length_from_volume_force(Vlv, 2.0);
        L5 = calculate_length_from_volume_force(Vlv, 3.0);
        L6 = calculate_length_from_volume_force(Vlv, 4.0);
        L7 = calculate_length_from_radius(calculate_radius_from_volume(Vlv));
        
        plot_with_multipoints(&gp, Vlv, 7, L1,L2,L3,L4,L5,L6,L7);

        print_var_csv(8, &csv_file, Vlv, L1, L2,L3,L4,L5,L6,L7);
    }

    output_buffers_at_cycle_end(&gp, 1000);
    terminate_plot_process(&gp);
    close_csv(&csv_file);

    return exit_process(success_exit);
}
```
