## Supplementary material for "A Theoretical Framework for the Hemodynamic Role of Sarcomere Length Dynamics During the Isovolumic Phase of the Left Ventricle": SourceCode: run_single_cycle_analysis.c

```
#include<stdio.h>
#include<string.h>
#include<stdlib.h>
#include<time.h>
#include<math.h>
#include "../csvmaker/csvmaker.h"
#include "../csvreader/csvreader.h"
#include "../logging/logging.h"
#include "../configuration_reader/configuration_reader.h"

const char *conf_path = "../../configuration/single_cycle_analysis.ini";
FILE *file = NULL;
char output_dir[128];
char input_file_path[512];

// key and value in ini file
char key[128];
char value_str[128];

// return pattern
int success_exit = 0;
int error_exit = 1;
int simulation_error_exit = -1;

// logging global variables
const char *ERROR = "ERROR";
const char *WARN = "WARN";
const char *INFO = "INFO";

int scan_initial_values(){

    ///// configuration check
    log_message(INFO, "start configuration check");

    if ( open_ini(&file, conf_path) == 0) return error_exit;
    char line[256] = "";
    char section[256] = "";

    while (fgets(line, sizeof(line), file)) {
        // cut comment-out
        if (line[0] == ';' || line[0] == '#') {
            continue;
        }
        // cut section name
        if (line[0] == '[' && line[strlen(line) - 2] == ']') {
            line[strlen(line) - 2] = '\0';
            strcpy(section, line + 1);
            continue;
        }

        // set variables which obtained from ini file to global variable
        if (sscanf(line, "%127[^= ] = %127s", key, value_str) == 2) {
            if (set_char_var("output_dir", output_dir) == -1 ) return error_exit;
            if (set_char_var("input_file_path", input_file_path) == -1 ) return error_exit;
        }
    }
    close_ini(&file);
    log_message(INFO, "end configuration check");

    return success_exit;
}

int main(){

    double *data;
    int rows = 0;
    int columns = 0;

    scan_initial_values();

    FILE *fp = get_csv_pointer(input_file_path);
    if( fp == NULL){
        log_message(ERROR, "failed to read input file");
        release_csv(fp);
        return 1;
    }

    read_csv(fp, &data, &rows, &columns);

    printf("rows: %d\n", rows);
    printf("columns: %d\n", columns);
    int t_column_num = -1;
    int Plv_column_num = -1;
    int Vlv_column_num = -1;

    // search Plv and Vlv column num from data array
    t_column_num = get_column_num(fp, "t");
    Plv_column_num = get_column_num(fp, "Plv");
    Vlv_column_num = get_column_num(fp, "Vlv");
    if(t_column_num == -1 || Plv_column_num == -1 || Vlv_column_num == -1){
        log_message_with_line(ERROR, "failed to find parameters to search",__FILE__, __LINE__);
        return 1;
    }

    int row_t_ed = -1;
    int row_t_ej = -1;
    int row_t_es = -1;
    int row_t_fl = -1;
    double t_ed = 0.0;
    double t_ej = 0.0;
    double t_es = 0.0;
    double t_fl = 0.0;

    double Vlv_ed = 0.0;
    double Vlv_ej = 0.0;
    double Vlv_es = 0.0;
    double Vlv_fl = 0.0;
    double Plv_ed = 0.0;
    double Plv_ej = 0.0;
    double Plv_es = 0.0;
    double Plv_fl = 0.0;

    int cycle_flag = 0;

    // search the each points of single cardiac cycle
    for(int row = 1 ; row < rows - 1 ; row ++){
        if(cycle_flag == 0 && data[row*columns + Vlv_column_num] - data[(row+1)*columns + Vlv_column_num]== 0.0){
            row_t_ed = row + 1;
            t_ed = data[row*columns + t_column_num];
            Vlv_ed = data[row*columns + Vlv_column_num];
            Plv_ed = data[row*columns + Plv_column_num];
            cycle_flag = 1;
        }
        if(cycle_flag == 1 && data[row*columns + Vlv_column_num] - data[(row+1)*columns + Vlv_column_num] > 0.0){
            row_t_ej = row + 1;
            t_ej = data[row*columns + t_column_num];
            Vlv_ej = data[row*columns + Vlv_column_num];
            Plv_ej = data[row*columns + Plv_column_num];
            cycle_flag = 2;
        }
        if(cycle_flag == 2 && data[row*columns + Vlv_column_num] - data[(row+1)*columns + Vlv_column_num]== 0.0){
            row_t_es = row + 1;
            t_es = data[row*columns + t_column_num];
            Vlv_es = data[row*columns + Vlv_column_num];
            Plv_es = data[row*columns + Plv_column_num];
            cycle_flag = 3;
        }
        if(cycle_flag == 3 && data[row*columns + Vlv_column_num] - data[(row+1)*columns + Vlv_column_num] < 0.0){
            row_t_fl = row + 1;
            t_fl = data[row*columns + t_column_num];
            Vlv_fl = data[row*columns + Vlv_column_num];
            Plv_fl = data[row*columns + Plv_column_num];
            cycle_flag = 4;
        }
    }

    if(row_t_ed == -1 || row_t_ej == -1 || row_t_es == -1 || row_t_fl == -1){
        log_message_with_line(ERROR, "failed to find time for cardiac cycle", __FILE__, __LINE__);
    }

    double SV = Vlv_ed - Vlv_es;
    double CO = SV * 60.0;
    double EF = ( SV / Vlv_ed ) * 100.0;
    double IRT = t_fl  - t_es;
    double ICT = t_ej - t_ed;
    double ET = t_es - t_ej;
    double tau = IRT / (log(Plv_es) - log(Plv_fl));
    double Tei_index = ( (ICT + IRT) / ET ) * 100.0;

    FILE *output_file_path = NULL;
    if( create_csv(&output_file_path, output_dir) == 0 ){
        log_message_with_line(ERROR, "failed to output csv file", __FILE__, __LINE__);
        return 1;
    }

    print_char_csv(22, &output_file_path, "tED", "tEJ", "tES", "tFL", "Vlv_ED", "Vlv_EJ", "Vlv_ES", "Vlv_FL", "Plv_ED", "Plv_EJ", "Plv_ES", "Plv_FL", "SV", "CO", "ET", "IRT", "ICT", "EF", "ICT/ET", "IRT/ET", "tau", "Tei-index"); 
    print_var_csv(22, &output_file_path, t_ed, t_ej, t_es, t_fl, Vlv_ed, Vlv_ej, Vlv_es, Vlv_fl, Plv_ed, Plv_ej, Plv_es, Plv_fl, SV, CO, ET, IRT, ICT, EF, ICT/ET, IRT/ET, tau, Tei_index); 
    close_csv(&output_file_path);

    return 0;
}
```
