## Supplementary material for "A Theoretical Framework for the Hemodynamic Role of Sarcomere Length Dynamics During the Isovolumic Phase of the Left Ventricle": SourceCode: windkessel.c

```
#include<stdio.h>
#include "../../logging/logging.h"
#include "../../calculation/euler_calculation.h"

// logging global variables
extern const char *ERROR;
extern const char *WARN;
extern const char *INFO;

// time variables
extern double dt;
extern double Rout;

// circulation variables
extern double Plv;
extern double Vlv;
extern double Pa;
extern double Va;
extern double Rpv;
extern double Rlo;
extern double PE1;
extern double PE2;
extern double CA;

extern int q1_valve_count;
extern int qin_valve_count;

// local use variables
extern double q0; // [ml/ms]
extern double q1; // =dVlvdt [ml/ms]
extern double q2; // =dVadt [ml/ms]
extern double qin; // [ml/ms]
extern double qout; // [ml/ms]
extern double dPadt; // [mmHg/ms]

static void flow_calculation_with_valve(double *input, double *output, double *resistance, double *flow, int *valve_count){
    if( *input > *output ){ 
        *valve_count = *valve_count + 1;
        if( *valve_count > 2 ){
            *flow = ( *input - *output ) / *resistance;
        }else{
            *flow = 0.0;
        }
    }else{
        *flow = 0.0;
        *valve_count = 0;
    }
}

double calculate_flow(double input, double output, double resistance, int valve_count){
    double ret_flow;
    if( input > output) {
        valve_count = valve_count + 1;
        if( valve_count > 2){
            ret_flow = ( input - output ) / resistance;
        }else{
            ret_flow = 0.0;
        }
    }
    else{
        ret_flow = 0.0;
        valve_count = 0;
    }
    return ret_flow;
}

double calculate_lv_volume(double lv_pressure){

    int q1_valve_count_local = q1_valve_count;
    int qin_valve_count_local = qin_valve_count;

    double ret_Vlv;

    // q0 calculation
    double q0_local = calculate_flow(PE1, lv_pressure, Rpv, q1_valve_count_local);

    // qin calculation
    double qin_local = calculate_flow(lv_pressure, Pa, Rlo, qin_valve_count_local);

    double qout_local = (Pa - PE2) / Rout;
    double q1_local = q0_local - qin_local; // dVlvdt
    double q2_local = qin_local - qout_local; // dVadt

    // next step's Volume calculation
    double Va_local = solve(Va, q2_local, dt);
    ret_Vlv = solve(Vlv, q1_local, dt);

    // next step's aorta pressure calculation
    double dPadt_local = q2_local / CA;
    double Pa_local = solve(Pa, dPadt_local, dt);

    return ret_Vlv;
}

void calculate_circulation(double t){

    // q0 calculation
    flow_calculation_with_valve(&PE1, &Plv, &Rpv, &q0, &q1_valve_count);

    // qin calculation
    flow_calculation_with_valve(&Plv, &Pa, &Rlo, &qin, &qin_valve_count);

    // Compliance varying (dVlvdt, dVadt) calculation
    qout = (Pa - PE2) / Rout;
    q1 = q0 - qin; // dVlvdt
    q2 = qin - qout; // dVadt

    // if(qin_valve_count > 2){printf("%f %f %f %f %f %f %f %f %f %f\n", Plv, Vlv, Pa, Va, q0, q1, q2, qin, qout, dPadt);} // for test

    // next step's Volume calculation
    Va = solve(Va, q2, dt);
    Vlv = solve(Vlv, q1, dt);

    // next step's aorta pressure calculation
    dPadt = q2 / CA;
    Pa = solve(Pa, dPadt, dt);
}
```
