## Supplementary material for "A Theoretical Framework for the Hemodynamic Role of Sarcomere Length Dynamics During the Isovolumic Phase of the Left Ventricle": SourceCode: windkessel.h

```
#ifndef WINDKESSEL_H
#define WINDKESSEL_H

static void flow_calculation_with_valve();

double calculate_flow(double input_pressure, double output_pressure, double resistance, int valve_count);

double calculate_lv_volume(double lv_pressure);

void calculate_circulation();

#endif
```
