## Supplementary material for "A Theoretical Framework for the Hemodynamic Role of Sarcomere Length Dynamics During the Isovolumic Phase of the Left Ventricle": SourceCode: NL08_in_circulation.c

```
#include<stdio.h>
#include<math.h>
#include "../../logging/logging.h"
#include "../../calculation/euler_calculation.h"

extern double dt;

extern double L;
extern double Xp;
extern double Xw;
extern double dXpdt;
extern double dXwdt;
extern double TSCap;
extern double TSp;
extern double TSCaw;
extern double TSw;

extern double TSCaeff;
extern double TSCa;
extern double TS;
extern double Ca;

extern double dTSCadt;
extern double dTSCapdt;
extern double dTSCawdt;
extern double dTSpdt;
extern double dTSwdt;

extern double hp;
extern double hw;

extern double Itropn;
extern double Qpump;
extern double Qrel;
extern double Qsr;
extern double dCadt;

extern double g;
extern double gd;

extern double Kbs;
extern double Kps;

double calculate_active_force(double length){
    double Ap = 2.7E6;
    double Aw = 0.54E6;
    return Ap * (TSCap + TSp) * ( length - Xp ) + Aw * ( TSCaw + TSw ) * ( length -Xw );
}

double calculate_passive_force(double length){
    double L0 = 1.0;
    double Ke = 105000.0;
    double Le = 10.0;
    return Ke * pow((length - L0), 5.0) + Le * (length - L0);
}

double calculate_total_force(double Fb_local, double Fp_local){
   return Kbs * Fb_local + Kps * Fp_local;
}

void calculate_cellular_chem_state(double t_cycle){

    double gamma_factor = 28000.0;
    double hpr = 0.006;
    double hwr = 0.0001;
    double B = 0.5;

    double Za = 0.0023;
    double Yv = 1.5;
    double Yd = 0.0333;
    double Yc = 1.0;
    double Lc = 1.2;
    double Yvd = 1.5;
    double R = 15.0;
    double La = 1.15;
    double TSt = 0.023;
    double Yb = 0.1816E9;
    double Zb = 0.1397;
    double f = 0.0023;
    double Yp = 0.1397;
    double Zp = 0.2095;
    double Zr = 7.2626E9;
    double Yr = 0.1397;
    double Zq = 0.3724;
    double Yq = 0.2328;
    
    double Kp = 0.00015;
    double Km = 0.2E-3;
    double Qm = 0.0032;
    double t1 = 8.0;
    double Car = 0.1E-3;

    hp = L - Xp;
    hw = L - Xw;

    // calculate present step's differential values
    g = Za + Yv * ( 1.0 - exp( -gamma_factor * pow( hw - hwr , 2.0 ) ) );
    gd = Yd + Yc * pow( L - Lc, 2.0 ) + Yvd * ( 1.0 - exp( -gamma_factor * pow( hw - hwr , 2.0 ) ) );
    TSCaeff = exp( -R * pow( L - La, 2.0 ) ) * TSCa;
    TS = TSt - TSCa - TSCaw - TSCap - TSw - TSp;

    dTSCadt = Yb * TS * pow( Ca, 3.0 ) - Zb * TSCa + g * TSCaw - f * TSCaeff;
    dTSCapdt = Yp * TSCaw - Zp * TSCap + Zr * TSp * pow( Ca, 3.0 ) - Yr * TSCap;
    dTSCawdt = f * TSCaeff - g * TSCaw + Zp * TSCap - Yp * TSCaw;
    dTSpdt = Yr * TSCap - Zr * TSp * pow( Ca, 3.0 ) + Zq * TSw - Yq * TSp;
    dTSwdt = Yq * TSp - Zq * TSw - gd * TSw;

    dXpdt = B * ( hp - hpr );
    dXwdt = B * ( hw - hwr );

    // calculate next step's Ca, TSCap, TSp, TSCaw, TSw, Xp, Xw
    Itropn = 3 * ( dTSCadt + dTSCawdt + dTSCapdt );
	Qpump = Kp / (1 + pow( Km / Ca, 2.0 ));
	Qrel = Qm * pow( t_cycle / t1, 4.0 ) * exp( 4.0 * ( 1.0 - t_cycle / t1) ) + Kp / ( 1.0 + pow( Km / Car, 2.0 ) );
	Qsr =  Qrel - Qpump - Itropn;
	dCadt = Qrel - Qpump - Itropn;

	Ca = solve( Ca, dCadt, dt);
    TSCa = solve( TSCa, dTSCadt, dt );
    TSCap = solve( TSCap, dTSCapdt, dt );
    TSCaw = solve( TSCaw, dTSCawdt, dt );
    TSp = solve( TSp, dTSpdt, dt );
    TSw = solve( TSw, dTSwdt, dt );
    Xp = solve( Xp, dXpdt, dt );
    Xw = solve( Xw, dXwdt, dt );
}
```
