## Supplementary material for "A Theoretical Framework for the Hemodynamic Role of Sarcomere Length Dynamics During the Isovolumic Phase of the Left Ventricle": SourceCode: NL08_in_circulation.h

```
#ifndef NL08_IN_CIRCULATION_H
#define NL08_IN_CIRCULATION_H

double calculate_active_force(double length);

double calculate_passive_force(double length);

double calculate_total_force(double Fb_local, double Fp_local);

void calculate_cellular_chem_state(double t_cycle);

#endif
```
