## Supplementary material for "A Theoretical Framework for the Hemodynamic Role of Sarcomere Length Dynamics During the Isovolumic Phase of the Left Ventricle": SourceCode: canine_lvf_relation.c

```
#include<stdio.h>
#include<math.h>
#include "../../logging/logging.h"
#include "../../calculation/euler_calculation.h"

double get_length_from_canine_volume_and_force(double scaled_volume, double scaled_force){

    double k1 = -0.000211703721409215;
	double k2 = 0.00595529412713629;
	double k3 = -0.0478606732468164;
	double k4 = 0.134218450826179;
	double k5 = -0.0173588701924699;
	double k6 = -0.11430132914139;
	double k7 = 3.03980858064257;

	double p1 = 0.02;
	double p2 = 0.22;
	double p3 = -0.02;
	double p4 = 0.075;
	double p5 = 1.65;

    double half_scale = 0.5; // convert sarcomere length to half one

    double h1 = k1 * pow(p1, 6.0) * pow(scaled_volume, 6.0);
	double h2 = (k1 * 6.0 * p2 * scaled_force + k2) * pow(p1, 5.0) * pow(scaled_volume, 5.0);
	double h3 = (k1 * 15.0 * pow(p2, 2.0) * pow(scaled_force, 2.0) + k2 * 5.0 * p2 * scaled_force + k3) * pow(p1, 4.0) * pow(scaled_volume, 4.0);
	double h4 = (k1 * 20.0 * pow(p2, 3.0) * pow(scaled_force, 3.0) + k2 * 10.0 * pow(p2, 2.0) * pow(scaled_force, 2.0) + k3 * 4.0 * p2 * scaled_force + k4) * pow(p1, 3.0) * pow(scaled_volume, 3.0);
	double h5 = (k1 * 15.0 * pow(p2, 4.0) * pow(scaled_force, 4.0) + k2 * 10.0 * pow(p2, 3.0) * pow(scaled_force, 3.0) + k3 * 6.0 * pow(p2, 2.0) * pow(scaled_force, 2.0) + k4 * 3.0 * p2 * scaled_force + k5) * pow(p1, 2.0) * pow(scaled_volume, 2.0);
	double h6 = (-p3 + (k1 * 6.0 * pow(p2, 5.0) * pow(scaled_force, 5.0) + k2 * 5.0 * pow(p2, 4.0) * pow(scaled_force, 4.0) + k3 * 4.0 * pow(p2, 3.0) * pow(scaled_force, 3.0) + k4 * 3.0 * pow(p2, 2.0) * pow(scaled_force, 2.0) + k5 * 2.0 * p2 * scaled_force + k6) * p1) * scaled_volume;
	double h7 = k1 * pow(p2, 6.0) * pow(scaled_force, 6.0) + k2 * pow(p2, 5.0) * pow(scaled_force, 5.0) + k3 * pow(p2, 4.0) * pow(scaled_force, 4.0) + k4 * pow(p2, 3.0) * pow(scaled_force, 3.0) + k5 * pow(p2, 2.0) * pow(scaled_force, 2.0) + (k6 * p2 - p4) * scaled_force + k7;

    return ( ( h1 + h2 + h3 + h4 + h5 + h6+ h7 ) / p5 ) * half_scale;
}
```
