## Supplementary material for "A Theoretical Framework for the Hemodynamic Role of Sarcomere Length Dynamics During the Isovolumic Phase of the Left Ventricle": SourceCode: laplace.c

```
#include<stdio.h>
#include<math.h>
#include "../../logging/logging.h"
#include "../../calculation/euler_calculation.h"
#include "../../calculation/numeric_conversion/scaling_force.h"

extern int model_flag;
extern int mode_change_flag;

double calculate_pressure(double force, double radius, double wall_thickness){

    double Ks = get_force_scale_param(model_flag, mode_change_flag);
    double Ku = 7.50064; // [mN/mm^2]->[mmHg] 

    return ( 2.0 * Ks *  force * wall_thickness * Ku) / radius ;

}
```
