## Supplementary material for "A Theoretical Framework for the Hemodynamic Role of Sarcomere Length Dynamics During the Isovolumic Phase of the Left Ventricle": SourceCode: length_by_radius.c

```
#include <stdio.h>
#include <math.h>
#include "../../logging/logging.h"

double calculate_length_from_radius(double lv_radius){
    const double KL = 0.01685393258;
    const double Lb = 0.670325;
    return ( KL * lv_radius + Lb);
}

double calculate_length_from_radius_in_isovolume(double lv_radius){
    const double KL = 0.01651083069876305;
    const double Lb = 0.6767141527782496;

    return (KL * lv_radius + Lb);
}
```
