## Supplementary material for "A Theoretical Framework for the Hemodynamic Role of Sarcomere Length Dynamics During the Isovolumic Phase of the Left Ventricle": SourceCode: length_by_volume.c

```
#include<stdio.h>
#include<math.h>
#include "../../logging/logging.h"
#include "../../calculation/euler_calculation.h"
#include "./rad_by_sphere_vol.h"
#include "./length_by_radius.h"

extern int mode_change_flag;

double calculate_length_from_volume(double volume){
    double radius = calculate_radius_from_volume(volume);
    double length;
    if(mode_change_flag == 0){
        length = calculate_length_from_radius(radius);
    }else{
        length = calculate_length_from_radius_in_isovolume(radius);
    } 
    return length;
}
```
