## Supplementary material for "A Theoretical Framework for the Hemodynamic Role of Sarcomere Length Dynamics During the Isovolumic Phase of the Left Ventricle": SourceCode: length_by_volume_force.c

```
#include<stdio.h>
#include<math.h>
#include "../../logging/logging.h"
#include "../../calculation/euler_calculation.h"
#include "../../calculation/numeric_conversion/conversion_to_canine_param.h"
#include "../../calculation/numeric_conversion/scaling_force.h"
#include "./canine_lvf_relation.h"

extern int model_flag;
extern int mode_change_flag;

double calculate_length_from_volume_force(double volume, double force){

    double Ks = get_force_scale_param(model_flag, mode_change_flag);
	double scaled_force = convert_to_canine_lv_Fext(Ks, force);
    double scaled_volume = convert_to_canine_lv_volume(volume);

    return get_length_from_canine_volume_and_force(scaled_volume, scaled_force);
}
```
