## Supplementary material for "A Theoretical Framework for the Hemodynamic Role of Sarcomere Length Dynamics During the Isovolumic Phase of the Left Ventricle": SourceCode: rad_by_sphere_vol.c

```
#include<stdio.h>
#include<math.h>
#include "../../logging/logging.h"

#ifndef M_PI
#define M_PI 3.14159265358979323846
#endif

double calculate_radius_from_volume(double volume){
    // internal use variables
    double KV = 8.90633;
    double KR = 10.80977;
    
    // fitting assumed Sphere
    return KR * pow( (  3.0  / ( 4.0 * M_PI ) * volume ), 1.0 / 3.0 ) - KV;

    //return 10.0 *(KR * volume + KV);

}
```
