## Supplementary material for "A Theoretical Framework for the Hemodynamic Role of Sarcomere Length Dynamics During the Isovolumic Phase of the Left Ventricle": SourceCode: wall_thickness_by_rad.c

```
#include<stdio.h>
#include<math.h>
#include "../../logging/logging.h"
#include "../../calculation/euler_calculation.h"

double calculate_wall_thickness(double radius){
    // internal use variables
    double hlvED = 9.8; // [mm]
    double hlvES = 17.2; // [mm]
    double RlvED = 25.5; // [mm]
    double RlvES = 16.6; // [mm]
    
    return (( hlvES - hlvED ) / ( RlvES - RlvED ) ) * (radius - RlvED) + hlvED;
    
}
```
