## Supplementary material for "A Theoretical Framework for the Hemodynamic Role of Sarcomere Length Dynamics During the Isovolumic Phase of the Left Ventricle": SourceCode: plot_utility.c

```
#include <stdio.h>
#include <math.h>
#include <time.h>
#include<stdlib.h>
#include <unistd.h>
#include "../logging/logging.h"
#include<stdarg.h>

// time variables
extern double duration;
extern double dt;
extern double cycle;

// logging global variables
extern const char *ERROR;
extern const char *WARN;
extern const char *INFO;

// plot settings
extern const int realtime_plot;
extern const int every_fixed_times;

int open_plot_process(FILE **gp){
    *gp = popen("gnuplot -persist", "w");
    
    if(*gp == NULL){
        log_message(ERROR, "failed to open gnuplot");
        return 0;
    }
    fprintf(*gp, "set multiplot\n");
    fprintf(*gp, "set termopt enhanced\n");
    
    log_message(INFO, "opened gnuplot");
    return 1;
}

void set_xaxis(FILE **gp, double lowest, double highest, char *label_name){
    fprintf(*gp, "set xrange [%f:%f]\n", lowest, highest);
    fprintf(*gp, "set xlabel \"%s\"\n", label_name);
}

void set_yaxis(FILE **gp, double lowest, double highest, char *label_name){
    fprintf(*gp, "set yrange [%f:%f]\n", lowest, highest);
    fprintf(*gp, "set ylabel \"%s\"\n", label_name);
}

void set_xyaxis(FILE **gp, double x_lowest, double x_highest, char *x_label_name, double y_lowest, double y_highest, char *y_label_name){
    fprintf(*gp, "set xrange [%f:%f]\n", x_lowest, x_highest);
    fprintf(*gp, "set xlabel \"%s\"\n", x_label_name);
    fprintf(*gp, "set yrange [%f:%f]\n", y_lowest, y_highest);
    fprintf(*gp, "set ylabel \"%s\"\n", y_label_name);
}

int initialize_plot_process(FILE **gp, double x_lowest, double x_highest, char *x_label_name, double y_lowest, double y_highest, char *y_label_name){
    if(realtime_plot != 1){
        log_message("INFO", "disabled realtime plot mode");
        return 0;
    }
    
    *gp = popen("gnuplot -persist", "w");
    
    if(*gp == NULL){
        log_message(ERROR, "failed to open gnuplot");
        return 0;
    }
    fprintf(*gp, "set multiplot\n");
    fprintf(*gp, "set termopt enhanced\n");
    
    log_message(INFO, "opened gnuplot");

    fprintf(*gp, "set xrange [%f:%f]\n", x_lowest, x_highest);
    fprintf(*gp, "set xlabel \"%s\"\n", x_label_name);
    fprintf(*gp, "set yrange [%f:%f]\n", y_lowest, y_highest);
    fprintf(*gp, "set ylabel \"%s\"\n", y_label_name);
    log_message(INFO, "configured xyaxis settings");
    return 1;
}

void open_plot_pipe(FILE **gp, char *point_type){
    fprintf(*gp, "plot '-' w p pointtype %s\n", point_type);
}

void plot_with_point(FILE **gp, double x_value, double y_value){
    if(realtime_plot == 1){
        fprintf(*gp, "%f\t%f\n", x_value, y_value);
    }
}

void plot_with_multipoints(FILE **gp, double x_value, int y_num, ...){
    if(realtime_plot == 1){
        va_list args;
        va_start(args, y_num);

        for(int i = 0 ; i < y_num ; i++){
            double y_value = va_arg(args, double);
            fprintf(*gp, "%f\t%f\n", x_value, y_value);
        }
        va_end(args);
    }
    
}


void plot_with_point_every_fixed_time(FILE **gp, double t_cycle, double x_value, double y_value ){
    if(realtime_plot == 1){
        if(fmod(t_cycle, every_fixed_times) < dt){
            fprintf(*gp, "%f\t%f\n", x_value, y_value);
            fflush(*gp);
        }
    }
}

void plot_with_multipoints_every_fixed_time(FILE **gp, double t_cycle, double x_value, int y_num, ...){
    if(realtime_plot == 1){
        if(fmod(t_cycle, every_fixed_times) < dt){
            va_list args;
            va_start(args, y_num);

            for(int i = 0 ; i < y_num ; i++){
                double y_value = va_arg(args, double);
                fprintf(*gp, "%f\t%f\n", x_value, y_value);
            }
            va_end(args);
            fflush(*gp);
        }
    }
}

void close_plot_pipe(FILE **gp){
    fprintf(*gp, "e\n");
}

void flush_plot(FILE **gp){
    fflush(*gp);
}

void clear_plot(FILE **gp){
    fprintf(*gp, "clear\n");
}

void close_plot_process(FILE **gp){
    fprintf(*gp, "exit\n");
    pclose(*gp);
    log_message(INFO, "closed gnuplot");
}

void define_pointtype_clear_prev_cycle(FILE **gp, char *point_type, double t_cycle){
    if(realtime_plot == 1 && t_cycle < dt){
        fprintf(*gp, "clear\n");
        fprintf(*gp, "plot '-' w p pointtype %s\n", point_type);
    }
}

void output_buffers_at_cycle_end(FILE **gp, double t_cycle){
    if(realtime_plot == 1 && (t_cycle >= (cycle - dt))){
        fprintf(*gp, "e\n");
    }
}

void terminate_plot_process(FILE **gp){
    if(realtime_plot == 1){
        fprintf(*gp, "exit\n");
        pclose(*gp);
        log_message(INFO, "closed gnuplot");
    }
}
```
