## Supplementary material for "A Theoretical Framework for the Hemodynamic Role of Sarcomere Length Dynamics During the Isovolumic Phase of the Left Ventricle": SourceCode: plot_utility.h

```
#ifndef PLOT_UTILITY_H
#define PLOT_UTILITY_H

int open_plot_process(FILE **gp);

void set_xaxis(FILE **gp, double lowest, double highest, char *label_name);

void set_yaxis(FILE **gp, double lowest, double highest, char *label_name);

void set_xyaxis(FILE **gp, double x_lowest, double x_highest, char *x_label_name, double y_lowest, double y_highest, char *y_label_name);

int initialize_plot_process(FILE **gp, double x_lowest, double x_highest, char *x_label_name, double y_lowest, double y_highest, char *y_label_name);

void open_plot_pipe(FILE **gp, char *point_type);

void plot_with_point(FILE **gp, double x_value, double y_value);

void plot_with_multipoints(FILE **gp, double x_value, int y_num, ...);

void plot_with_point_every_fixed_time(FILE **gp, double t_cycle, double x_value, double y_value );

void plot_with_multipoints_every_fixed_time(FILE **gp, double t_cycle, double x_value, int y_num, ...);

void close_plot_pipe(FILE **gp);

void flush_plot(FILE **gp);

void clear_plot(FILE **gp);

void close_plot_process(FILE **gp);

void define_pointtype_clear_prev_cycle(FILE **gp, char *point_type, double t_cycle);

void output_buffers_at_cycle_end(FILE **gp, double t_cycle);

void terminate_plot_process(FILE **gp);

#endif
```
