## Supplementary material for "A Theoretical Framework for the Hemodynamic Role of Sarcomere Length Dynamics During the Isovolumic Phase of the Left Ventricle": ModelDescription

### 1 Model equations

#### 1.1 Circulation model

$$q_0 = \begin{cases} (P_{E1} - P_{lv})/R_{pv} & P_{E1} > P_{lv} \\ 0 & otherwise \end{cases} \tag{1}$$

$$q_{in} = \begin{cases} (P_{lv} - P_a)/R_{lo} & P_{lv} > P_a \\ 0 & otherwise \end{cases} \tag{2}$$

$$q_{out} = (P_a - P_{E2})/R_{out} \tag{3}$$

$$P_a = V_a/C_a \tag{4}$$

$$\frac{dV_a}{dt} = q_{in} - q_{out} \tag{5}$$

$$\frac{dV_{lv}}{dt} = q_0 - q_{in} \tag{6}$$

#### 1.2 Ventricular cell contraction force model

##### 1.2.1 Chemical reaction component

$$\frac{d[TSCa_3]}{dt} = Y_b[TS][Ca^{2+}]^3 - Z_b[TSCa_3] + g[TSCa_3^{\sim}] - f[TSCa_3]_{\text{eff}} \tag{7}$$

$$\frac{d[TSCa_3^{\sim}]}{dt} = f[TSCa_3]_{\text{eff}} - g[TSCa_3^{\sim}] + Z_p[TSCa_3^*] - Y_p[TSCa_3^{\sim}] \tag{8}$$

$$\frac{d[TSCa_3^*]}{dt} = Y_p[TSCa_3^{\sim}] - Z_p[TSCa_3^*] + Z_r[TS^*][Ca^{2+}]^3 - Y_r[TSCa_3^*] \tag{9}$$

$$\frac{d[TS^*]}{dt} = Y_r[TSCa_3^*] - Z_r[TS^*][Ca^{2+}]^3 + Z_q[TS^{\sim}] - Y_q[TS^*] \tag{10}$$

$$\frac{d[TS^{\sim}]}{dt} = Y_q[TS^*] - Z_q[TS^{\sim}] - g_d[TS^{\sim}] \tag{11}$$

$$[TS] = [TS]_t - [TSCa_3] - [TSCa_3^{\sim}] - [TSCa_3^*] - [TS^*] - [TS^{\sim}] \quad (12)$$

$$[TSCa_3]_{\text{eff}} = e^{-R(L-L_a)^2}[TSCa_3] \quad (13)$$

$$g = Z_a + Y_v\left(1 - e^{-\gamma_m(h_w-h_{wr})^2}\right) \quad (14)$$

$$g_d = Y_d + Y_c(L-L_c)^2 + Y_{vd}\left(1 - e^{-\gamma_m(h_w-h_{wr})^2}\right) \quad (15)$$

$$\gamma_m = \begin{cases} \gamma\frac{1}{K_\gamma} & \frac{dX_w}{dt} > 0 \\ \gamma & otherwise \end{cases} \quad (16)$$

##### 1.2.2 Calcium dynamics of chemical reaction component

$$Q_{rel} = Q_m\left(\frac{t}{t_1}\right)^4 e^{4\left(1-\frac{t}{t_1}\right)} + Q_{pump\_rest} \quad (17)$$

$$Q_{pump} = K_p\frac{1}{1+\left(\frac{K_m}{[Ca^{2+}]}\right)^2} \quad (18)$$

$$\frac{d[Ca^{2+}]}{dt} = Q_{rel} - Q_{pump} - I_{troponin} \quad (19)$$

$$I_{troponin} = 3\left(\frac{d[TSCa_3]}{dt} + \frac{d[TSCa_3^{\sim}]}{dt} + \frac{d[TSCa_3^*]}{dt}\right) \quad (20)$$

##### 1.2.3 Mechanical component

$$h_w = L - X_w \quad (21)$$

$$h_p = L - X_p \quad (22)$$

$$\frac{dX_w}{dt} = B(h_w - h_{wr}) \quad (23)$$

$$\frac{dX_p}{dt} = B(h_p - h_{pr}) \quad (24)$$

$$F_p = K_e(L-L_0)^5 + L_e(L-L_0) \quad (25)$$

$$F_b = A_w([TSCa_3^{\sim}] + [TS^{\sim}])h_w + A_p([TSCa_3^*] + [TS^*])h_p \quad (26)$$

$$F = K_{bs}F_b + K_{ps}F_p \quad (27)$$

### 1.3 Left Ventricular pressure and cell contraction force relation model

#### 1.3.1 Common equations for the two models

$$P_{lv} = \frac{2K_u F_{ext} h_{lv}}{R_{lv}} \tag{28}$$

$$F_{ext} = K_S F \tag{29}$$

$$R_{lv} = K_R(\frac{3}{4\pi}V_{lv})^{\frac{1}{3}} - K_V \tag{30}$$

$$h_{lv} = \frac{R_{lv} - R_{lvED}}{R_{lvES} - R_{lvED}}(h_{lvES} - h_{lvED}) + h_{lvED} \tag{31}$$

#### 1.3.2 Sarcomere length of volume-based length model (VL)

$$L = K_L R_{lv} + L_b \tag{32}$$

#### 1.3.3 Sarcomere length of volume-force-coupled length model (VFL)

$$L = \frac{h_1 + h_2 + h_3 + h_4 + h_5 + h_6 + h_7}{2P_5} \tag{33}$$

$$V_{lv,canine} = K_{va}V_{lv} + K_{vb} \tag{34}$$

$$F_{ext,canine} = K_{fa}F + K_{fb} \tag{35}$$

$$h_1 = K_1 P_1^6 V_{lv,canine}^6 \tag{36}$$

$$h_2 = (6K_1 P_2 F_{ext,canine} + K_2)P_1^5 V_{lv,canine}^5 \tag{37}$$

$$h_3 = (15K_1 P_2^2 F_{ext,canine}^2 + 5K_2 P_2 F_{ext,canine} + K_3)P_1^4 V_{lv,canine}^4 \tag{38}$$

$$\begin{aligned} h_4 = (&20K_1 P_2^3 F_{ext,canine}^3 + 10K_2 P_2^2 F_{ext,canine}^2 \\ &+4K_3 P_2 F_{ext,canine} + K_4)P_1^3 V_{lv,canine}^3 \end{aligned} \tag{39}$$

$$h_5 = (15K_1P_2^4F_{ext,canine}^4 + 10K_2P_2^3F_{ext,canine}^3 + 6K_3P_2^2F_{ext,canine}^2 \\ +3K_4P_2F_{ext,canine} + K_5)P_1^2V_{lv,canine}^2 \tag{40}$$

$$h_6 = ((6K_1P_2^5F_{ext,canine}^5 + 5K_2P_2^4F_{ext,canine}^4 + 4K_3P_2^3F_{ext,canine}^3 \\ +3K_4P_2^2F_{ext,canine}^2 + K_5P_2F_{ext,canine} + K_6)P_1 - P_3)V_{lv,canine} \tag{41}$$

$$h_7 = K_1P_2^6F_{ext,canine}^6 + K_2P_2^5F_{ext,canine}^5 + K_3P_2^4F_{ext,canine}^4 + K_4P_2^3F_{ext,canine}^3 \\ +K_5P_2^2F_{ext,canine}^2 + (K_6P_2 - P_4)F_{ext,canine} + K_7 \tag{42}$$

### 2 Parameter values and model schematics

Table 1: **Descriptions of the circulation model variable**

| parameter | description | value | unit |
|---|---|---|---|
| $P_{E1}$ | Pulmonary venous pressure | 10.0 – 14.0 | [mmHg] |
| $P_{lv}$ | LV pressure | variable | [mmHg] |
| $P_a$ | Aortic pressure | variable | [mmHg] |
| $P_{E2}$ | Peripheral pressure | 15.0 | [mmHg] |
| $R_{pv}$ | Pulmonary venous resistance | 27.0 | [mmHg · ms/mL] |
| $R_{lo}$ | Aortic resistance | 1.0 | [mmHg · ms/mL] |
| $R_{out}$ | Peripheral resistance | 900 – 1200 | [mmHg · ms/mL] |
| $C_{lv}$ | LV compliance | variable | [mL/mmHg] |
| $C_a$ | Aortic compliance | 1.5 | [mL/mmHg] |

Table 2: **Parameters used for chemical reaction component of ventricular cell contraction force model**

| parameter | value | unit |
|---|---|---|
| $Y_b$ | $0.1816 \times 10^9$ | [$\mathrm{mM}^3$/ms] |
| $Y_p$ | 0.1397 | [/ms] |
| $Y_r$ | 0.1397 | [/ms] |
| $Y_q$ | 0.2328 | [/ms] |
| $Z_b$ | 0.1397 | [/ms] |
| $Z_p$ | 0.2095 | [/ms] |
| $Z_r$ | $7.2626 \times 10^9$ | [$\mathrm{mM}^3$/ms] |
| $Z_q$ | 0.3724 | [/ms] |
| $f$ | 0.0023 | [/ms] |
| $Z_a$ | 0.0023 | [/ms] |
| $Y_v$ | 1.5 | [/ms] |
| $Y_d$ | 0.0333 | [/ms] |
| $Y_c$ | 1.0 | [ms/$\mu\mathrm{m}^2$] |
| $Y_{vd}$ | 1.5 | [/ms] |
| $Q_m$ | 3.2 | [$\mu$M/ms] |
| $t_1$ | 8.0 | [ms] |
| $Q_{pump\_rest}$ | 0.03 | [$\mu$M/ms] |
| $K_p$ | 0.15 | [$\mu$M/ms] |
| $K_m$ | 0.2 | [$\mu$M] |

Table 3: **Parameters used for mechanical component of ventricular cell contraction force model**

| parameter | value | unit |
|---|---|---|
| $B$ | 0.5 | [/ms] |
| $h_{wr}$ | 0.0001 | [$\mu$m] |
| $h_{pr}$ | 0.006 | [$\mu$m] |
| $K_\gamma$ | 85.3 | - |
| $\gamma$ | 28000 | [/$\mu\mathrm{m}^2$] |
| $L_a$ | 1.15 | [$\mu$m] |
| $L_c$ | 1.2 | [$\mu$m] |
| $R$ | 15.0 | [/$\mu\mathrm{m}^2$] |
| $K_e$ | 105000.0 | [$\mathrm{mN/mM^2/\mu m^5}$] |
| $L_e$ | 10.0 | [$\mathrm{mN/mM^2/\mu m}$] |
| $A_w$ | $0.54 \times 10^6$ | [$\mathrm{mN/mm^2/\mu m/mM}$] |
| $A_p$ | $2.7 \times 10^6$ | [$\mathrm{mN/mm^2/\mu m/mM}$] |
| $L_0$ | 1.0 | [$\mu$m] |
| $K_{bs}$ | 2.0 | - |
| $K_{ps}$ | 0.25 | - |

Table 4: **Parameters used for Left Ventricular pressure and the cell contraction force relation model**

| parameter | value | unit |
|---|---|---|
| $K_u$ | 7.50064 | [mmHg/mN/mm$^2$] |
| $K_S$(VL) | 3.79 | - |
| $K_S$(VFL) | 3.6195 | - |
| $K_R$ | 10.80977 | [mm/mL] |
| $K_V$ | 8.90633 | [mm] |
| $K_{R,canine}$ | 3.4271728683883786 | [mm/mL] |
| $K_{V,canine}$ | -9.72901684720539 | [mm] |
| $K_L$ | 0.01685393258 | [$\mu$m/mm] |
| $L_b$ | 0.670325 | [$\mu$m] |
| $h_{lvED}$ | 9.8 | [mm] |
| $h_{lvES}$ | 17.2 | [mm] |
| $R_{lvED}$ | 25.5 | [mm] |
| $R_{lvES}$ | 16.6 | [mm] |
| $h_{lvED,canine}$ | 10.0 | [mm] |
| $h_{lvES,canine}$ | 14.0 | [mm] |
| $R_{lvED,canine}$ | 17.0 | [mm] |
| $R_{lvES,canine}$ | 15.5 | [mm] |
| $K_{va}$ | 0.2317073170733968 | - |
| $K_{vb}$ | 6.32926829266959 | [mL] |
| $K_{fa}$ | 1.0 | - |
| $K_{fb}$ | 1.5 | [mN/mm$^2$] |
| $K_1$ | -0.000211703721409215 | - |
| $K_2$ | 0.00595529412713629 | - |
| $K_3$ | -0.0478606732468164 | - |
| $K_4$ | 0.134218450826179 | - |
| $K_5$ | -0.0173588701924699 | - |
| $K_6$ | -0.11430132914139 | - |
| $K_7$ | 3.03980858064257 | - |
| $P_1$ | 0.02 | - |
| $P_2$ | 0.22 | - |
| $P_3$ | -0.02 | - |
| $P_4$ | 0.075 | - |
| $P_5$ | 1.65 | - |

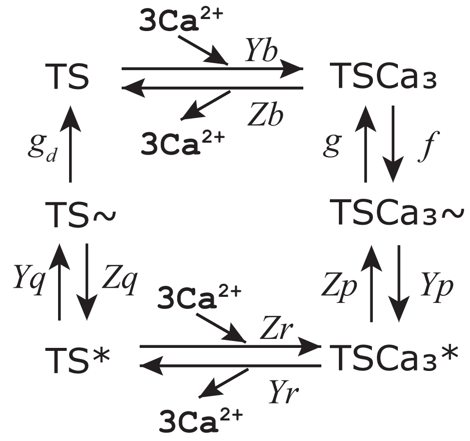


Figure 1: **NL08 chemical state transition model.** Reproduced from Kato et al. 2024, under the terms of the Creative Common Attribution License. [$TSCa_3^{\sim}$], [$TSCa_3^{*}$], [$TS^{\sim}$], and [$TS^{*}$] represent the concentrations of the troponin system binding the crossbridges with the weak($\sim$) and power($*$) states.
